# Recent hybrids recapitulate ancient hybrid outcomes

**DOI:** 10.1101/769901

**Authors:** Samridhi Chaturvedi, Lauren K. Lucas, C. Alex Buerkle, James A. Fordyce, Matthew L. Forister, Chris C. Nice, Zachariah Gompert

## Abstract

Genomic outcomes of hybridization depend on selection and recombination in hybrids. Whether these processes have similar effects on hybrid genome composition in contemporary hybrid zones versus ancient hybrid lineages is unknown. Here we show that patterns of introgression in a contemporary hybrid zone in *Lycaeides* butterflies predict patterns of ancestry in geographically adjacent, older hybrid populations. We find a particularly striking lack of ancestry from one of the hybridizing taxa, *Lycaeides melissa*, on the Z chromosome in both the old and contemporary hybrids. The same pattern of reduced *L. melissa* ancestry on the Z chromosome is seen in two other ancient hybrid lineages. More generally, we find that patterns of ancestry in old or ancient hybrids are remarkably predictable from contemporary hybrids, which suggests selection and recombination affect hybrid genomes in a similar way across disparate time scales and during distinct stages of speciation and species breakdown.

---

Numerous species hybridize [1, 2] or show genomic evidence of ancient admixture [3, 4, 5]. Consequently, many organisms have genomes that are mosaics of chromosomal segments with ancestry from different lineages or species (e.g., [6, 7, 8, 9, 10, 11]). In hybrid zones, where gene flow and hybridization are ongoing, parental chromosomal segments are repeatedly introduced, and hybrids often vary substantially in genome composition [12, 13, 14]. In contrast, when gene flow and hybridization cease, as occurs during hybrid speciation or if one of the hybridizing taxa goes extinct, genome stabilization can occur whereby recombination causes ancestry segment size to decay and ancestry segments fix by drift or selection in the nascent hybrid lineage [7, 15, 16].

In both cases, the genomic mosaic of ancestry segments is shaped by recombination and selection, including selection against developmental incompatibilities and selection for locally adaptive ancestry combinations [12, 14]. In contemporary hybrid zones with ongoing gene flow, selection results in differential or restricted introgression of some ancestry segments (e.g., barrier loci) [17, 18, 19, 20, 21], whereas selection causes shifts in the frequency of ancestry segments (including fixation) in old or ancient, stabilized or partially stabilized hybrid populations [22, 10, 16]. Recombination interacts with selection to shape hybrid genomes [23, 24, 16, 25]. In regions of higher recombination, neutral or adaptive foreign alleles can more readily disassociate from deleterious alleles allowing them to introgress, whereas in regions of low recombination selection can prevent large blocks of ancestry from introgressing. Changes in genome composition and patterns of linkage disequilibrium (mediated by recombination) during the transition from hybrid zone to stabilized hybrid lineage could alter selection on ancestry segments. Likewise, the primary sources of selection could change from, for example, selection against developmental incompatibilities to selection for novel allele combinations that enhance fitness and persistence in a new environment.

Unfortunately, comparisons of genome composition in contemporary versus old or ancient hybrids (i.e., those showing progress towards genome stabilization) are mostly lacking, especially for natural hybrids (for lab hybrids, see [22, 16]). Consistency between patterns of introgression in contemporary hybrid zones and the genomic mosaic of ancestry in ancient hybrids would suggest a major role for selection in determining hybrid genome composition, and would establish connections between early and late stages of speciation, especially speciation with gene flow and hybrid speciation. For example, consistent patterns would suggest that the same genes or gene regions that prevent the fusion of hybridizing species also experience selection during the origin of hybrid lineages or species. Such consistency could allow genes or genetic regions of general importance for adaptation and speciation to be identified (e.g., genes with environment-independent effects on fitness) and aid in interpreting patterns of ancestry in cases where ancient admixture occurred but where contemporary hybrids are lacking (e.g., *Homo sapiens* × *H. neanderthalensis* [8]). In contrast, a lack of consistency might imply that genomic outcomes of hybridization are highly context dependent, and thus difficult to predict (e.g., [26]).

Here, we take advantage of natural hybridization in North American *Lycaeides* butterflies to test the consistency of genome composition between a contemporary hybrid zone and multiple, old or ancient hybrid lineages that have progressed towards genome stabilization. In North America, *Lycaeides* consists of a complex of four nominal species of small blue butterflies and numerous partially stabilized (i.e., old or ancient) hybrid populations or lineages [6, 27, 28]. Partially stabilized hybrid populations in the central Rocky mountains and Jackson Hole (hereafter Jackson Hole *Lycaeides*) arose following hybridization between *L. idas* and *L. melissa* within the past 14,000 years (we refer to these as ancient hybrids hereafter) [29]. Like *L. idas* (and unlike *L. melissa*), these Jackson Hole *Lycaeides* exhibit obligate diapause and are univoltine; they also use the same larval host plant (*Astragalus miser*) as nearby *L. idas* populations [29, 30, 31]. In contrast, partially stabilized hybrid lineages in the Sierra Nevada and Warner mountains of western North America occupy extreme alpine habitats not used by the parental species and exhibit novel, transgressive phenotypes, such as strong preference for an alpine endemic host plant (*A. whitneyi*) and a lack of egg adhesion to the host substrate [6, 32]. We recently found evidence suggestive of a narrow (1–2 kilometers), contemporary hybrid zone between *L. melissa* and Jackson Hole *Lycaeides* near the town of Dubois, Wyoming at the edge of the Rocky mountains [28]. Butterflies in the putative hybrid zone use feral (i.e., naturalized), roadside alfalfa (*Medicago sativa*) as their host plant (similar to some *L. melissa* populations), and are found in close proximity (along the road) to this nonnative plant.

In the current study, we first verify that the Dubois population constitutes a contemporary hybrid zone between *L. melissa* and Jackson Hole *Lycaeides*; Jackson Hole *Lycaeides* is an ancient, partially stabilized hybrid lineage derived from *L. melissa* and *L. idas* (see [29, 30] and the Results below). We then use a mixture of genotyping-by-sequencing (GBS) and whole-genome sequence data to compare patterns of introgression in this hybrid zone to the genomic mosaic of ancestry in Jackson Hole *Lycaeides*, and to additional ancient hybrid lineages in the Sierra Nevada and Warner mountains, and thereby quantify the consistency of hybrid genome composition across these cases. We expect less consistency with the Sierra Nevada and Warner mountains lineages as they differ in their ecology and in their taxonomic origin; nonetheless, these lineages represent genomic outcomes of hybridization between *Lycaeides* and are useful for understanding the general aspects of what transpires from initial hybridization, through isolation and stabilization. We thus use the contemporary Dubois hybrid zone as a window on the evolutionary process and ask whether similar or very different processes operated during the origin and establishment of multiple, ancient hybrid lineages.

## Results

### Evidence of a contemporary hybrid zone in Dubois

Patterns of genetic variation across 39,193 SNPs and 23 populations (*N* = 835 butterflies) show that the Dubois, Wyoming population constitutes a contemporary hybrid zone between *L. melissa* and the ancient Jackson Hole hybrid lineage (Table S1, Figs. 1, 2). Admixture proportions from `entropy` (ver. 1.2; [28]) and a principal component analysis (PCA) of genetic variation show that Jackson Hole *Lycaeides* and the Dubois population are genetically intermediate between *L. idas* and *L. melissa* (Figs. 2B, S1, S2). Jackson Hole *Lycaeides* form a relatively tight cluster in PCA space (especially individuals within populations) and show similar admixture proportions, consistent with past admixture but little ongoing gene flow from *L. idas* or *L. melissa* (Fig. 2B,C) (i.e., consistent with ancient rather than contemporary hybridization; see [29, 30], Figs. S3, S4, and ‘Patterns of isolation-by-distance and taxon’ in the on-line supplemental material [OSM] for additional details regarding geographic patterns of genetic differentiation in Jackson Hole *Lycaeides*). In contrast, butterflies from the Dubois population span the entire genomic gradient from *L. melissa* to Jackson Hole *Lycaeides* (Fig. 2B,D). Thus, this single population, which occupies ~1–2 kilometers (kms) along roadside alfalfa, exhibits greater variation in genome composition than the entirety of Jackson Hole *Lycaeides*, with a range of more than 10,000 km^2^ [29]. Also consistent with ongoing hybridization in the Dubois population, these butterflies exhibit elevated coupling (positive) linkage disequilibrium (Figs. S5, S6) and intermediate allele frequencies (Figs. S7, S8, S9; see Figs. S10, S11 and S12 for analogous results with Jackson Hole *Lycaeides*) at ancestry informative SNPs (i.e., SNPs with an allele frequency difference of 0.3 or more between *L. idas* and *L. melissa*). Lastly, a discriminant analysis of genetic PCs confirmed that *L. melissa* and Jackson Hole *Lycaedies*, but not *L. idas*, occur in the Dubois hybrid zone (Fig. S13). Taken together, these results demonstrate ongoing hybridization in Dubois between *L. melissa* and nearby Jackson Hole *Lycaeides*, which are themselves a product of ancient hybridization between *L. idas* and *L. melissa* [29].

**Figure 1:**
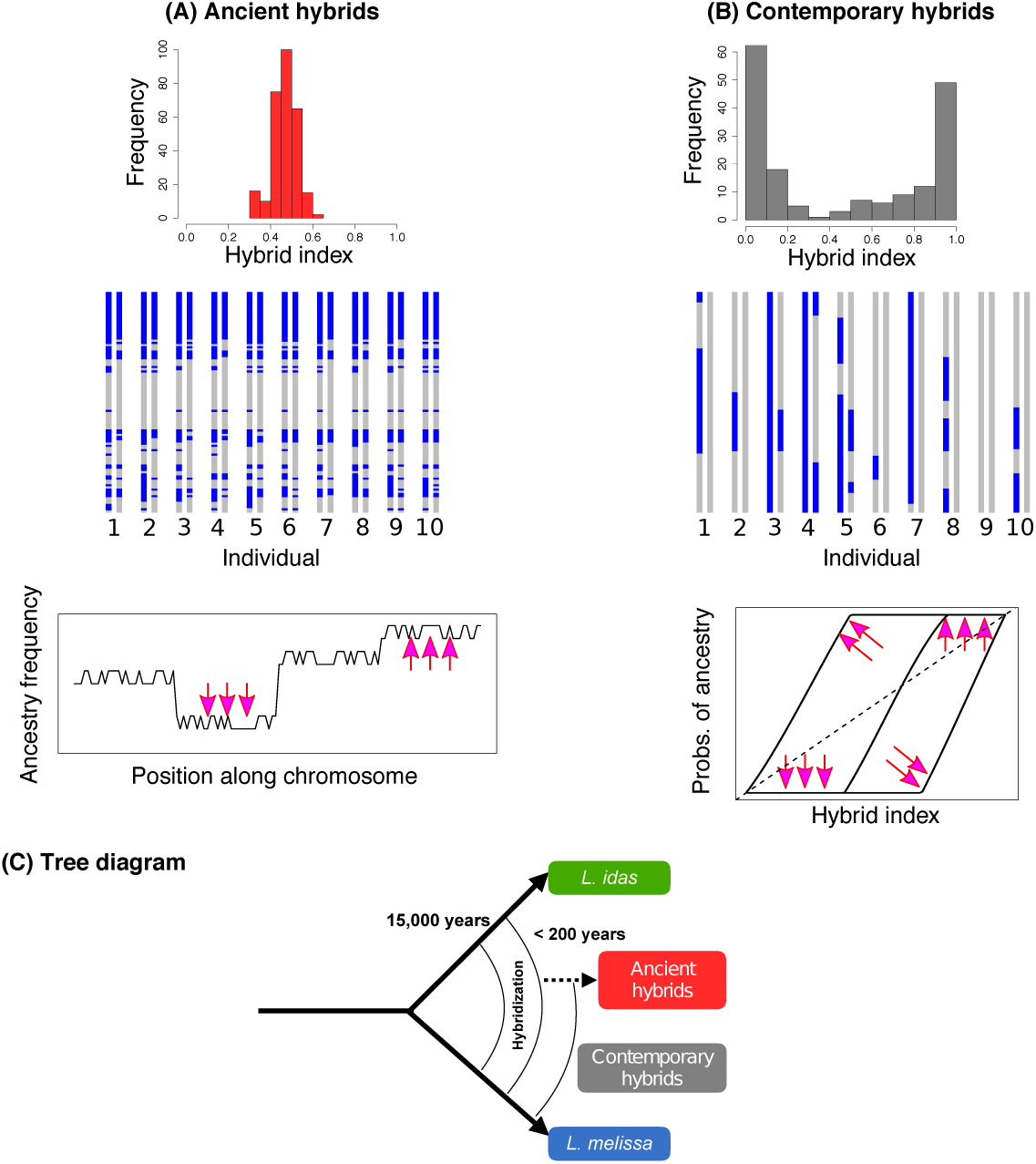
Conceptual overview and a comparative summary of genomic patterns expected in ancient hybrids (A) versus contemporary hybrids (B). Histograms show narrow (ancient) versus wide (contemporary) distributions of hybrid indexes. In ancient hybrids, ancestry blocks (gray versus blue segments) have been broken up by recombination and some have stabilized, that is fixed within the hybrid lineage. In contemporary hybrids, larger ancestry blocks are expected and these vary more among individuals. Plots show the expected effect of selection on ancestry frequencies and patterns of introgression in ancient and contemporary hybrids, respectively. In ancient hybrids, selection (pink arrows) shifts ancestry frequencies. In contemporary hybrids, selection shifts genomic clines for individual loci relative to the genome-wide average (dashed line). (C) Diagram represents the hypothesized history of hybridization in *Lycaeides*. Our results suggest Jackson Hole *Lycaeides* are ancient hybrids, with ancestry blocks from *L. idas* and *L. melissa* (akin to panel A), whereas Dubois are contemporary hybrids with ancestry blocks from Jackson Hole *Lycaeides* and *L. melissa* (akin to panel B with colors denoting Jackson Hole versus *L. melissa* ancestry).

**Figure 2:**
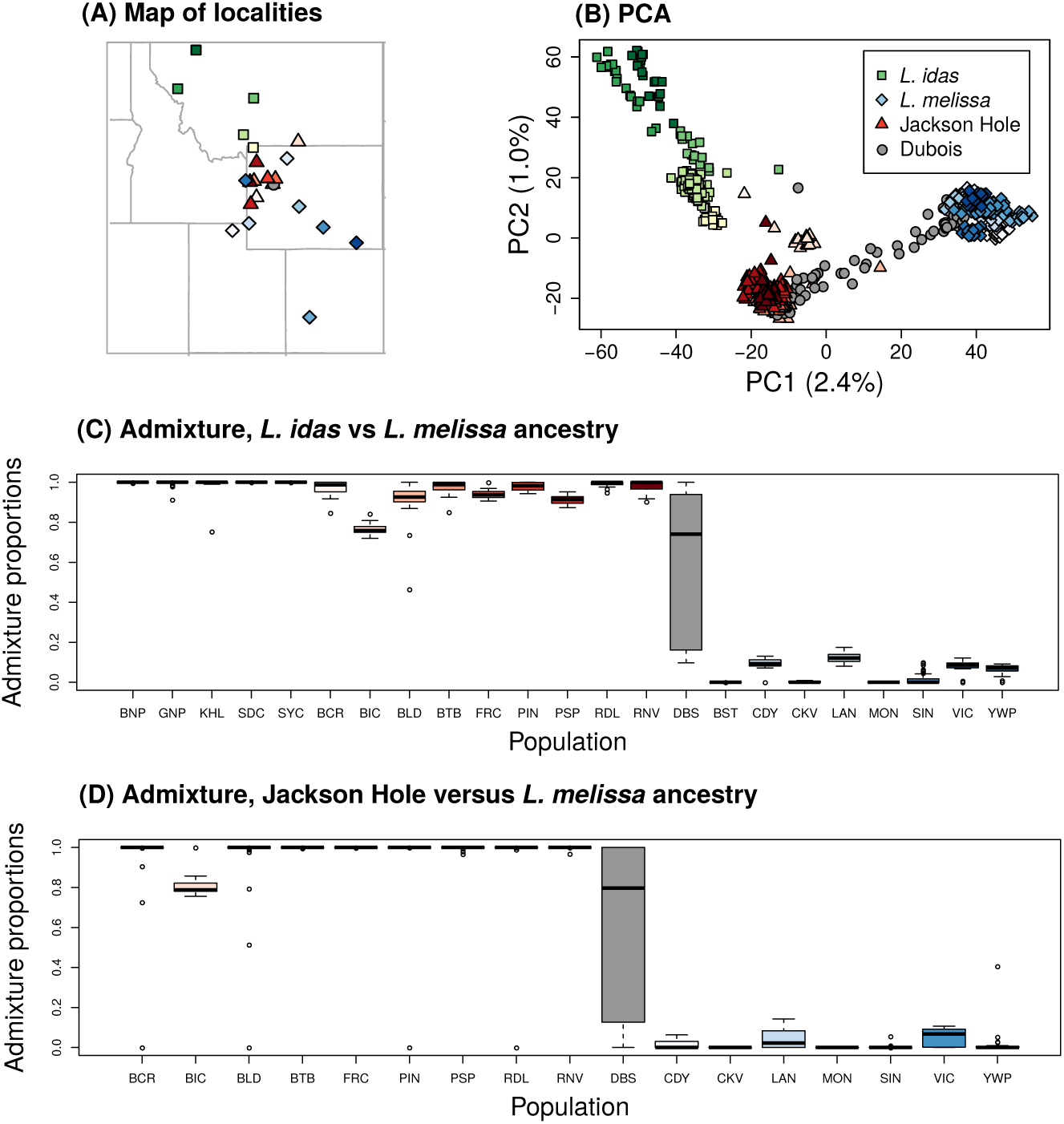
(A) Map of sample locations with points of shapes based on nominal taxa and colored for different populations within taxa. (Table S1). (B) Ordination of genetic variation via principal component analysis (PCA). Points denote individuals (a few low-coverage individuals were removed for visualization). (C) Boxplots of admixture proportion estimates from `entropy` with *k* = 2 source populations for all populations included in the study. Tick marks below the plots identify populations based on the population abbreviations in Table S1. (D) Boxplots of admixture proportion estimates from `entropy` with *k* = 2 source populations for all populations except *L. idas*. Tick marks below the plots identify populations based on the population abbreviations in Table S1. Source data are provided as a Source Data file.

### Genome composition and genome stabilization in the ancient Jackson Hole hybrid lineage

We next quantified genome-wide variation in the frequency of *L. idas* versus *L. melissa* ancestry segments in each of nine Jackson Hole *Lycaeides* populations (Table S1). This includes a population adjacent to the Dubois hybrid zone (Bald Mountain [BLD] which is ~5 km from Dubois), and much more distant populations (*>*100 km from Dubois, Fig. 2A). We focused this analysis on 1164 ancestry informative markers (AIMs; SNPs with allele frequency differences between *L. idas* and *L. melissa ≥* 0.3). We estimated ancestry segment frequencies using the correlated beta process model implemented in `popanc` (ver. 0.1; [15]). This method is similar to a hidden Markov model and accounts for the expected auto-correlation in ancestry along chromosomes, but allows ancestry frequencies to vary along the genome. Thus, it is suitable for quantifying ancestry frequencies in partially stabilized, old or ancient hybrid lineages. The average frequency of *L. idas* ancestry varied from 0.63 (RDL) to 0.51 (BCR), resulting in a modest northwest-southeast cline in mean genome composition (consistent with [29]) (Figure 3). However, ancestry frequencies varied considerably within and among chromosomes, with 7.7–33.2% of the genome fixed or nearly fixed (i.e., frequency *≥* 0.95) for *L. melissa* or *L. idas* ancestry (Figure 3, S14). More of the genome was fixed (or nearly fixed) for *L. idas* ancestry than *L. melissa* ancestry (mean = 10.9% versus 3.5%), and overall rates of fixation were higher on the Z chromosome than on autosomes (mean = 30.1% on the Z). Similar results were obtained when only male butterflies were analyzed (Fig. S15, S16).

**Figure 3:**
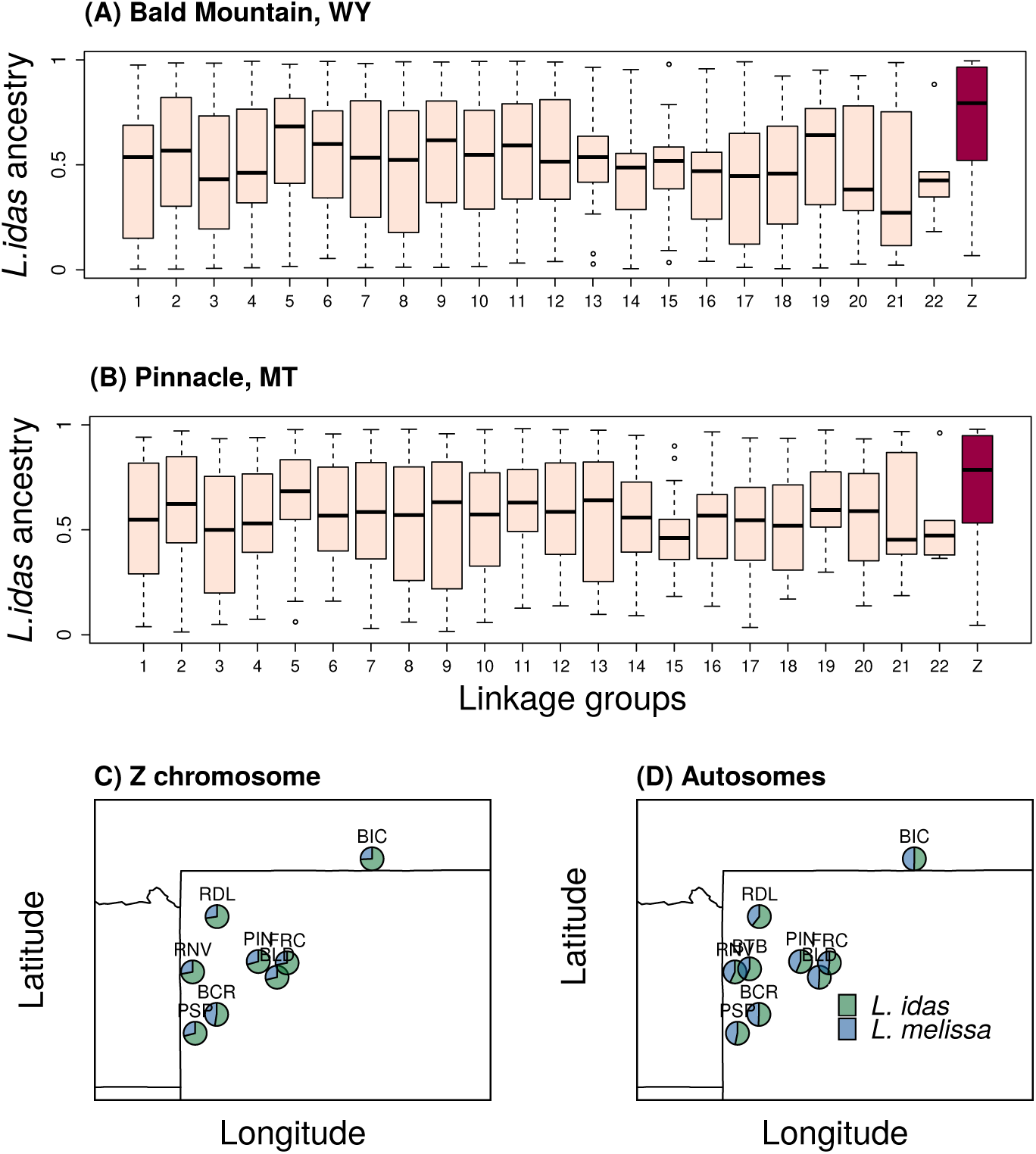
Boxplots show the distribution of *L. idas* ancestry across ancestry informative SNPs (AIMs) for each linkage group in two representative Jackson Hole *Lycaeides* populations–Bald Mountain, WY (BLD) (A) and Pinnacle, WY (PIN) (B). See Fig. S16 for additional populations. The Z sex chromosome is shown in red. Panels (C) and (D) show maps with pie charts reflecting the proportion of *L. idas* and *L. melissa* ancestry (mean) for the Z chromosome (C) and autosomes (D) for each of the nine populations (see Table S1 for population IDs). Source data are provided as a Source Data file.

Regions of exceptionally high *L. idas* ancestry frequencies were especially pronounced on the Z chromosome, and this held across all nine Jackson Hole populations (21.3–32.0% of the 225 AIMs on the Z had *L. idas* ancestry frequencies *≥*0.95). Specifically, randomization tests showed that the 10% of AIMs with the highest *L. idas* ancestry frequencies were found on the Z chromosome 1.28 to 3.32 times more often than expected by chance (Table S2). In contrast, we found little evidence that regions of high *L. melissa* ancestry were over represented on the Z chromosome (Table S3). Genomic regions with the highest levels of *L. idas* ancestry (i.e., the top 10% of AIMs with the highest *L. idas* ancestry) were in or near (within 1000 bp) genes (observed = 64, x-fold enrichment = 1.22, *P* = 0.03) and in or near gene coding sequences (observed = 56, x-fold enrichment = 1.25, *P* = 0.02) more often than expected by chance (Table S4). A similar pattern held for regions of high *L. melissa* ancestry (Table S5). Finally, regions of majority *L. melissa* ancestry (*>*50%) were less common on larger chromosomes (Pearson *r* = −0.5, *P* = 0.015), which instead harbored a greater proportion of genetic regions fixed or nearly fixed for *L. idas* ancestry (*>*95%) (Pearson *r* = 0.48, *P* = 0.021) (this pattern was weaker when the Z chromosome was excluded; Fig. S17). Such a pattern is expected when selection acts on many loci and against alleles from the minor parent (the one that contributes less to overall ancestry) and when recombination (per bp) is lower on larger chromosomes, as neutral minor parent alleles then have less opportunity to recombine away from deleterious minor parent alleles on larger chromosomes [25].

### Patterns of restricted and directional introgression in the Dubois hybrid zone

We then fit a Bayesian genomic cline model with `bgc` (ver. 1.04b; [33]) to quantify genome-wide variability in introgression between *L. melissa* and Jackson Hole *Lycaeides* in the Dubois hybrid zone. This method estimates clines in ancestry for individual genetic loci (e.g., SNPs) along a genome-average admixture gradient [34, 14] (Fig. S18). As such, it can be applied in cases in which hybrid zones are confined to a single geographic locality, such as the Dubois population. Unlike the method used for the ancient Jackson Hole hybrids, the genomic cline method performs best when a wide range of hybrids with different genome compositions exist, as is the case for the Dubois hybrid zone. Whereas our analysis of the Jackson Hole populations defined ancestry with respect to *L. idas* or *L. melissa*, here we classify genomic regions as having been inherited from Jackson Hole or *L. melissa*. We once again focused on the 1164 AIMs (these SNPs showed appreciable allele frequency differences between the reference Jackson Hole and *L. melissa* populations used to define source population ancestry in this analysis, e.g., mean = 0.28, with differences greater than 0.1 for 78% of the AIMs). We detected credible variation in patterns of introgression across the genome (defined as cases where Bayesian 95% credible intervals [CIs] for genomic cline parameters did not span zero; Fig. 4). Of the 1164 AIMs, 34 (2.9%) showed credible evidence of restricted introgression (95% CIs for cline parameter *β* > 0) and constitute candidates for genomic regions harboring barrier loci between *L. melissa* and Jackson Hole *Lycaeides* ([35]; Figure 4B). 189 of the AIMs (16.2%) had credible evidence of excess Jackson Hole *Lycaeides* introgression relative to genome-average expectations (i.e., directional introgression of Jackson Hole alleles; 95% CIs for cline parameter *α* > 0), and 273 AIMs (23.5%) had credible evidence of excess *L. melissa* introgression (i.e., directional introgression of *L. melissa* alleles; 95% CIs for cline parameter *α* < 0). Estimates of *α* were mostly independent of the degree of difference in allele frequencies between Jackson Hole and *L. melissa* (e.g., Pearson *r* = 0.007; *r* = −0.041 for *|α|*), but allele frequency differences explained 6.9% of the variation in point estimates of *β* and 20.9% of the variation in whether estimates of *β* were credibly greater than 0 (the latter likely reflects a combination of biological and statistical causes, e.g., high allele frequency differences likely reflect reduced introgression over time but also result in increased power to detect credible evidence of restricted introgression). Similar results were obtained when only male butterflies were analyzed; males carry two copies of the Z chromosome (Fig. S19).

**Figure 4:**
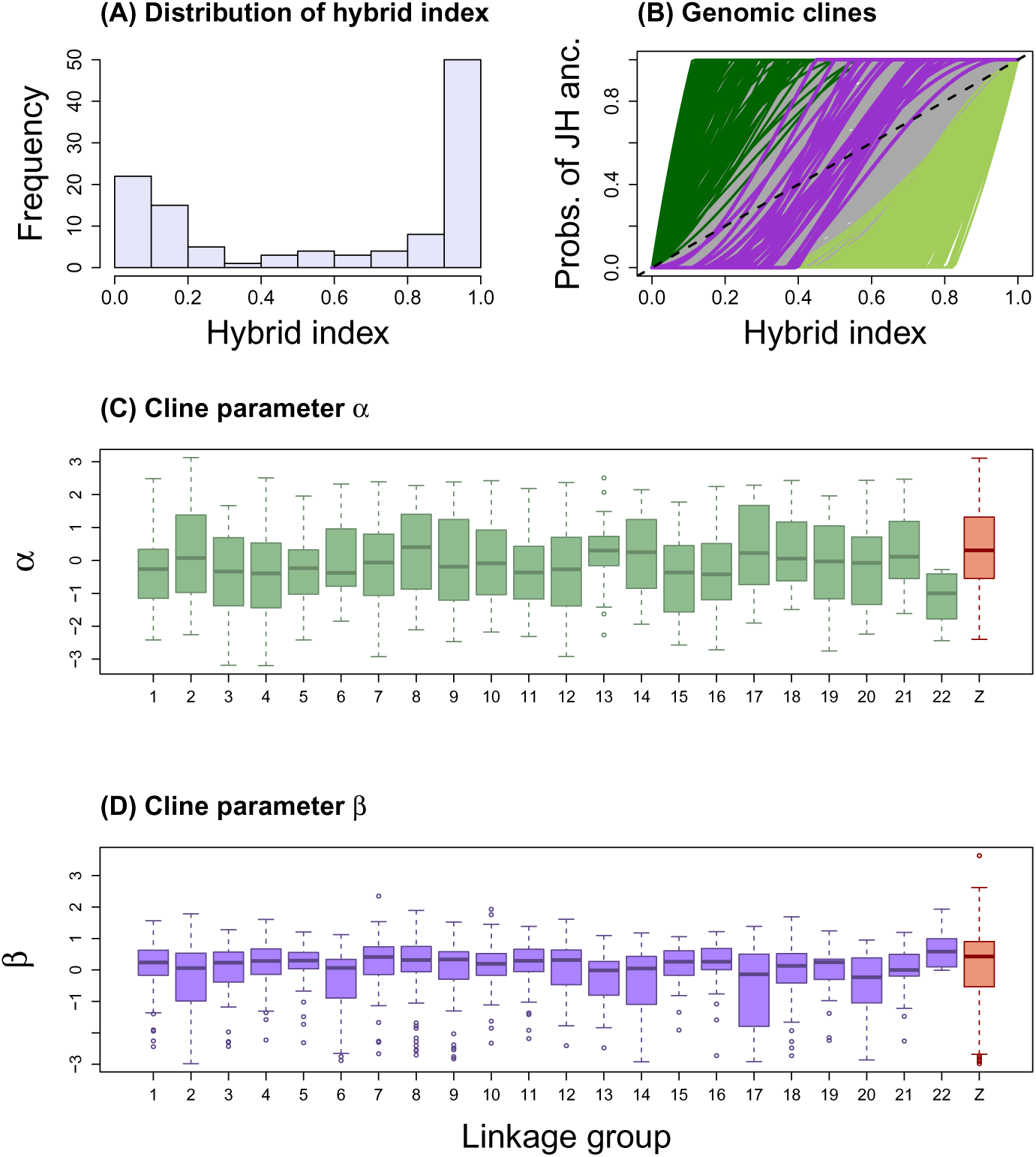
Summary of the genomic clines analysis. (A) The histogram depicts the distribution of hybrid indices in the Dubois hybrid zone. (B) This plot shows estimated genomic clines for a subset of ancestry informative SNPs (AIMs). Each solid line gives the estimated probability of Jackson Hole (JH) ancestry for an AIM. Green lines denote cases of credible directional introgression (95% CIs for *α* that exclude zero) and purple lines denote credible cases of restricted introgression (95% CIs for *β* > 0) (gray lines denote clines not credibly different from the genome-average). The dashed line gives the null expectation based on genome-wide admixture. Boxplots show the distribution of cline parameters *α* (C) and *β* (D) across loci for each linkage group. Source data are provided as a Source Data file.

Genetic loci exhibiting directional or restricted introgression were distributed across the 23 *Lycaeides* chromosomes (Figure 4B). However, randomization tests showed that AIMs with restricted introgression (34 loci with *β* > 0) and those with excess Jackson Hole introgression (189 loci with *α* > 0) were found on the Z sex chromosome more than expected by chance (*β* > 0, number on Z = 31, x-fold enrichment = 4.70, *P* < 0.001; *α* > 0, number on Z = 67, x-fold enrichment = 1.84, *P* < 0.001) (Fig. 4C,D). This was not true for AIMs with directional introgression of *L. melissa* alleles (273 loci with *α* < 0, number on Z = 37, x-fold enrichment = 0.70, *P* = 0.998). A greater proportion of AIMs exhibited restricted introgression on larger chromosomes (Pearson correlation between chromosome size and proportion of AIMs with *β* > 0, *r* = 0.46, *P* = 0.03), but this pattern was contingent on the Z chromosome and similar patterns were not seen with respect to directional introgression AIMs (Fig. S20). There was little evidence that genetic loci showing exceptional patterns of introgression in Dubois were clustered on genes or other structural genomic features (we did find some evidence of AIMs with credible *L. melissa* introgression being over-represented in coding sequences and proteins; Tables S6, S7 and S8).

### Introgression in contemporary hybrids predicts genome composition in ancient Jackson Hole hybrids

Regions of the genome showing exceptional introgression in the contemporary Dubois hybrid zone coincided with genomic regions exhibiting extreme ancestry frequencies in the ancient Jackon Hole hybrid lineage to a much greater extent than expected by chance (Figure 5). For example, AIMs exhibiting restricted introgression in the Dubois hybrid zone (top 10% of AIMs with the highest *β* values from `bgc`) overlapped with those with the highest *L. idas* ancestry frequencies in Jackson Hole (top 10%) about four times more often than expected by chance (observed = 51, x-fold enrichment = 4.41, *P* < 0.001; Figure 5E). The degree of overlap increased when we considered more extreme cutoffs for the comparison, up to the top 1% of AIMs with the greatest degree of restricted introgression and highest *L. idas* ancestry frequencies (x-fold enrichments ranged from 4.41 to 42.48; Figure 5G). Similar patterns were observed when comparing AIMs with evidence of excess directional introgression of Jackson Hole alleles in Dubois (top 10% with highest values or *α*) and those with the highest (i) *L. idas* (observed = 22, x-fold enrichment = 1.91, *P* = 0.0014) or (ii) *L. melissa* (observed = 26, x-fold enrichment = 2.24, *P* < 0.001) ancestry frequencies in Jackson Hole *Lycaeides* (Figure 5A,C). The strength of this signal of excess overlap was again more pronounced when considering more extreme cutoffs up to the top 1% of AIMs (x-fold enrichment ranged from 1.91 to 16.93; Fig. 5F). In contrast, we found no evidence that AIMs with excess directional introgression of *L. melissa* alleles in Dubois (top 10% with lowest values or *α*) coincided with those AIMs with the highest *L. idas* or *L. melissa* ancestry frequencies in Jackson Hole *Lycaeides*, nor evidence of excess overlap between AIMs exhibiting restricted introgression in Dubois and those with the highest *L. melissa* ancestry in Jackson Hole (Figs. 5B,D,F).

**Figure 5:**
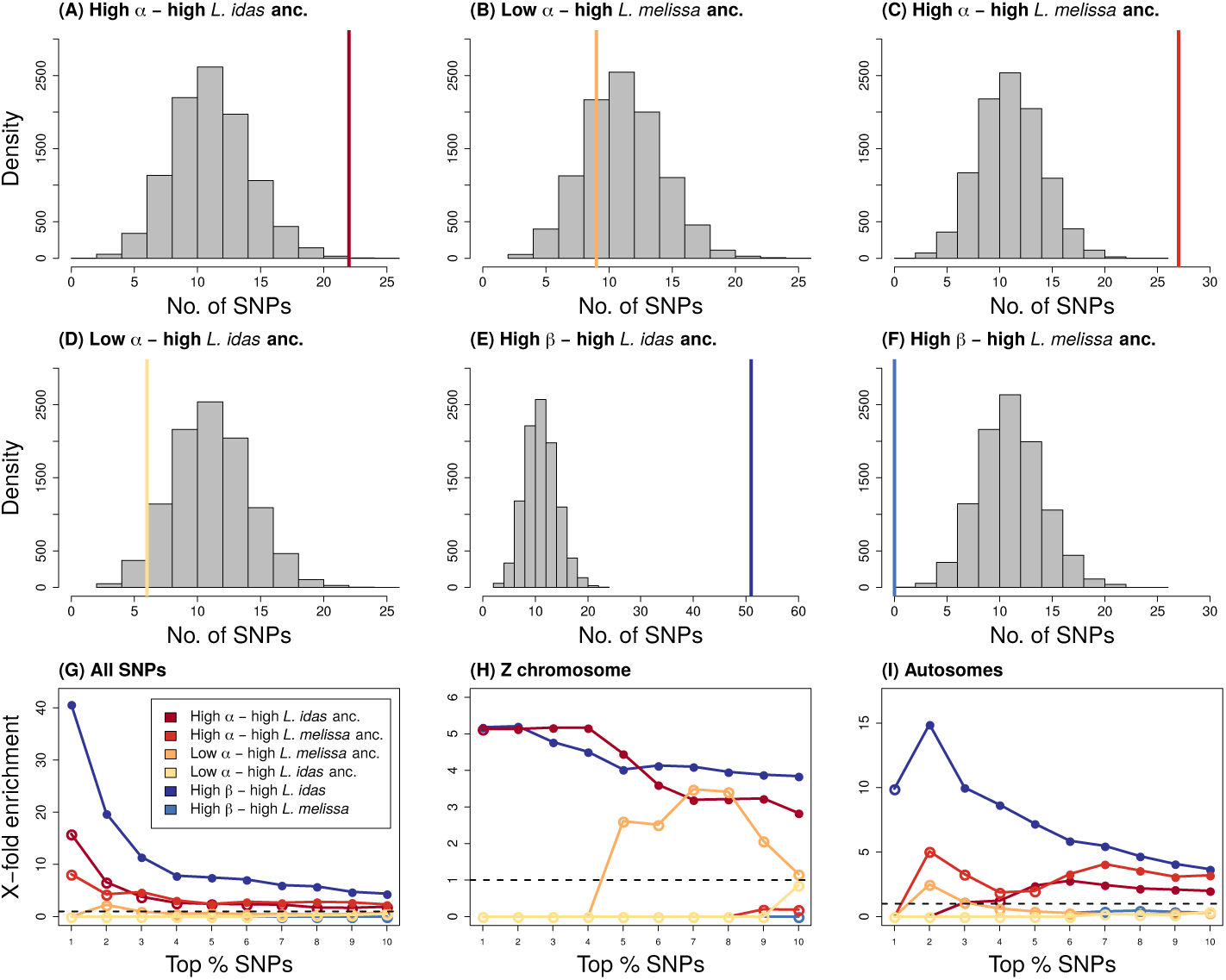
Expected and observed numbers of SNPs with exceptional patterns of introgression in the Dubois hybrid zone and extreme ancestry frequencies in Jackson Hole *Lycaeides*. Panels A-F show results when considering the top 10% of AIMs in each category. Histograms give null expectations from randomization tests and vertical solid lines show the observed number of AIMs exhibiting a given pattern. Comparisons shown are: directional introgression of Jackson Hole alleles (high *α*) and high *L. idas* ancestry (in Jackson Hole) (A), directional introgression of *L. melissa* alleles (low *α*) and high *L. melissa* ancestry (B), directional introgression of Jackson Hole alleles (high *α*) and high *L. melissa* ancestry (C), directional introgression of *L. melissa* alleles (low *α*) and high *L. idas* ancestry (D), restricted introgression (high *β*) and high *L. idas* ancestry (E), and restricted introgression (high *β*) and high *L. melissa* ancestry (F). Panels G-I show how these results are affected by considering different levels of stringency (i.e., by examining the most extreme 10% to the top 1% of AIMs with each pattern), and when considering only the Z chromosome (G) or only the autosomes (H). Here, circles denote the ratio of the observed to expected overlap from the null, and the circles are filled (*p* ≤ 0.05) or not (*p* > 0.05) to denote whether the overlap is greater than expected by chance. Source data are provided as a Source Data file.

Evidence of consistency (i.e., predictability) of genome composition between contemporary Dubois hybrids and the ancient Jackson Hole hybrids comes from both the autosomes and Z sex chromosome. Specifically, when we repeated the above comparisons with either the 22 autosomes or the Z chromosome alone, we obtained mostly similar results. For example, AIMs showing restricted introgression (top 10% with the highest values for *β*) and those with high *L. idas* ancestry frequencies in Jackson Hole *Lycaeides* (i.e., closest to being fixed for *L. idas* ancestry) coincided 3.7 times (observed = 37, *P* < 0.0001) and 3.8 times (observed = 38, *P* < 0.0001) more often than expected by chance for the autosomes and Z chromosome, respectively (Fig. 5H,I). Similarly, AIMs showing evidence of excess directional introgression of Jackson Hole alleles in Dubois (top 10%) and the highest frequencies of *L. idas* ancestry in the Jackson Hole populations coincided 1.9 (autosomes, observed = 22, *P* = 0.0018) and 2.8 (Z, observed = 12, *P* < 0.0001) times more often than expected by chance (Fig. 5G,H). We obtained similar results when defining AIMs as SNPs with an allele frequency difference of greater than 0.2 rather than 0.3 (2126 SNPs) (Figure S21), and when basing our analyses only on male butterflies, which carry two copies of the Z chromosome (Figs. S22, S23). Consistent results were also obtained when considering AIMs with the highest *L. idas* or *L. melissa* ancestry together (i.e., closest to being fixed for *L. idas* or *L. melissa* ancestry; Fig. S24).

We next analyzed the potential functional significance of genetic regions harboring the AIMs that were in the top 10% for both restricted introgression in Dubois (high *β*) and high *L. idas* ancestry frequencies in the ancient Jackson Hole *Lycaeides* lineage. We focus on this set of 51 loci as we think these are our best candidates for tagging regions of the genome harboring barrier loci, that is regions of the genome that constitute the basis of (partial) reproductive isolation among these lineages. This assertion stems from the fact that these loci exhibited restricted introgression in the contemporary hybrid zone combined with extreme ancestry frequencies (putatively arising from selection) in the Jackson Hole populations (specifically, high *L. idas* ancestry frequencies as we did not detect excess overlap between AIMs exhibiting restricted introgression and those with high *L. melissa* ancestry frequencies). Twenty-nine of these 51 AIMs were in or near genes (i.e., within 1000 bps of annotated gene boundaries) (in general, these barrier loci were not over-represented in or near specific structural features of the genome; Table S9, also see Tables S10 and S11). These genes exhibited a range of predicted functions (Tables S12, S13), with several genes standing out as being of particular interest. This includes the Z-linked 6-phosphogluconate dehyrdogenase gene, *6-pgd* (Fig. S25), which is associated with cold hardiness and diapause in other insects [36, 37, 38], and constitutes or is linked to a barrier locus in swallowtail butterflies [39, 40, 41]. Four AIMs were in or near two immunoglobulin superfamily genes; these were also on the Z chromosome (Tables S12, S13, Fig. S25). This gene superfamily is crucial for pathogen defense in insects [42], and is associated with reproductive isolation in mice [43]. Also among this set of genes was an autosomal olfactory receptor/odorant binding gene (Fig. S25). Such genes are known to affect host plant use in other butterflies [44, 45]. One of the 51 AIMs was within the autosomal nuclear pore gene *Nup93* (Tables S12, S13). This is part of the nuclear pore complex, which is involved in multiple Dobzhansky-Muller incompatabilities in *Drosophila*. Finally, two AIMs in *armadillo* or *armadillo*-like genes (one autosomal and one Z-linked) that are involved in the Wnt-signalling pathway and affect wing development in *Heliconius* and other butterflies were among this set of candidate barrier loci [46]. A diversity of functions were also predicted for the subset of AIMs with both high directional introgression of Jackson Hole alleles in Dubois and high *L. idas* ancestry in Jackson Hole *Lycaeides*; this includes immunoglobulin superfamily genes (Tables S14, S15) (also see Tables S16 and S17).

### Extending predictability to additional, ancient hybrid lineages

We next asked whether patterns of introgression in the contemporary, Dubois hybrid zone were also predictive of patterns of ancestry in two additional, ancient hybrid lineages from the Sierra Nevada and Warner mountains of western North America [6, 32, 28]. These ancient hybrid lineages occupy alpine habitat on isolated mountains ~900–1000 kilometers from Dubois. Past work suggests that these ecologically distinct populations arose within the past two million years (and perhaps more recently) following hybridization between *L. anna* and *L. melissa* (with a possible contribution from *L. idas* or a close relative of *L. idas*/*L. melissa*) [6, 47, 32, 28]. Using whole genome sequences from *L. anna*, *L. melissa*, *L. idas*, and the Sierra Nevada and Warner mountain lineages, we estimated phylogenetic relationships for the entire genome and for 1000 SNP windows along each chromosome to characterize patterns of ancestry in the ancient hybrids. Based on the results from Dubois and the Jackson Hole populations, we predicted reduced *L. melissa* ancestry on the Z chromosome, and topologies suggesting reduced introgression for the candidate barrier loci (i.e., regions containing the 51 AIMs with restricted introgression in Dubois and high *L. idas* ancestry frequencies in Jackson Hole).

Consistent with past work, the whole-genome consensus or “species” tree showed that the Warner mountain population was intermediate between *L. anna* and *L. melissa*/*L. idas*, whereas the Sierra Nevadan population was genetically more similar to *L. anna* (Fig. 6). Nonetheless, trees based on 1000 SNP windows varied across the genome (Fig. S26). The species tree was the most common topology overall (29.9%), but several “introgression” trees were also common, especially on the autosomes (Figs. 6, S27, S28). Whereas the trees that differ from the species tree could reflect incomplete lineage sorting, tree topologies were auto-correlated along the genome which suggests that introgression has contributed to these patterns as well (Fig. S28).

**Figure 6:**
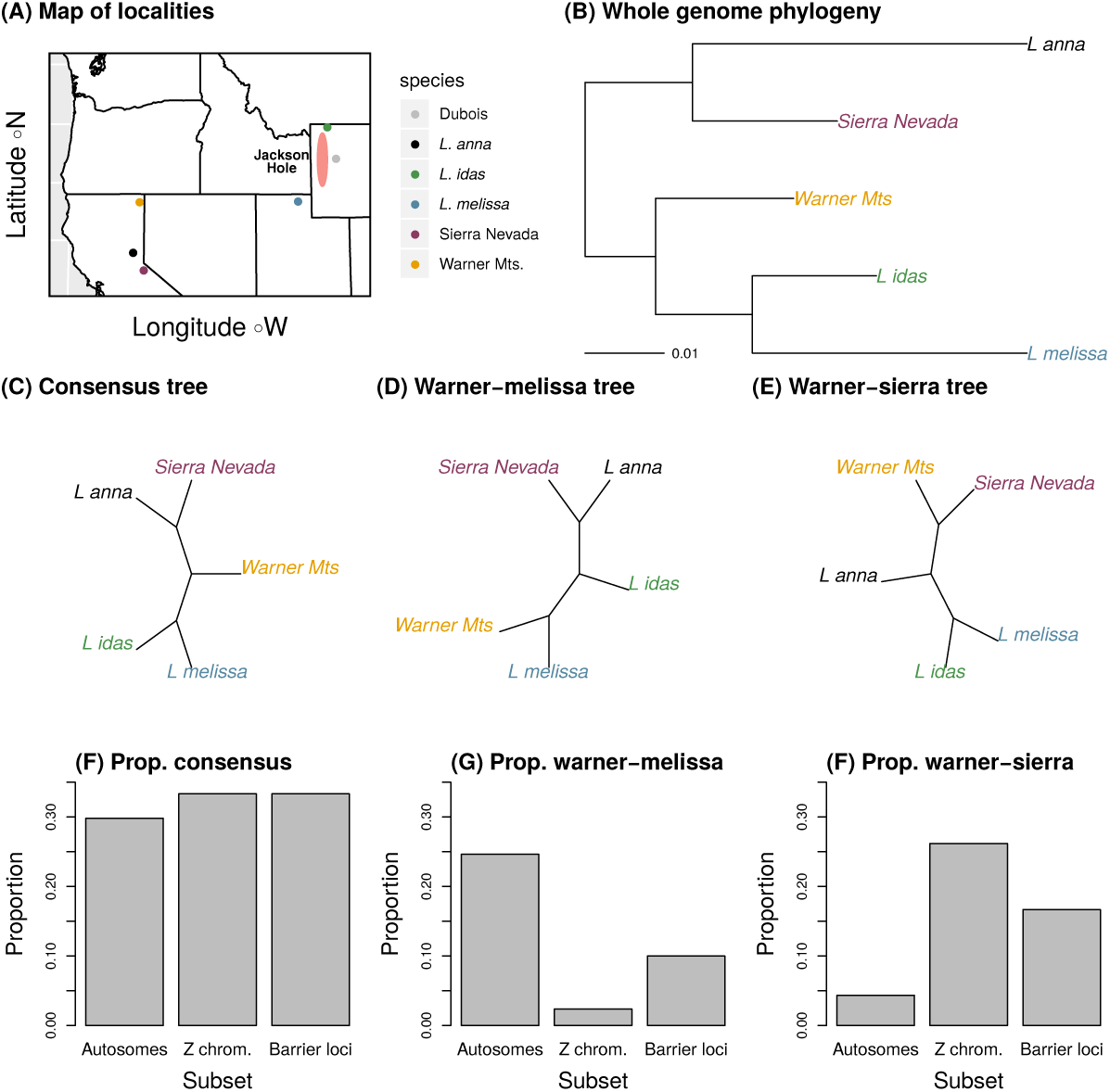
Whole-genome phylogenetic analyses. Panel (A) shows a map of the sampling localities in the western USA; Dubois is included for reference but was not sampled for whole-genome phylogenetics. The Jackson Hole ancestry cline (i.e., the range of the ancient Jackson Hole hybrids) is shown for reference as well (red zone). Panel B shows the midpoint rooted maximum likelihood phylogram inferred from the whole-genome SNP set (2,013,201 autosomal SNPs). Panels (C), (D) and (E) show three common unrooted tree topologies inferred from 1000 SNP windows. The tree in panel (C) is the most common topology and matches the whole-genome tree in (B). Trees in panels (D) and (E) differ by grouping the Warner Mts. population with *L. melissa* or the Sierra Nevada population, respectively. Panels (F), (G), and (H) show the proportion of 1000 SNP topologies matching the trees in (C)–(E) for autosomal windows, Z chromosome windows, and the 1000 SNP windows that contain the candidate barrier loci (see main text). There is a significant deficit of the topology in (D) on the Z chromosome and among the barrier loci (x-fold = 0.10, *P* < 0.001 and x-fold = 0.43, *P* = 0.047, respectively), and a significant excess of topology (E) on the Z chromosome and among the barrier loci (x-fold = 5.36, *P* < 0.001 and x-fold = 3.62, *P* = 0.008). See Fig. S26 for the full set of tree topologies recovered. Source data are provided as a Source Data file.

The species (i.e., consensus) tree was seen about 30% of the time for autosomal windows, Z chromosome windows and the 1000 SNP windows that contained the 51 candidate barrier loci (Fig. S27). The second most common autosomal tree suggested introgression from *L. melissa* into the Warner mountain lineage (24.6% of topologies). As predicted, this tree was significantly and substantially underrepresented on the Z chromosome (2.3%, x-fold = 0.10, *P* < 0.001) and for the candidate barrier loci (6.1%, x-fold = 0.43, *P* = 0.047) (Fig. 6). In contrast, a tree uniting the two ancient hybrid lineages was rare on the autosomes (4.3%), but commonly observed on the Z chromosome (26.2%, x-fold = 5.36, *P* < 0.001) and for the barrier loci (28.6%, x-fold = 3.62, *P* = 0.008) (Fig. 6). This suggests an ancient shared ancestry between these hybrid lineages that has been especially resistant to gene flow, or alternatively, Z-biased introgression between them. Finally, the introgression tree uniting the Warner mountain lineage and *L. melissa* was recovered more often on smaller chromosomes (Pearson correlation with LG size, *r* = −0.68, *P* < 0.001; autosomes only, *r* = −0.61, *P* = 0.003), whereas the tree uniting the ancient hybrid lineages was more common on large chromosomes (Pearson correlation with LG size, *r* = 0.55, *P* = 0.006; autosomes only, *r* = 0.46, *P* = 0.033) (Fig. S29).

To explicitly test that the above patterns were due to differential introgression, we calculated admixture proportions (*f_d_*) for each LG in the Warner mountain and Sierra Nevada lineages [48]. We found reduced introgression on the Z in both lineages (Sierra Nevada, *P* = 0.043; Warner mountain, *P* < 0.001; Fig. S30A,B), and evidence of a negative association between admixture and chromosome size in the Sierra Nevada lineage (*β* = 1.6*e^−^*^9^, std. error = 6.4*e^−^*^10^, *P* = 0.02; Fig. S30C,D). Thus, we find evidence of patterns of introgression and ancestry in these ancient hybrids consistent with patterns observed in the Dubois hybrid zone and ancient Jackson Hole hybrid lineage. Because the majority of the candidate barrier loci were Z-linked, the signal for these loci is not independent of the signal for the Z chromosome.

## Discussion

Biologists have endeavored to connect microevolutionary processes to macroevolutionary changes that occur over longer periods of time (e.g., [49, 50, 51]). Here we illustrate one way in which this gap can be bridged by showing that microevolutionary patterns and processes in a contemporary hybrid zone predict the genome composition of ancient, (partially) stabilized hybrid taxa (a macroevolutionary-scale outcome). This mirrors past work in *Helianthus* showing the genome composition of ancient hybrid sunflower species could be predicted from evolution in synthetic hybrids and QTL mapping experiments [52, 22]. However, our results are novel in that they demonstrate that the genomic outcomes of hybridization can be predicted from natural hybrids as well, and despite differences in the ecological and genomic context of the different hybridization events. Additional work in other organisms where ancient and contemporary hybrids are found is needed to determine the generality of this result (e.g., *Populus* and *Helianthus*; [53, 54, 55]). Beyond this, genomic analyses of multiple lineages at various points along the continuum from hybrid zone to stabilized hybrid species could prove particularly interesting. Such analyses could help distinguish between consistency arising from rapid fixation of ancestry segments versus sustained, consistent selection pressures over long periods of time. More generally, our results suggest that genomic analyses of both hybrid zones and hybrid lineages or species might offer a particularly tractable framework for assessing the ways in which and degree to which speciation (especially hybrid speciation) might be predictable from microevolutionary processes of selection and recombination.

Patterns of parallel or repeatable evolution strongly suggest a major role for the deterministic process of natural selection [56, 57]. Thus, genes and gene regions with restricted introgression in Dubois and extreme ancestry frequencies or phylogenetic relationships in Jackson Hole and the other ancient hybrid lineages represent strong candidates for harboring barrier loci (i.e., speciation genes). Given the differences in environment and ecological context across these instances of admixture, our results suggest that many of these putative barrier loci operate in a manner that is not highly environment dependent, which is consistent with a major role of intrinsic incompatibilities in the speciation process [58, 59]. Predictability (i.e., consistency) was highest when considering the Z chromosome, where we detected a lack of *L. melissa* ancestry across the hybrid zone and hybrid lineages. This could be explained by an increased efficacy of selection against Z loci in hemizygous individuals or linkage and a higher density of selected loci on the Z in general [60, 61]. An excess of intrinsic incompatibilities arising from epistatic interactions is an especially likely explanation, as the hybrid populations generally harbored less *L. melissa* ancestry (lower *L. melissa* ancestry could reflect selection or demographic aspects of the hybridization process). Nonetheless, our ability to predict the genome composition of ancient hybrids, especially the Jackson Hole lineage, held even when excluding the Z. Thus, while our results are consistent with theory and other empirical studies that suggest a disproportionate role for sex chromosomes in speciation (e.g., [59, 60, 21, 14]), this was not the sole factor that made genome composition predictable in this system. Indeed, our analyses highlighted candidate barrier loci on autosomes (e.g., an autosomal olfactory receptor/odorant binding gene and *Nup93*), as well as on the Z chromosome. Our results also suggest reduced introgression often occurs on longer chromosomes, which should undergo less recombination per base pair, in *Lycaeides*. This is consistent with selection acting on many loci scattered across the genome, and has been shown in several systems (e.g., [23, 16, 25]). In our case, this pattern was driven in part by the Z chromosome, which is one of the largest chromosomes, but which should not otherwise experience reduced recombination (i.e., recombination in butterflies is completely suppressed in the heterogametic sex, including recombination between homologous autosomes). We do not yet know whether similar results hold based on finer-scale variation in recombination rate within chromosomes, driven for example by structural variants or other regions of reduced recombination. However, our results do show that this pattern holds early and late in the hybridization and genome stabilization process, and that selection and recombination interact during this process in a manner that results in consistent or repeatable patterns of hybrid genome composition.

In conclusion, the fact that putative barrier loci in the Dubois hybrid zone exhibit extreme ancestry patterns in ancient hybrid lineages is consistent with the hypothesis that the same genes or gene regions that prevent species fusion during hybridization experience selection during the evolution of hybrid lineages or species. This hypothesis is also supported by studies in other systems showing that incompatibility loci contribute to hybrid speciation [52, 16], and is generally consistent with accumulating data suggesting the same genes are repeatedly involved in adaptation (e.g., [62, 63]). Thus, seemingly distinct facets of speciation, such as the maintenance of taxonomic boundaries across hybrid zones and the origin of novel biodiversity through admixture, may have a predictable, common genetic basis.

## Methods

### Genome assembly and annotation

We generated a new, chromosome-scale reference genome for *L. melissa* from information on proximity ligation of DNA in chromatin and reconstituted chromatin. Our previous genome assembly comprised 14,029 scaffolds (total assembly length = 360 megabase pairs [mbps]; scaffold N50 = 65 kilobase pairs [kbps]), which had been joined in a linkage map with 23 linkage groups (*Lycaeides* has 22 autosomes and a ZW sex chromosome system) [64, 65]. In the current study, we improved upon this assembly using DNA sequence data from Chicago and Hi-C libraries [66, 67]. Creation and sequencing of the Chicago and Hi-C libraries was outsourced to Dovetail Genomics. The new sequence data and our old assembly were combined using the HiRise assembler (also outsourced to Dovetail Genomics). The new *L. melissa* genome assembly has a final N50 of 15.5 mbps, and 90% of the genome comprises only 21 scaffolds. A whole genome comparative alignment with `mummer` (version 3.2 with the maximal unique matches setting; [68]) of this new genome assembly with the previously published genome assembly and linkage map showed that each of our previously defined linkage groups corresponds with one (or in one case two) of the new, large scaffolds.

We annotated structural and functional features of the new *L. melissa* genome using the `maker` pipeline (version 2.31.10) [69, 70]. This pipeline uses repeat masking, protein and RNA alignment and *ab initio* gene prediction to perform evidence-based gene prediction, which generates annotations that are supported by quality scores. Prior to using `maker`, we identified repeats in the *L. melissa* genome using `repeatscout` (version 1.0.5) [71]. This program identifies repeat elements, including tandem repeats and low complexity elements, and removes them from the genome. We took this approach to avoid missing repeat elements not already in standard data bases. We provided this *de novo* repeat library to `maker`, and used this along with `repbase` in `repeatmasker` to mask repetitive elements in the genome. `maker` can use protein and RNA sequence data for genome annotation. Since we lacked protein sequences for *L. melissa*, we downloaded 28 protein sequence fasta files from 15 butterfly species (see Table S18) from `LepBase` (version 4) [72]. We concatenated these fasta files to generate a protein sequence data base for `maker`. We used data from 24 *L. melissa* transcriptomes (Forister *et al.*, manuscript in prep.) as additional evidence for genome annotation. We first used `trimgalore` (version 2.6.6, https://github.com/FelixKrueger/TrimGalore) for adapter trimming and quality filtering of paired-end RNA sequences. We then used these trimmed reads to generate a *de novo* transcriptome assembly with `trinity` (version 2.6.6) [73, 74]. The assembled transcriptome was passed to `maker`.

We ran two rounds of the `maker` pipeline. We first ran `maker` without using any information from *ab initio* gene predictors (e.g., `augustus`), to generate *de novo* gene models for the *L. melissa* genome. We then ran the `maker` pipeline again and used the gene models from the first run to train two gene predictors: `augustus` and `snap`. We ran `snap` (version 2006-07-08) [75] by using models with AED scores of 0.25 or better and a length of 50 or more amino acids. We ran `augustus` (version 3.3) with the insect predictions [76]. We then used both of these sets of gene predictions in the second run of `maker`. We then used the output from `maker` to obtain functional annotations of the *L. melissa* genome. We assigned putative gene functions by using `blastp` to query the `maker` output against the UNIPROT/SWISSPROT database [77]. We also used `interproscan` [78] to add protein and gene ontology information to gene models. The final annotation included 11,247 putative genes, 48,765 putative coding sequences, and 8893 UTR sequences.

### DNA sequencing, alignment, and genetic variant detection

We analyzed genotyping-by-sequencing (GBS) data from 835 *Lycaeides* butterflies from 23 populations: eight *L. melissa* populations (N = 238 butterflies), five *L. idas* populations (N = 176 butterflies), nine Jackson Hole *Lycaeides* populations (N = 306 butterflies) and the Dubois hybrid zone (N = 115 butterflies) (Table S1). The sequence data from 643 of these butterflies were previously described in a study of admixture across the *Lycaeides* species complex [28]. Data from 192 of the butterflies were generated for the current study, and this includes many (but not all) of the Dubois individuals. We extracted DNA, generated genotyping-by-sequencing (GBS) libraries and sequenced these libraries following the protocols described in [28]. The GBS libraries were sequenced on an Illumina HiSeq 2500 (100 bp, single-end reads) by the Genome Sequencing and Analysis Facility at the University of Texas (Austin, TX).

We used `bwa` (version 0.7.17) to align the GBS sequences from 835 individuals to the draft *L. melissa* genome by using the `mem` algorithm [79, 80]. We ran `bwa` `mem` with a minimum seed length of 15, internal seeds of longer than 20 bp, and only output alignments with a quality score of *≥*30. We then used `samtools` (version 1.5) to compress, sort and index the alignments [81]. We used `samtools` (version 1.5) and `bcftools` (version 1.6) for variant calling. For variant calling, we used the recommended mapping quality adjustment for Illumina data (`-C 50`), skipped alignments with mapping quality less than 20, skipped bases with base quality less than 15, and ignored insertion-deletion polymorphisms. We set the prior on SNPs to 0.001 (`-P`) and called SNPs when the posterior probability that the nucleotide was invariant was *≤* 0.01 (`-p`). We filtered the initial set of variants to retain only those SNPs with sequence data for at least 80% of the individuals, a mean sequence depth of 2 per individual, at least 4 reads of the alternative allele, a minimum quality score of 30, a minimum (overall) minor allele frequency of at least 0.005, and no more than 1% of the reads in the reverse orientation (this is an expectation for our GBS method). We further removed SNPs with excessive coverage (3 standard deviations above the mean) or that were tightly clustered (within 3 bp of each other), as these could reflect poor alignments (e.g., reads from multiple paralogs mapping to the same region of the genome). Finally, because we combined data from two different sequencing runs, we also removed any SNPs with a difference in sequence coverage between the published and new data that was more than half the mean coverage for the two data sets combined. This left us with 39,139 SNPs for downstream analyses.

### Estimating genotypes, admixture proportions, population allele frequencies and linkage disequilibrium

We used the admixture model from `entropy` (version 1.2) [28] to obtain Bayesian estimates of genotypes and admixture proportions. This analysis was based on the full data set of 835 individuals and 39,139 SNPs. The admixture model in `entropy` is similar to that in `structure` [82], but differs by accounting for uncertainty in genotypes arising from limited sequence coverage and sequence errors, and by allowing simultaneous estimation of genotypes and admixture proportions [28]. We fit the model with *k* ∈ {2 … 5} source populations. For each value of *k*, we ran three Markov chain Monte Carlo (MCMC) chains, each with 15,000 MCMC iterations, a burnin of 5000 iterations and a thinning interval of 5. We used assignments from a discriminant analysis of principal components to initialize the MCMC algorithm; this speeds convergence to the posterior and avoids label switching during MCMC without affecting the posterior probability distribution [28]. We obtained genotype estimates as the posterior mean allele count for each individual and locus across chains and values of *k* (i.e., this integrates over uncertainty in the number of hypothetical source populations). We focused on admixture proportions for *k* = 2, as we were interested in the two nominal species and hybrids between them. We summarized patterns of population structure and admixture across the sampled populations and individuals based on these admixture proportions and a principal component analysis (PCA) of the genotypic data. We performed the PCA in `R` on the centered but not standardized genotype matrix with the `prcomp` function.

We used the expectation-maximization (EM) algorithm implemented in `estpEM` (version 0.1) [57] to estimate allele frequencies for each SNP (39,139 SNPs) and population (N=23 populations). The EM algorithm estimates allele frequencies while allowing for uncertainty in genotypes [83, 57]. We used the genotype like-lihoods calculated with `bcftools` for this analysis, and ran the algorithm with a convergence tolerance of 0.001 and allowing for 20 EM iterations. We used these allele frequency estimates to designate ancestry informative SNPs/markers (AIMs) as those with an allele frequency difference *≥*0.3 between *L. melissa* (mean of BST, SIN, CDY, and CKV) and *L. idas* (mean of KHL, SDC, and SYC) (population IDs are defined in Table S1).

Next, we calculated a metric of linkage disequilibrium (LD), the Pearson correlation coefficient between genotypes at pairs of loci, for all pairwise combinations of AIMs in each population where 20 or more butterflies were collected. Our goal was to ask whether LD was elevated in the Dubois hybrid zone, as predicted by theory (e.g., [84]). We first polarized genotypes such that positive LD (i.e., positive Pearson correlations) coincided with an association between coupling alleles, that is alleles more common in *L. idas* or *L. melissa*, whereas negative LD (i.e., negative Pearson correlations) coincided with associations between repulsion alleles, that is positive associations between *L. melissa* and *L. idas* alleles. Ongoing hybridization should cause a shift towards higher positive estimates of LD, even for unlinked markers (via admixture LD). Correlations were calculated in `R` (version 3.5.1).

Lastly, we used discriminant analysis of the genetic PCs to assign butterflies from Dubois to reference taxa. We did not do this to estimate admixture proportions, but rather to determine the likely taxonomic origin of the hybridizing taxa. We first ran k-means clustering on the PC scores (PCs 1 and 2) for all sampled *Lycaeides* except those from Dubois. We assumed three groups, allowed 50 starts and 50 iterations. This was done with the `kmeans` function in `R`. We then used the inferred clusters, which corresponded with *L. melissa*, *L. idas* and Jackson Hole *Lycaeides* (see Results), as training information for a linear discriminant analysis where we classified the Dubois individuals. We assumed prior probabilities of 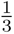 for assignment to each cluster. This was done with the `lda` function in `R`.

### Analysis of population ancestry in Jackson Hole-*Lxycaeides*

We estimated ancestry segment frequencies across the genome for each of the nine Jackson Hole ancient hybrid populations using the correlated beta process model implemented in `popanc` (ver. 0.1; [15]). This method is similar to a hidden Markov model and accounts for the expected autocorrelation in ancestry along chromosomes, but allows ancestry frequencies to vary along the genome. It is particularly well suited for cases where genome stabilization has begun but is not yet complete (that is, where populations consist of individuals with similar admixture proportions but where ancestry frequencies vary across the genome)[15]. We ran `popanc` for each of the Jackson Hole populations to estimate the frequency of *L. idas*-derived alleles along the genome. We used a window size of 3 SNPs and focused on the 1164 AIMs. We set SYC and KHL as representative of the *L. idas* parental species, and BST, SIN, CDY and CKV as the putative *L. melissa* parents. Maximum likelihood allele frequency estimates from estpEM were used as input for the program/analysis. We ran the MCMC analysis for each population with a 10,000 iteration chain, a 5000 iteration burn-in and thinning interval of 10. We based our inferences on point estimates (posterior mean) of *L. idas* ancestry frequencies for individuals populations or on averages of these estimates across all nine populations. We repeated this analysis with only males and with AIMs defined as those SNPs with an allele frequency difference of *≥*0.2 between *L. melissa* and *L. idas* to assess the robustness of our results. Randomization tests were used to ask whether and to what extent AIMs with the highest *L. idas* or *L. melissa* ancestry frequencies (top 10% in each case) were over-represented on the Z chromosome or in or near certain structural features of the genome such as genes, coding sequences, transposable elements and annotated protein sequences or motifs. Null expectations were derived from 1000 permutations of ancestry frequency estimates across the 1164 AIMs. Lastly, we asked whether largest chromosomes harbored more *L. idas* ancestry than smaller chromosomes (as noted in the Results, Jackson Hole *Lycaeides* inherited more of their genomes from *L. idas* than *L. melissa*). Such a pattern is expected when selection acts on many loci and against alleles from the minor parent (the one that contributes less to overall ancestry, in this case *L. melissa*) and when recombination (per bp) is lower on larger chromosomes (as has been seen in Lepidoptera), as neutral alleles from the minor parent then have less opportunity to recombine away from deleterious alleles from the minor parent when they occur on larger chromosomes [25].

### Estimating cline parameters in the Dubois hybrid zone

We fit Bayesian genomic clines for each of the 1164 AIMs using `bgc` (ver. 1.04b; [33]) to quantify genome-wide variability in introgression between *L. melissa* and Jackson Hole *Lycaeides* in the Dubois hybrid zone. This method estimates clines in ancestry for individual genetic loci (e.g., SNPs) along a genome-average admixture gradient [34, 14]. As such, it can be applied in cases where hybrid zones are confined to a single geographic locality, such as the Dubois population. Unlike the method used for the ancient Jackson Hole hybrids, the genomic cline method performs best when a wide range of hybrids with different genome compositions exist, as is the case for the Dubois hybrid zone. Deviations between genome-average introgression and introgression for each locus are measured with two cline parameters, *α* and *β*. Cline parameter *α* denotes an increase (for positive values of *α*) or decrease (for negative values of *α*) in the probability of ancestry from reference (parental) species 1 relative to null expectations from an individual’s hybrid index. Genomic cline parameter *β* describes an increase (positive values) or decrease (negative values) in the rate of transition from parental species 0 ancestry to parental species 1 ancestry along the genome-wide admixture gradient. When placed in a geographic context, *α* is equivalent to twice the shift in cline center, and *β* measures the decrease (or increase when *β* < 0) in cline width relative to the average [84, 14]. Cline parameters can be affected by genetic drift and selection in hybrids [35, 14]. However, in the absence of major geographic barriers to gene flow, high positive values of *β* (i.e., the equivalent of narrow clines in a geographic context) are most readily explained by selection.

We fit the `bgc` genomic clines model for the 115 *Lycaeides* butterflies from the Dubois hybrid zone. We used *L. melissa* (LAN, SIN, and CKV; N = 131 butterflies) and Jackson Hole *Lycaeides* (BLD and FRC; N = 94 butterflies) as reference or parental species for the analysis. These specific populations were chosen as the source populations because they were nearest to the Dubois hybrid zone. Using these populations as references, allele frequencies for the AIMs differed sufficiently between source populations to obtain meaningful estimates of ancestry (e.g., mean = 0.28, 78% with allele frequency differences *>* 0.1, see also simulations in [35]). Genotype likelihoods from `bcftools` were used as input for the analysis (the model fit incorporates uncertainty in genotype as captured by the genotype likelihoods). We ran the analysis using only the 1164 AIMs. We fit the model using MCMC, with five chains each with 25,000 iterations, a 5000 iteration burn-in and a thinning interval of 5. We inspected the MCMC output to assess convergence of chains to the stationary distribution and combined the output of the five chains. We repeated this analysis with only males and with AIMs defined as those SNPs with an allele frequency difference of *≥*0.2 between *L. melissa* and *L. idas* (2126 SNPs) to assess the robustness of our results.

We defined credible deviations from null expectations given genome-wide admixture as cases where the 95% credible intervals (specifically the equal-tail probability intervals) for *α* or *β* for a given locus (AIM) excluded 0. We focused specifically on cases of credible directional (*α* ≠ 0) or restricted (*β* > 0) introgression relative to the genome-wide average. We used randomization tests to ask whether (and to what extent) such loci were over (or under) represented on the Z chromosome or in or near (within 1000 bp of) certain annotated structural features of the genome, such as genes, coding sequences, transposable elements and annotated protein sequences or motifs. Null expectations were derived from 1000 permutations of cline parameter estimates across the 1164 AIMs. We also tested for correlations between chromosome size and the proportion of AIMs showing evidence of restricted (*β* > 0) or directional (*α* > 0 or *α* < 0) introgression.

### Quantifying consistency in genomic outcomes of hybridization between contemporary and ancient hybrids

We next tested for excess overlap between AIMs showing the most extreme ancestry frequencies in the ancient Jackson Hole hybrids and those with the greatest deviations from genome-wide average introgression in the contemporary Dubois hybrid zone. We focused primarily on the following six comparisons: (i) excess introgression of Jackson Hole alleles in Dubois (*α* > 0) and high *L. idas* ancestry in Jackson Hole, (ii) excess introgression of Jackson Hole alleles in Dubois (*α* > 0) and high *L. melissa* ancestry in Jackson Hole, (iii) excess introgression of *L. melissa* alleles in Dubois (*α* < 0) and high *L. idas* ancestry in Jackson Hole, (iv) excess introgression of *L. melissa* alleles in Dubois (*α* < 0) and high *L. melissa* ancestry in Jackson Hole, (v) restricted introgression in Dubois (*β* > 0) and high *L. idas* ancestry in Jackson Hole, and (vi) restricted introgression in Dubois (*β* > 0) and high *L. melissa* ancestry in Jackson Hole. We were especially interested in the last two comparisons (v and vi), as the restricted introgression AIMs constitute our best candidates for regions of the genome associated with reproductive isolation. We initially focused on the top 10% of AIMs in each of the categories (e.g., the 116 out of 1164 AIMs with the highest point estimates of *α* from `bgc`). We conducted randomization tests by permuting parameters across AIMs to test whether and to what extent AIMs in the top 10% for each category coincided more for each comparison than expected by chance. We conducted 10,000 permutations to generate null expectations. The null distribution was used to calculate a *P* value and x-fold enrichment for each comparison. As an example, an x-fold enrichment of 2.0 would indicate that twice as many AIMs exhibited a pair of patterns (e.g., restricted introgression in Dubois and high *L. idas* ancestry frequencies in Jackson Hole) as expected by chance (based on the mean of the null). We repeated these analyses considering the top 9%, 8%, 7%, 6%, 5%, 4%, 3%, 2% and 1% of AIMs in each category (i.e., across more and more extreme quantiles), and with only autosomal or only Z-linked AIMs. We also repeated this analysis with only males and with AIMs defined as those SNPs with an allele frequency difference of *≥*0.2 between *L. melissa* and *L. idas* to further assess the robustness of our results. As a final assessment of robustness, we considered three additional comparisons: (vii) excess introgression of Jackson Hole alleles in Dubois (*α* > 0) and high *L. idas* or *L. melissa* ancestry in Jackson Hole (i.e., genomic regions nearest fixation for either ancestry type, hereafter extreme ancestry), (viii) excess introgression of *L. melissa* alleles in Dubois (*α* < 0) and extreme ancestry in Jackson Hole, and (ix) restricted introgression in Dubois (*β* > 0) and extreme ancestry in Jackson Hole. We includes these comparisons as the direction of selection in Jackson Hole (for or against *L. idas* or *L. melissa* ancestry) might not predict the direction of selection in Dubois, but still the same subsets of loci could be affected by selection in both cases.

Finally, we extracted IPR and GO terms and descriptions generated by `maker` for the set of AIMs in the top 10% for each comparison where we had evidence of significantly greater overlap between categories than expected by chance. We filtered the IPR terms based on their hierarchical classifications (i.e., superfamily, family, domain or repeat). We retained unique terms and dropped terms that at lower or overlapping levels in the Interproscan database [78]. For example, if a SNP was annotated for a superfamily IPR term and multiple domains within the superfamily, we retained only the superfamily term.

### Whole genome phylogenomic analyses

We next asked whether patterns of introgression in the contemporary, Dubois hybrid zone were also predictive of patterns of ancestry in two additional, ancient hybrid lineages from the Sierra Nevada and Warner mountains of western North America [6, 32, 28]. To do this, we generated whole genome sequences from *L. anna* (from Yuba Gap, CA), *L. melissa* (from Bonneville Shoreline, UT), *L. idas* (from Trout Lake, WY), and the Sierra Nevada (from Carson Pass, NV), and Warner mountain (from Buck Mountain, CA) lineages. We extracted DNA from one female (ZW) butterfly per population using Qiagen’s MagAttract HMW DNA extraction kit (Qiagen, inc.) following the manufacture’s suggested protocol. We then outsourced library construction and sequencing to Macrogen Inc. (Seoul, South Korea). One standard paired-end shotgun library (180 bp insert) was constructed for each butterfly/lineage using a TruSeq library preparation kit (Illumina, Inc.). Each library was sequenced in its own lane on a HiSeq 2000 with 2 *×* 100 bp paired reads. We obtained *>* 20 million high quality (*≥* Q30) reads for each library (~8*X* coverage of each genome).

We ran `bwa` `mem` with a minimum seed length of 15, internal seeds of longer than 20 bp, and only output alignments with a quality score of *≥*30. We then used `samtools` (version 1.5) to compress, sort and index the alignments, and to remove PCR duplicates [81]. We used GATK’s HaplotypeCaller (version 3.5) to call SNPs across the five genomes [85]. We excluded bases with mapping base quality scores less than 30, assumed a prior heterozygosity of 0.001, applied the aggressive PCR indel model, and only called variants with a minimum confidence threshold of 50. We used the intermediate g.vcf file approach followed by joint variant calling with the `GenotypeGVCFs` command when calling variants. We then filtered the initial set of variants to include only SNPs with an average minimum coverage of 6*×*, maximum absolute values of 2.5 for the base quality rank sum test, the mapping quality rank sum test and the read position rank sum test, a minimum ratio of variant confidence to non-reference read depth of 2.5, and a minimum mapping quality of 30. We also excluded all indels and SNPs with more than two alleles, and all SNPs on smaller scaffolds not assigned to linkage groups. Finally, SNPs with exceptionally high coverage (*>*450*×*) or that were clustered together (within 5 bps of each other) were dropped. This left us with 2,054,096 SNPs for phylogenomic analyses.

We first estimated a genome consensus or “species” tree from a concatenated alignment of the autosomal SNPs (i.e., this analysis did not include SNPs on the Z chromosome). We estimated the tree with `RAxML` (version 8.2.9) [86] under the GTR model with no rate heterogeneity. 100 bootstrap replicates were generated and analyzed to assess confidence in the bifurcations in the estimated phylogeny. Next, to examine variation in (unrooted) tree topologies across the genome, we split the alignment into non-overlapping 1000 SNP windows. We estimated phylogenies for each window using `RAxML` as described above. We then used the `R` packages `ape` (version 5.2) [87] and `phytools` (version 0.6.60) [88] to identify trees (across windows) with the same topology and to quantify the proportion/number of trees on each linkage group with each topology. We used randomization tests to determine whether autosomes, the Z chromosome, or the 49 candidate barrier loci (i.e., regions containing the 49 AIMs with restricted introgression in Dubois and high *L. idas* ancestry frequencies in Jackson Hole) exhibited a significant excess or deficit of a given topology. This was done in `R` as well. We used 1000 randomizations (permutations) of tree topologies across 1000 SNP windows (and thus linkage groups) for each analysis. Finally, we tested for correlations between the frequency of specific topologies (i.e., an “introgression” tree and tree uniting the ancient hybrid lineages) and chromosome size.

Variation in the frequency of different topologies across the genome, and particularly in the prevalence of the species-tree topology, can arise from introgression or incomplete lineage sorting with differences in effective population sizes across chromosomes (e.g., [25]). Thus, as an explicit test for reduced introgression on the Z chromosome, we calculated *f_d_* admixture proportions for each linkage group [48]. We assumed the species topology for this calculations was (*L. idas*,*L. melissa*),(H,*L. anna*), where H denotes either the Sierra Nevada or Warner mountain lineage. *f_d_* was estimated following [48] using our own script written in `R`. We then tested for correlations between admixture at the chromosome level (*f_d_*) and chromosome size. Finally, we asked whether the set of 1000 SNP windows containing the the 49 candidate barrier loci showed less evidence of admixture than expected by chance. For this, we calculated *f_d_* for the set of 49, 1000 SNP windows and compared this to a null distribution generated by repeatedly sampling 49, 1000 bp windoes at random and computed *f_d_* for the sampled set of windows. This was done in `R`.

## Acknowledgements

Thanks to Amy Springer and Megan Brady for their help in the field. We are also grateful for comments on earlier drafts of this manuscript from Jeff Feder, Patrik Nosil, Simon Martin and two anonymous reviewers. This work was funded by the National Science Foundation (DEB-1638768 to ZG, DEB-1050355 to CCN, DEB-1050149 to CAB), Utah State University and the Ecology Center at Utah State University (Ecology Center Graduate Student Award to SC). The support and resources from the Center for High Performance Computing at the University of Utah are gratefully acknowledged.

## Author contributions

S.C. generated and analyzed the GBS data, and wrote the manuscript. L.K.L. collected samples, generated some of the GBS data, and edited the manuscript. C.A.B. collected samples and edited the manuscript. J.A.F. collected samples and edited the manuscript. M.L.F. collected samples and edited the manuscript. C.C.N. collected samples, generated some of the GBS data, generated the whole genome sequence data, and edited the manuscript. Z.G. collected samples, generated and analyzed the GBS data, generated and analyzed the whole genome sequence data, and wrote the manuscript.

## Data availability

DNA sequence data have been archived on NCBIs SRA (PRJNA577236). Computer code and other key input files are available on DRYAD (accession number pending). The source data underlying Figs. 2-6 are provided as a Source Data file. Correspondence for materials (data, scripts, or samples) should be addressed to Zachariah Gompert.

## Supplemental Methods and Results

### Patterns of isolation-by-distance and taxon

We quantified patterns of gene flow and isolation-by-distance among the *L. idas*, *L. melissa* and Jackson Hole *Lycaeides* populations. We especially wanted to know whether the whole system (excluding Dubois) was well described by a simple isolation-by-distance model, as might be expected for primary divergence with limited dispersal within a single taxon, or alternatively if there was evidence of restricted gene flow among these three entities, consistent with past work and with the hypothesis of secondary contact and admixture in the Jackson Hole area. We used two complementary approaches to address this question. First, we estimated relative effective migration rates among the populations based on a population allele frequency covariance matrix, which we calculated from all 39,193 SNPs. This was done using the program `eems` (version 0.0.0.9000) [89]. This method does not aim to estimate absolute migration rates, but rather identifies regions in space with low or high gene flow relative to a simple two-dimensional stepping-stone isolation-by-distance model [89]. Based on past work [29, 30, 28], we expected higher effective gene flow among conspecific populations, and lower effective gene flow between species and especially in the Jackson Hole admixture area. We assumed 300 demes on a triangular grid, and fit the model using MCMC with three chains, 6 million sampling iterations, 3 million burnin iterations, and a thinning interval of 10,000. As predicted, higher effective gene flow was inferred along paths connecting conspecific populations, and a region of low effective migration (i.e., a barrier top gene flow) was observed in the Jackson Hole *Lycaeides* region (Fig. S3). However, evidence of a barrier was not evident for paths between different Jackson Hole populations within this region.

Next, we fit Bayesian linear mixed models to assess the relative contributions of geographic distance and differences in nominal taxon (*L. idas*, *L. melissa* or Jackson Hole *Lycaeides*) in explaining patterns of genetic differentiation among populations (as described in [28]). This Bayesian regression analysis is based on the mixed model framework proposed by [90] to account for the correlated error structure inherent in pairwise observations such as genetic distances. We fit a model for logit transformed F_ST_ (defined as 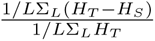, where *H_S_* and *H_T_* are the expected heterozygosities for the individual and combined population pairs) as a function of log geographic distance (great circle distance) and taxon distance (defined as 0 for populations from the same nominal taxon and 1 for populations from different nominal taxa). We fit the Bayesian model using MCMC via the `rjags` interface with `JAGS`. We placed minimally informative priors on the covariates, normal(*µ* = 0*, τ* = 0.001), and on the population random effects and residual errors, both gamma(1, 0.01). We compared the full models to models with only geographic or taxon distance using deviance information criterion (DIC). We ran three chains for each model, each with 10,000 iterations, a 2000 iteration burnin, and a thinning interval of 5. A model including geographic distance and same vs. different taxon was preferred (DIC = 92.82) relative to one with only taxon (DIC = 98.67) or only geographic distance (DIC = 360.9) (Fig. S4). Thus, whereas there is a pattern of isolation-by-distance within taxa, genetic distances between heterospecific taxa (including Jackson Hole vs. *L. idas* or *L. melissa*) are higher than expected from geographic distances along (Fig. S4). Taken together, these results show that structure exists within all of these entities, but that the system is not well-described by a simple isolation-by-distance model.

## Supplemental Tables and Figures

**Table S1:** Locality information and sample sizes (*N*) for the populations included in this study. * denote new sequence data generated for this study, other sequence data were previously presented in [28].

| Locality | ID | Taxon | Longitude | Latitude | N |
| --- | --- | --- | --- | --- | --- |
| Bonneville, UT | BST | *L. melissa* | -111.7939 | 41.7250 | *24 |
| Cody, WY | CDY | *L. melissa* | -108.9780 | 44.5121 | 23 |
| Cokeville, WY | CKV | *L. melissa* | -110.9373 | 42.0052 | 10 |
| Lander, WY | LAN | *L. melissa* | -108.3551 | 42.6533 | 24 |
| Montague, CA | MON | *L. melissa* | -107.8162 | 38.3746 | 20 |
| Yellow Pine, WY | YWP | *L. melissa* | -105.4006 | 41.2519 | 20 |
| Sinclair, WY | SIN | *L. melissa* | -107.1131 | 41.8511 | 97 |
| Victor, ID | VIC | *L. melissa* | -111.1114 | 43.6590 | 20 |
| Bunsen Peak, WY | BNP | *L. idas* | -110.7212 | 44.9337 | 20 |
| Garnet Peak, MT | GNP | *L. idas* | -111.2245 | 45.4323 | 98 |
| King's Hill, MT | KHL | *L. idas* | -110.6990 | 46.8407 | 18 |
| Soldier Creek, MT | SDC | *L. idas* | -114.6119 | 47.2062 | 20 |
| Siyeh Creek, MT | SYV | *L. idas* | -113.6681 | 48.7026 | 20 |
| Bald Mountain, WY | BLD | Jackson Hole | -109.7161 | 43.5225 | *74 |
| Bull Creek, WY | BCR | Jackson Hole | -110.5530 | 43.0076 | 46 |
| Big Ice Cave, WY | BIC | Jackson Hole | -108.4012 | 45.1617 | 18 |
| Blacktail Butte, WY | BTB | Jackson Hole | -110.6820 | 43.6382 | 46 |
| Frontier Creek, WY | FRC | Jackson Hole | -109.5776 | 43.7206 | *20 |
| Pinnacle, WY | PIN | Jackson Hole | -109.9762 | 43.7401 | 20 |
| Periodic Springs, WY | PSP | Jackson Hole | -110.8493 | 42.7468 | 20 |
| Riddle Lake, WY | RDL | Jackson Hole | -110.5465 | 44.3617 | 30 |
| Rendezvous Mtn., WY | RNV | Jackson Hole | -110.8847 | 43.5957 | 32 |
| Dubois, WY | DBS | Hybrid zone | -109.6991 | 43.5623 | *115 |

**Table S2:** Summary of the observed and expected number of AIMs with the highest *L. idas* ancestry frequencies in Jackson Hole *Lycaeides* (top 10%) that were on the Z sex chromosome. Results are shown for each Jackson Hole population. We report the observed number, and x-fold enrichment and *P* value from randomization tests. Values of *P* ≤ 0.05 are in bold font.

| Locality | No. observed | x-fold enrichment | *P* |
| --- | --- | --- | --- |
| BCR | 29 | 1.28 | 0.08 |
| BIC | 75 | 3.32 | < **0.01** |
| BLD | 66 | 2.92 | < **0.01** |
| BTB | 58 | 2.57 | < **0.01** |
| FRC | 65 | 2.86 | < **0.01** |
| PIN | 64 | 2.83 | < **0.01** |
| PSP | 70 | 3.11 | < **0.01** |
| RDL | 52 | 2.30 | < **0.01** |
| RNV | 60 | 2.65 | < **0.01** |

**Table S3:** Table shows summary of randomization tests for presence of top 10% SNPs showing excess *L. melissa* ancestry frequency on Z chromosome for 10 Jackson Hole-*Lycaeides* localities (x-fold = Number of SNPs observed is how much more than chance; *P* = randomization-based *P*-values for the null hypothesis that the proportion of top SNPs observed is not greater than the genomic proportion). *P* ≤ 0.05 are in bold.

| Locality | No. observed | x-fold enrichment | *P* |
| --- | --- | --- | --- |
| BCR | 36 | 1.59 | <**0.01** |
| BIC | 9 | 0.39 | 0.99 |
| BLD | 8 | 0.35 | 1.00 |
| BTB | 13 | 0.57 | 0.99 |
| FRC | 6 | 0.26 | 1.00 |
| PIN | 12 | 0.53 | 0.99 |
| PSP | 12 | 0.52 | 0.98 |
| RDL | 15 | 0.66 | 0.98 |
| RNV | 15 | 0.65 | 0.99 |

**Table S4:** Summary of the number of AIMs with the highest *L. idas* ancestry frequencies in Jackson Hole *Lycaeides* (top 10%) in or near (with 1000 bp) genes (this includes all of the sequence elements necessary to encode a functional transcript), coding sequences (including only sequences between start and stop codons; CDS), transposable elements (TEs) and proteins (matches against protein sequences). We report the observed number, and x-fold enrichment and *P* value from randomization tests. Values of *P* ≤ 0.05 are in bold font.

| Category | No. observed | x-fold enrichment | $P$ |
| --- | --- | --- | --- |
| on gene | 52 | 1.17 | 0.07 |
| near gene | 64 | 1.22 | **0.01** |
| on CDS | 24 | 1.44 | **0.03** |
| near CDS | 56 | 1.25 | **0.02** |
| on TE | 3 | 0.90 | 0.67 |
| near TE | 14 | 0.99 | 0.57 |
| on protein | 62 | 1.06 | 0.28 |
| near protein | 87 | 1.18 | **<0.00** |

**Table S5:** Summary of the number of AIMs with the highest *L. melissa* ancestry frequencies in Jackson Hole *Lycaeides* (top 10%) in or near (with 1000 bp) genes (this includes all of the sequence elements necessary to encode a functional transcript), coding sequences (including only sequences between start and stop codons; CDS), transposable elements (TEs) and proteins (matches against protein sequences). We report the observed number, and x-fold enrichment and *P* value from randomization tests. Values of *P* ≤ 0.05 are in bold font.

| Category | No. observed | x-fold enrichment | $P$ |
| --- | --- | --- | --- |
| on gene | 55 | 1.24 | **0.02** |
| near gene | 61 | 1.16 | 0.06 |
| on CDS | 18 | 1.08 | 0.41 |
| near CDS | 52 | 1.16 | 0.09 |
| on TE | 3 | 0.91 | 0.66 |
| near TE | 9 | 0.63 | 0.96 |
| on protein | 70 | 1.20 | **0.02** |
| near protein | 79 | 1.07 | 0.16 |

**Table S6:** Summary of the number of AIMs with credible evidence of excess directional introgression of Jackson Hole *Lycaeides* alleles in the Dubois hybrid zone (*α* > 0) in or near (with 1000 bp) genes (this includes all of the sequence elements necessary to encode a functional transcript), coding sequences (including only sequences between start and stop codons; CDS), transposable elements (TEs) and proteins (matches against protein sequences). We report the observed number, and x-fold enrichment and *P* value from randomization tests. Values of *P* ≤ 0.05 are in bold font.

| Category | No. observed | x-fold enrichment | $P$ |
| --- | --- | --- | --- |
| on gene | 79 | 1.10 | 0.13 |
| near gene | 93 | 1.10 | 0.10 |
| on cds | 29 | 1.08 | 0.36 |
| near cds | 78 | 1.07 | 0.21 |
| on TE | 5 | 0.94 | 0.64 |
| near TE | 24 | 1.05 | 0.44 |
| on protein | 98 | 1.04 | 0.30 |
| near protein | 124 | 1.04 | 0.23 |

**Table S7:**
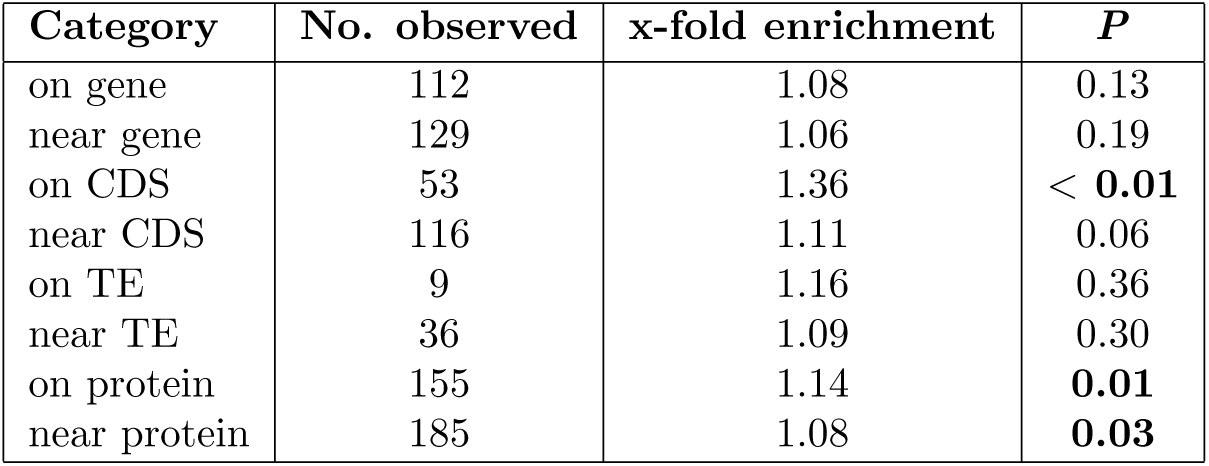
Summary of the number of AIMs with credible evidence of excess directional introgression of *L. melissa* alleles in the Dubois hybrid zone (*α* < 0) in or near (with 1000 bp) genes (this includes all of the sequence elements necessary to encode a functional transcript), coding sequences (including only sequences between start and stop codons; CDS), transposable elements (TEs) and proteins (matches against protein sequences). We report the observed number, and x-fold enrichment and *P* value from randomization tests. Values of *P* ≤ 0.05 are in bold font.

| Category | No. observed | x-fold enrichment | $P$ |
| --- | --- | --- | --- |
| on gene | 112 | 1.08 | 0.13 |
| near gene | 129 | 1.06 | 0.19 |
| on CDS | 53 | 1.36 | **< 0.01** |
| near CDS | 116 | 1.11 | 0.06 |
| on TE | 9 | 1.16 | 0.36 |
| near TE | 36 | 1.09 | 0.30 |
| on protein | 155 | 1.14 | **0.01** |
| near protein | 185 | 1.08 | **0.03** |

**Table S8:**
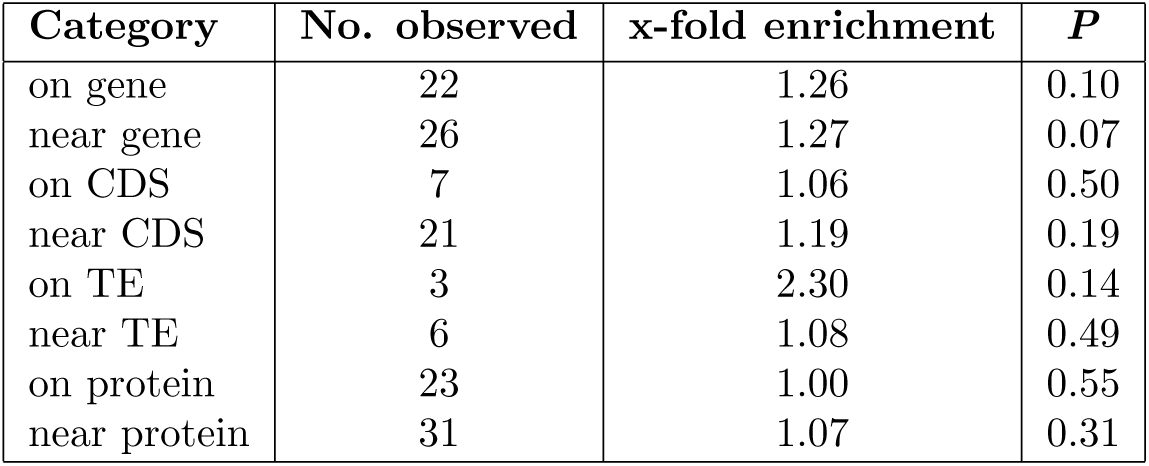
Summary of the number of AIMs with credible evidence of restricted introgression in the Dubois hybrid zone (*β* > 0) in or near (with 1000 bp) genes (this includes all of the sequence elements necessary to encode a functional transcript), coding sequences (including only sequences between start and stop codons; CDS), transposable elements (TEs) and proteins (matches against protein sequences). We report the observed number, and x-fold enrichment and *P* value from randomization tests. Values of *P* ≤ 0.05 are in bold font.

**Table S9:** Summary of the number of AIMs with restricted introgression in the Dubois hybrid zone (10% with the highest estimates of *β* from `bgc`) and with extreme (high *L. idas* ancestry frequencies in Jackson Hole (top 10%) that were in or near (with 1000 bp) genes (this includes all of the sequence elements necessary to encode a functional transcript), coding sequences (including only sequences between start and stop codons; CDS), transposable elements (TEs) and proteins (matches against protein sequences). We report the observed number, and x-fold enrichment and *P* value from randomization tests. Values of *P* ≤ 0.05 are in bold font.

| Category | No. observed | x-fold enrichment | $P$ |
| --- | --- | --- | --- |
| on gene | 25 | 1.30 | 0.06 |
| near gene | 29 | 1.27 | **0.05** |
| on CDS | 9 | 1.25 | 0.28 |
| near CDS | 23 | 1.17 | 0.20 |
| on TE | 3 | 2.07 | 0.17 |
| near TE | 6 | 0.97 | 0.60 |
| on protein | 26 | 1.02 | 0.49 |
| near protein | 34 | 1.06 | 0.33 |

**Table S10:** Summary of the number of AIMs with directional introgression of Jackson Hole alleles in the Dubois hybrid zone (10% with the highest estimates of *α* from `bgc`) and with high *L. idas* ancestry frequencies in Jackson Hole (top 10%) that were in or near (with 1000 bp) genes (this includes all of the sequence elements necessary to encode a functional transcript), coding sequences (including only sequences between start and stop codons; CDS), transposable elements (TEs) and proteins (matches against protein sequences). We report the observed number, and x-fold enrichment and *P* value from randomization tests. Values of *P* ≤ 0.05 are in bold font.

| Category | No. observed | x-fold enrichment | $P$ |
| --- | --- | --- | --- |
| on gene | 9 | 1.08 | 0.46 |
| near gene | 12 | 1.22 | 0.24 |
| on CDS | 5 | 1.59 | 0.20 |
| near CDS | 11 | 1.30 | 0.18 |
| on TE | 0 | 0.00 | 1.00 |
| near TE | 2 | 0.75 | 0.77 |
| on protein | 12 | 1.10 | 0.41 |
| near protein | 16 | 1.16 | 0.23 |

**Table S11:** Summary of the number of AIMs with directional introgression of Jackson Hole alleles in the Dubois hybrid zone (10% with the highest estimates of *α* from `bgc`) and with high *L. melissa* ancestry frequencies in Jackson Hole (top 10%) that were in or near (with 1000 bp) genes (this includes all of the sequence elements necessary to encode a functional transcript), coding sequences (including only sequences between start and stop codons; CDS), transposable elements (TEs) and proteins (matches against protein sequences). We report the observed number, and x-fold enrichment and *P* value from randomization tests. Values of *P* ≤ 0.05 are in bold font.

| Category | No. observed | x-fold enrichment | $P$ |
| --- | --- | --- | --- |
| on gene | 14 | 1.38 | 0.09 |
| near gene | 15 | 1.24 | 0.17 |
| on CDS | 6 | 1.56 | 0.18 |
| near CDS | 15 | 1.45 | **0.05** |
| on TE | 1 | 1.30 | 0.55 |
| near TE | 2 | 0.61 | 0.86 |
| on protein | 15 | 1.12 | 0.34 |
| near protein | 17 | 1.00 | 0.59 |

**Table S12:**
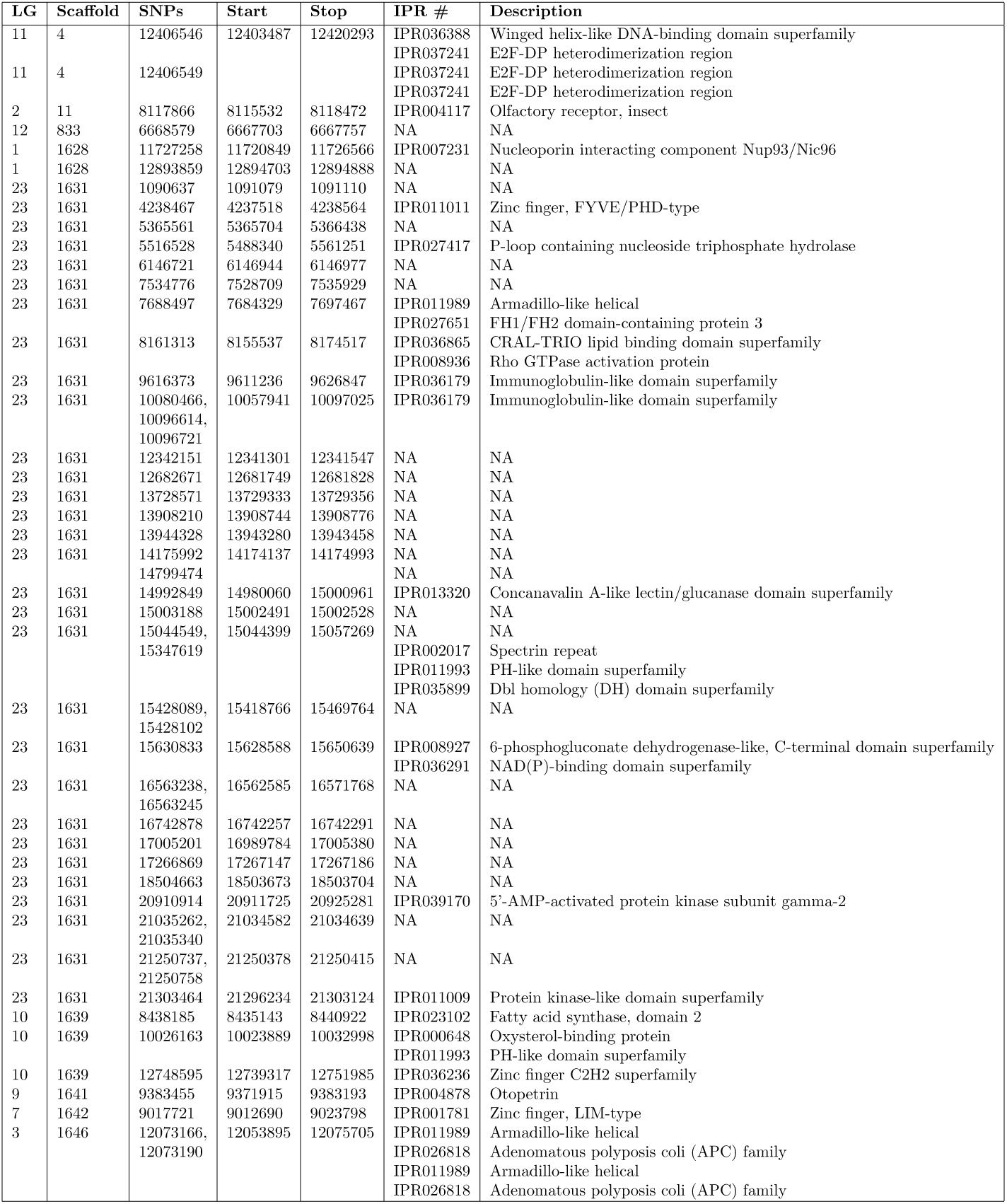
Summary of protein descriptions from `interproscan` for AIMs which had high *L. idas* in the ancient, Jackson Hole hybrids and restricted introgression in the contemporary Dubois hybrid zone (top 10% in each case). For each gene, the linkage group (LG) (LG 23 = Z), genome scaffold number, positions of SNP or SNPs in or near the gene (SNPs) and start and stop of the gene annotation are given along with the IPR number and associated description from `interproscan`.

**Table S13:**
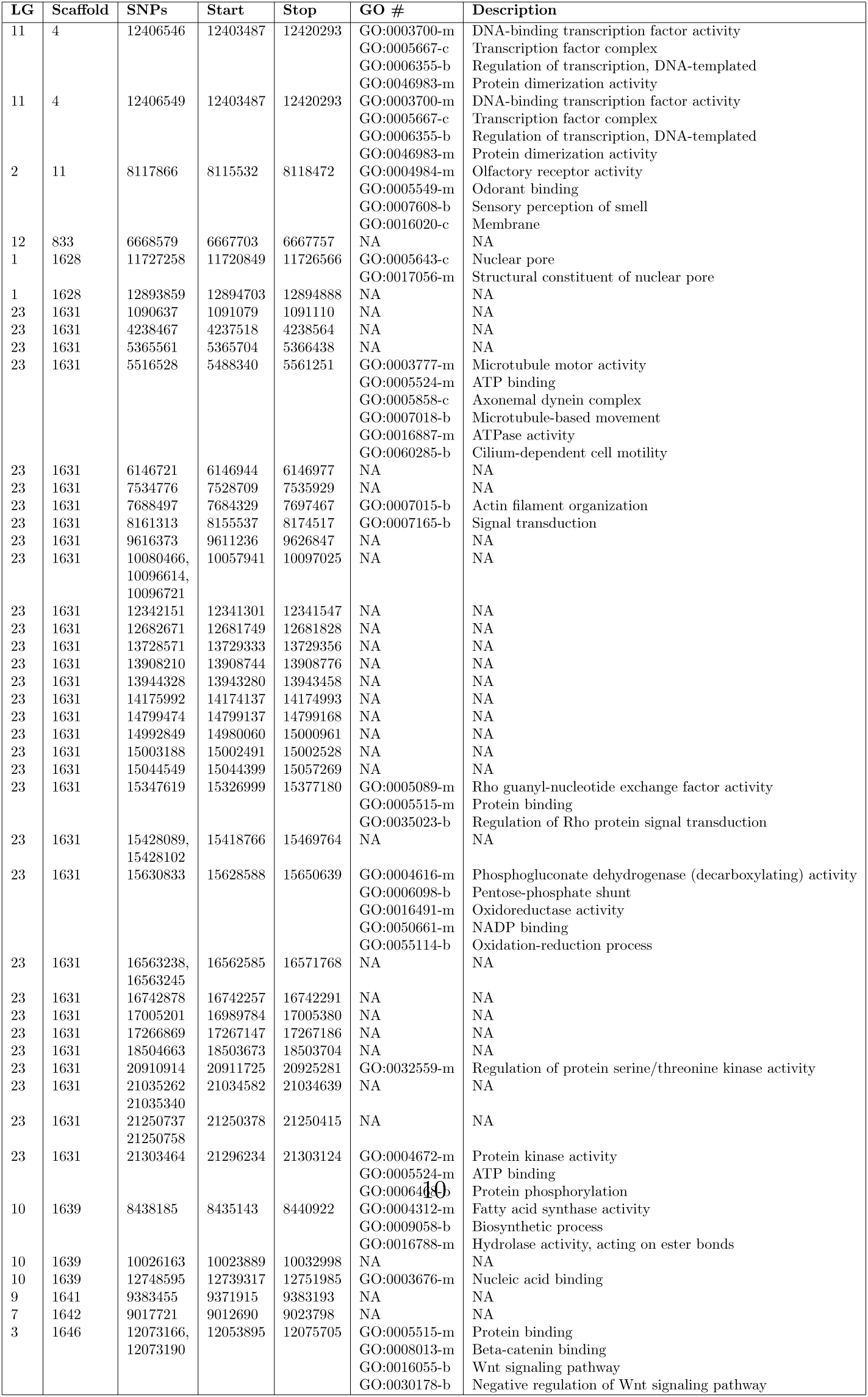
Summary of gene ontology (GO) terms (i.e., classifications) for AIMs which had high *L. idas* ancestry in the ancient, Jackson Hole hybrids and restricted introgression in the contemporary Dubois hybrid zone (top 10% in each case). For each gene, the linkage group (LG) (LG 23 = Z), genome scaffold number, positions of SNP or SNPs in or near the gene (SNPs) and start and stop of the gene annotation are given along with the GO term numbers (IDs) and descriptions. Symbols denote whether each term corresponds to a biological process (-b), molecular function (-m), or cellular component (-c).

**Table S14:** Summary of protein descriptions from `interproscan` for AIMs which had high *L. idas* ancestry in the ancient, Jackson Hole hybrids and directional Jackson Hole *Lycaeides* introgression (*α* > 0) in the contemporary Dubois hybrid zone (top 10% in each case). For each gene, the linkage group (LG) (LG 23 = Z), genome scaffold number, positions of SNP or SNPs in or near the gene (SNPs) and start and stop of the gene annotation are given along with the IPR number and associated description from `interproscan`.

| LG | Scaffold | SNPs | Start | Stop | IPR # | Description |
| --- | --- | --- | --- | --- | --- | --- |
| 2 | 11 | 6234637 | 6215867 | 6240330 | IPR001930 | Peptidase M1, alanine aminopeptidase/leukotriene A4 hydrolase |
|  |  |  |  |  | IPR024571 | ERAP1-like C-terminal domain |
| 2 | 11 | 15712950 | 15710210 | 15713015 | IPR009851 | Modifier of rudimentary, Modr |
| 2 | 11 | 15712950 | NA | NA | IPR037859 | Vacuolar protein sorting-associated protein 37 |
| 16 | 503 | 6014039 | NA | NA | NA | NA |
| 12 | 833 | 6668579 | 6667703 | 6667757 | NA | NA |
| 23 | 1631 | 1200324 | 1197667 | 1204058 | IPR036719 | Neurotransmitter-gated ion-channel transmembrane domain superfamily |
| 23 | 1631 | 1200324 | NA | NA | IPR036734 | Neurotransmitter-gated ion-channel ligand-binding domain superfamily |
| 23 | 1631 | 2570345 | 2570129 | 2570178 | NA | NA |
| 23 | 1631 | 4238526 | 4237518 | 4238564 | IPR011011 | Zinc finger, FYVE/PHD-type |
| 23 | 1631 | 9958857 | 9955713 | 9968188 | IPR027266 | GTP-binding protein TrmE/Glycine cleavage system T protein, domain 1 |
| 23 | 1631 | 9958857 | NA | NA | IPR036188 | FAD/NAD(P)-binding domain superfamily |
| 23 | 1631 | 10080421 | 10057941 | 10097025 | IPR036179 | Immunoglobulin-like domain superfamily |
| 23 | 1631 | 10080466 | 10057941 | 10097025 | IPR036179 | Immunoglobulin-like domain superfamily |
| 23 | 1631 | 12954191 | 12954213 | 12954434 | NA | NA |
| 23 | 1631 | 14218357 | 14216920 | 14217629 | NA | NA |
| 23 | 1631 | 14760795 | 14746596 | 14779181 | IPR000034 | Laminin IV |
|  |  |  |  |  | IPR002049 | Laminin EGF domain |
|  |  |  |  |  | IPR036055 | LDL receptor-like superfamily |
|  |  |  |  |  | IPR036179 | Immunoglobulin-like domain superfamily |
| 23 | 1631 | 15408704 | 15409126 | 15409167 | NA | NA |
| 23 | 1631 | 15428179 | 15418766 | 15469764 | NA | NA |
| 23 | 1631 | 20910914 | 20911725 | 20925281 | IPR000644 | CBS domain |
|  |  |  |  |  | IPR039170 | 5'-AMP-activated protein kinase subunit gamma-2 |
| 13 | 1632 | 6146628 | 6145294 | 6150912 | IPR011047 | Quinoprotein alcohol dehydrogenase-like superfamily |
| 10 | 1639 | 8378132 | 8374436 | 8380586 | NA | NA |
| 10 | 1639 | 13668785 | 13668901 | 13668943 | NA | NA |
| 9 | 1641 | 9384153 | 9371915 | 9383193 | IPR004878 | Otopetrin |
| 7 | 1642 | 9023853 | 9012690 | 9025104 | IPR001781 | Zinc finger, LIM-type |
| 4 | 1648 | 13426790 | 13426596 | 13426775 | NA | NA |

**Table S15:**
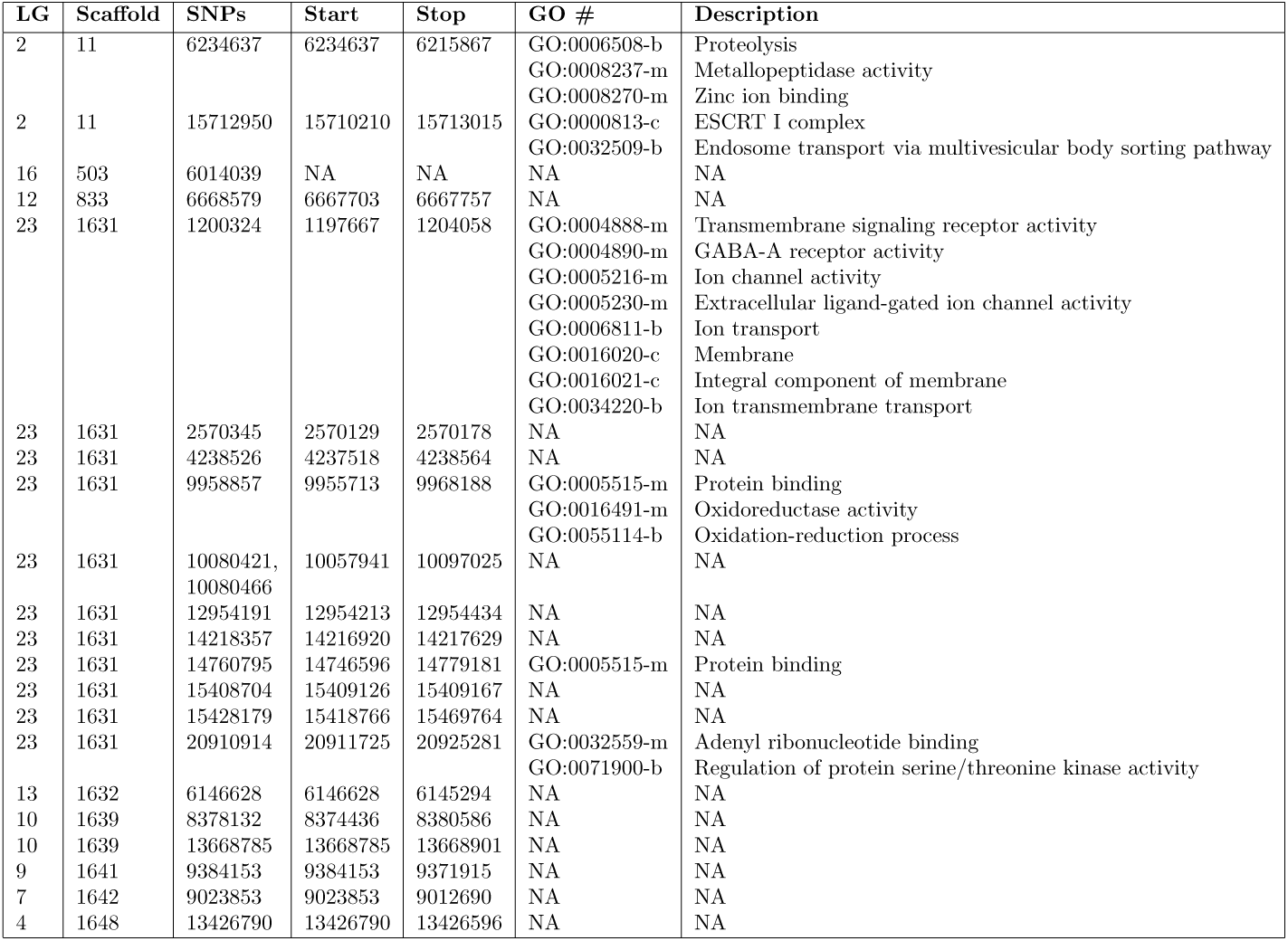
Summary of gene ontology (GO) terms (i.e., classifications) for AIMs which had high *L. idas* ancestry in the ancient, Jackson Hole hybrids and directional Jackson Hole *Lycaeides* introgression (*α* > 0) in the contemporary Dubois hybrid zone (top 10% in each case). For each gene, the linkage group (LG) (LG 23 = Z), genome scaffold number, positions of SNP or SNPs in or near the gene (SNPs) and start and stop of the gene annotation are given along with the GO term numbers (IDs) and descriptions. Symbols denote whether each term corresponds to a biological process (-b), molecular function (-m), or cellular component (-c).

**Table S16:**
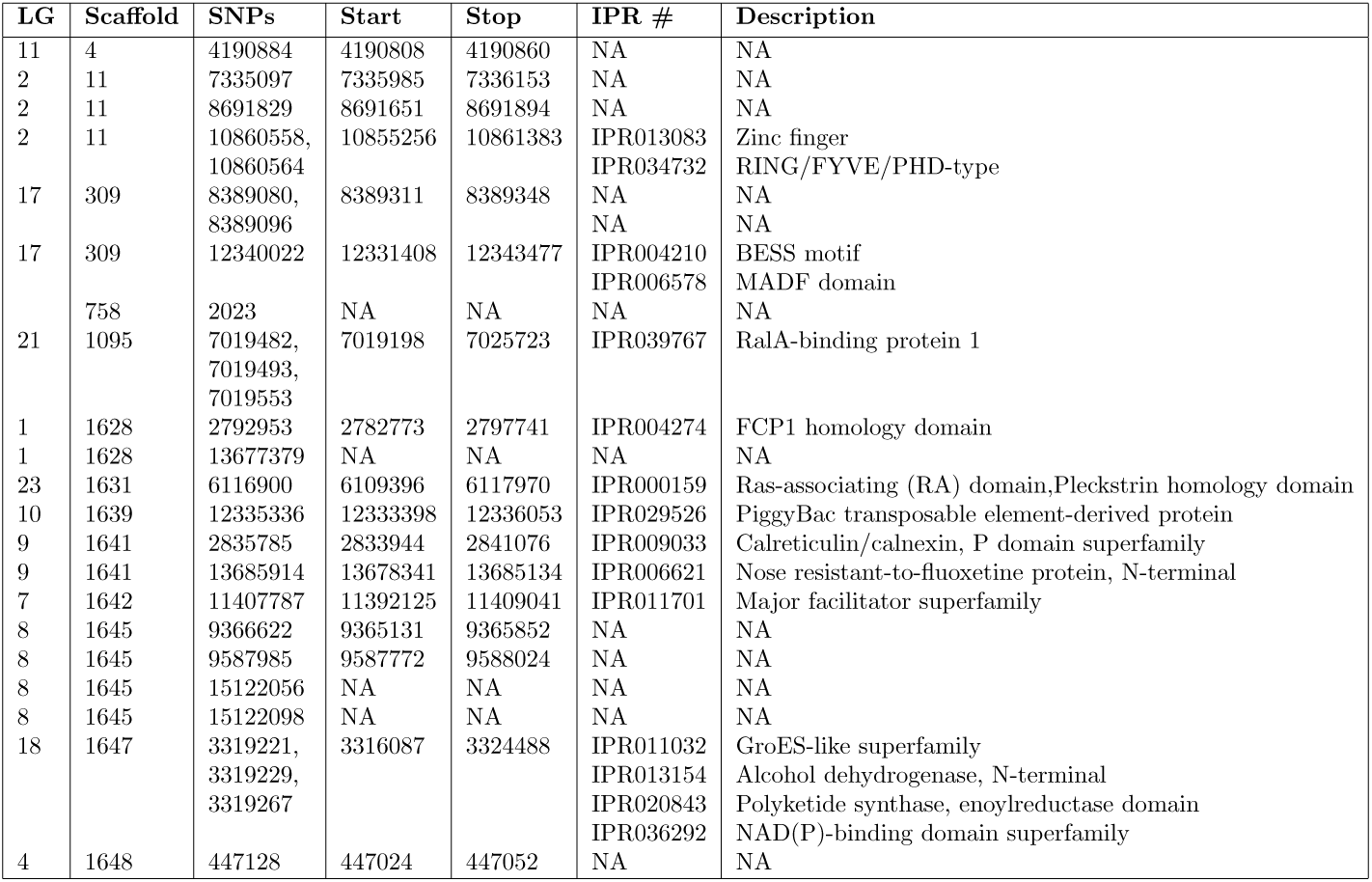
Summary of protein descriptions from `interproscan` for AIMs which had high *L. melissa* ancestry in the ancient, Jackson Hole hybrids and directional Jackson Hole *Lycaeides* introgression (*α* > 0) in the contemporary Dubois hybrid zone (top 10% in each case). For each gene, the linkage group (LG) (LG 23 = Z), genome scaffold number, positions of SNP or SNPs in or near the gene (SNPs) and start and stop of the gene annotation are given along with the IPR number and associated description from `interproscan`.

**Table S17:** Summary of gene ontology (GO) terms (i.e., classifications) for AIMs which had high *L. melissa* ancestry in the ancient, Jackson Hole hybrids and directional Jackson Hole *Lycaeides* introgression (*α* > 0) in the contemporary Dubois hybrid zone (top 10% in each case). For each gene, the linkage group (LG) (LG 23 = Z), genome scaffold number, positions of SNP or SNPs in or near the gene (SNPs) and start and stop of the gene annotation are given along with the GO term numbers (IDs) and descriptions. Symbols denote whether each term corresponds to a biological process (-b), molecular function (-m), or cellular component (-c).

| LG | Scaffold | SNPs | Start | Stop | GO # | Description |
| --- | --- | --- | --- | --- | --- | --- |
| 11 | 4 | 4190884 | 4190808 | 4190860 | NA | NA |
| 2 | 11 | 7335097 | 7335985 | 7336153 | NA | NA |
| 2 | 11 | 8691829 | 8691651 | 8691894 | NA | NA |
| 2 | 11 | 10860558,<br>10860564 | 10855256 | 10861383 | NA | NA |
| 17 | 309 | 8389080,<br>8389096 | 8389311 | 8389348 | NA | NA |
| 17 | 309 | 12340022 | 12331408 | 12343477 | GO:0003677-m | DNA binding |
|  | 758 | 2023 | NA | NA | NA | NA |
| 21 | 1095 | 7019482,<br>7019493,<br>7019553 | 7019198 | 7025723 | GO:0005096-m | GTPase activator activity |
| 1 | 1628 | 2792953 | 2782773 | 2797741 | GO:0016791-m | Phosphatase activity |
| 1 | 1628 | 13677379 | NA | NA | NA | NA |
| 23 | 1631 | 6116900 | 6109396 | 6117970 | GO:0007165-b | Signal transduction |
| 10 | 1639 | 12335336 | 12333398 | 12336053 | NA | NA |
| 9 | 1641 | 2835785 | 2833944 | 2841076 | GO:0005509 | calcium ion binding |
|  |  |  |  |  | GO:0051082 | protein folding |
| 9 | 1641 | 13685914 | 13678341 | 13685134 | NA | NA |
| 7 | 1642 | 11407787 | 11392125 | 11409041 | GO:0016021-c | Integral component of membrane |
| 8 | 1645 | 9366622 | 9365131 | 9365852 | NA | NA |
| 8 | 1645 | 9587985 | 9587772 | 9588024 | NA | NA |
| 8 | 1645 | 15122056 | NA | NA | NA | NA |
| 8 | 1645 | 15122098 | NA | NA | NA | NA |
| 18 | 1647 | 3319221,<br>3319229,<br>3319267 | 3316087 | 3324488 | GO:0016491-m | Oxidoreductase activity |
| 4 | 1648 | 447128 | 447024 | 447052 | NA | NA |

**Table S18:**
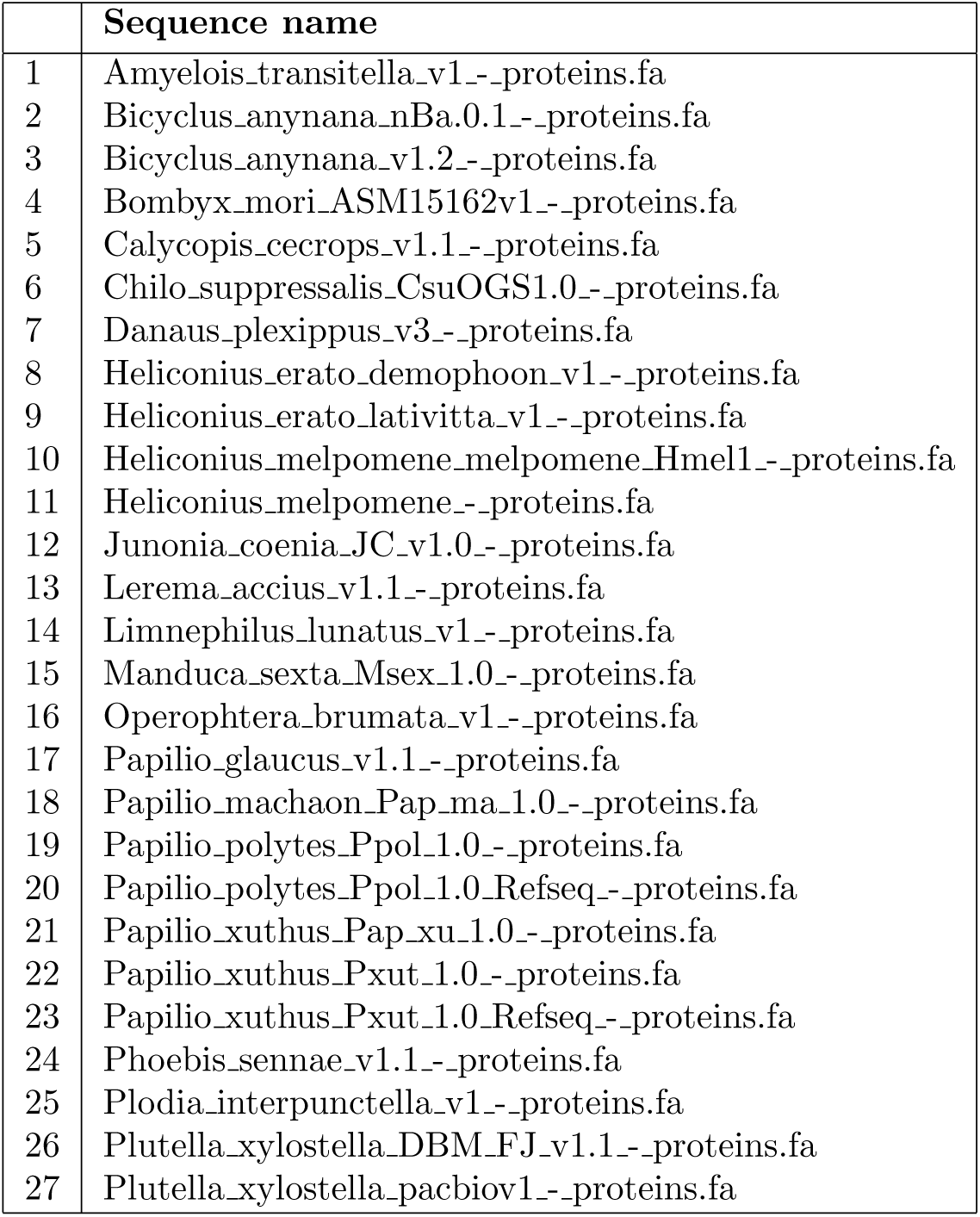
Table gives a list of sequences used from LepBase version 4 to create the protein homology file for Genome Annotation using MAKER pipeline.

**Figure S1:**
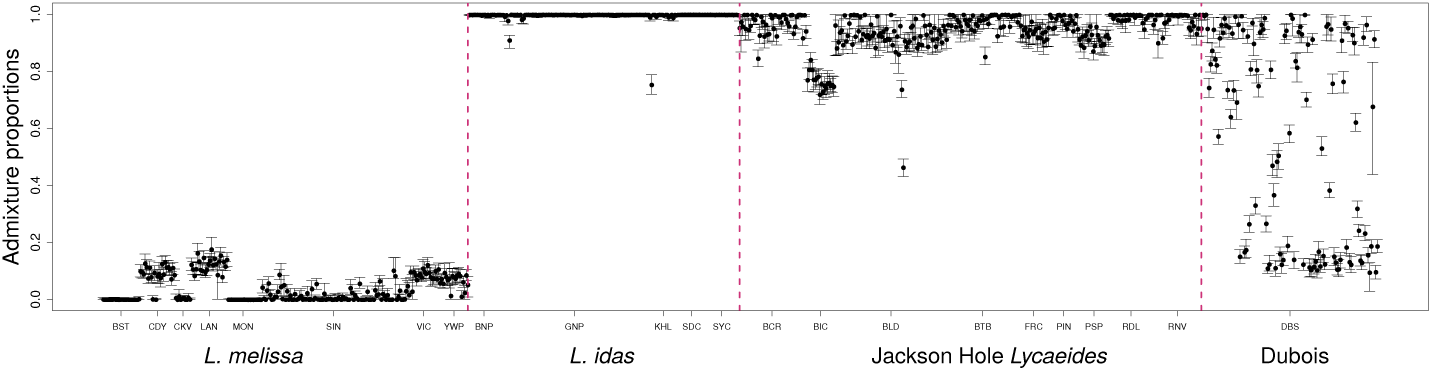
Admixture proportion estimates from `entropy` with *k* = 2 source populations. Points (posterior median) and error bars (95% credible intervals) denote the estimates of the proportion of the genome with *L. idas* ancestry. Tick marks below the plots identify populations based on the population abbreviations in Table S1. Individuals with low coverage are not shown in this figure.

**Figure S2:**
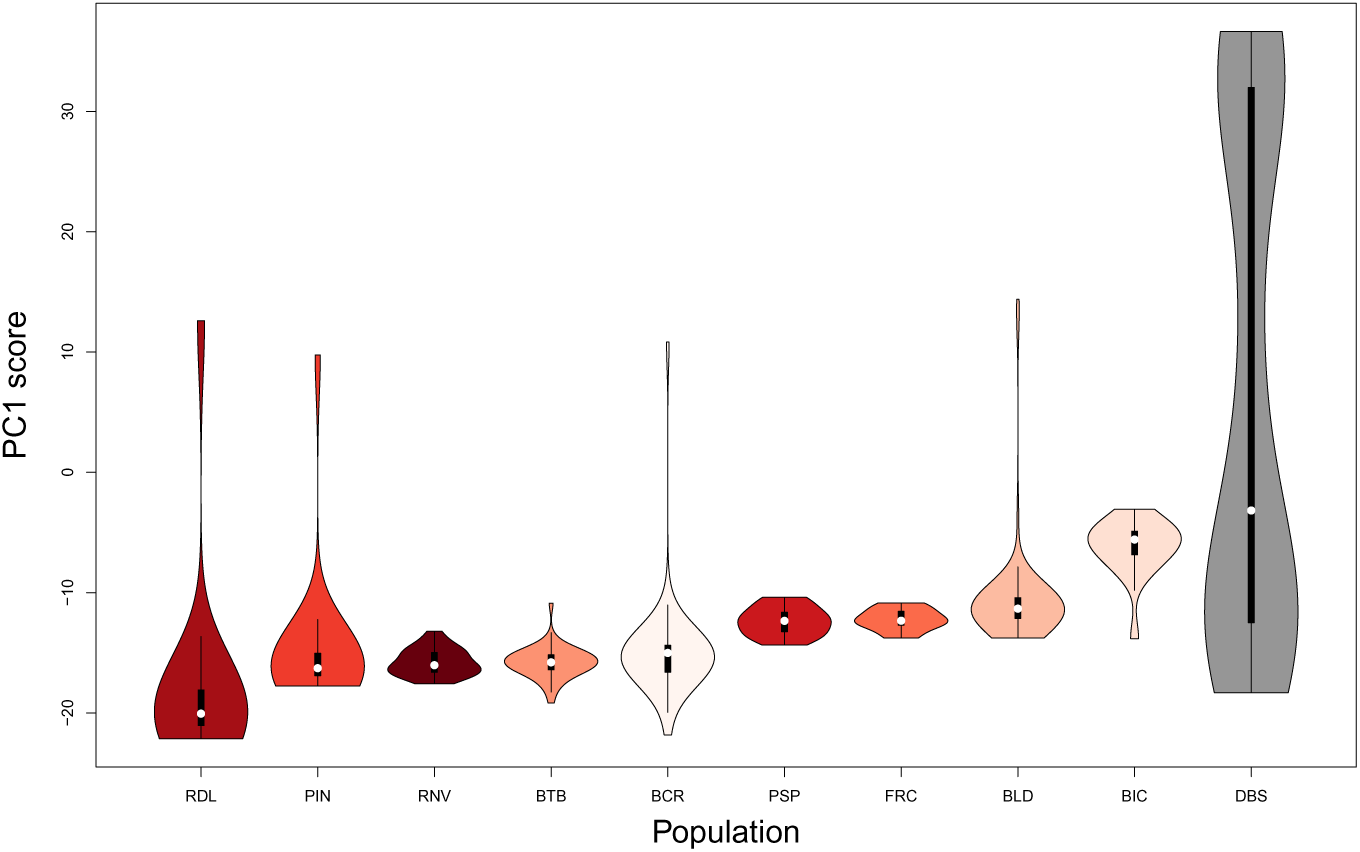
Violin plots show the distribution of genetic PC1 scores among individuals for each population. Tick marks below the plots identify populations based on the population abbreviations in Table S1.

**Figure S3:**
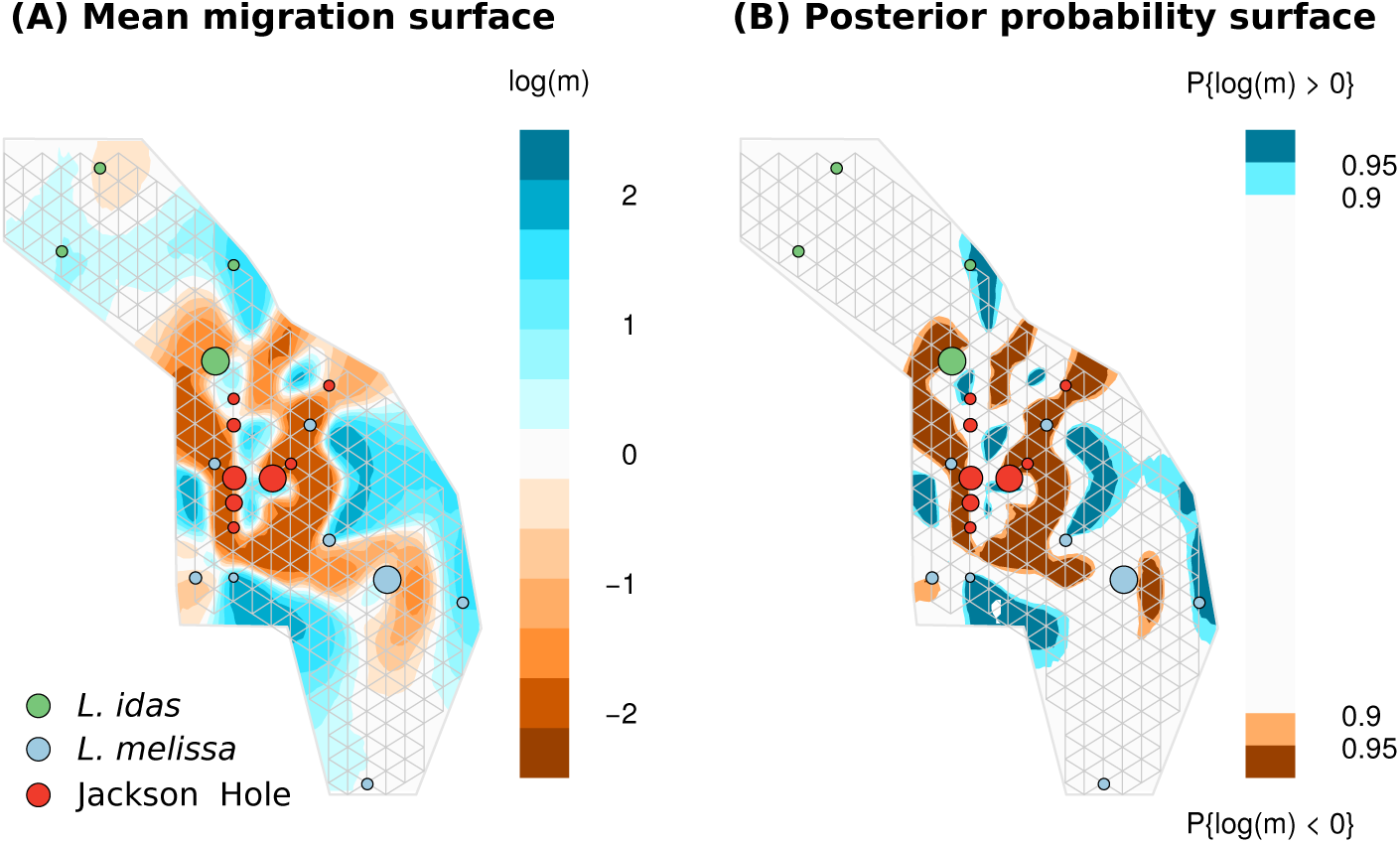
Plots summarize effective migration rates among *L. idas*, *L. melissa* and Jackson hole populations in terms of (A) the posterior mean effective migration point estimate and (B) the posterior probability that the log effecitve migration rate (*m*) is greater than or less than 0. The grid denotes 300 hypothetical demes (nodes) potentially connected by migration (edges). Point sizes are proportional to sample sizes, and individual points (demes) sometimes included individuals from two nearby populations. Dark brown regions of the map indicate reduced effective migration rates relative to expectations if gene flow were constant per unit distance across the map. This shows a barrier to gene flow centered on the Jackson Hole populations, and thus differs from a simple isolation-by-distance pattern. Dubois was not included in this analysis.

**Figure S4:**
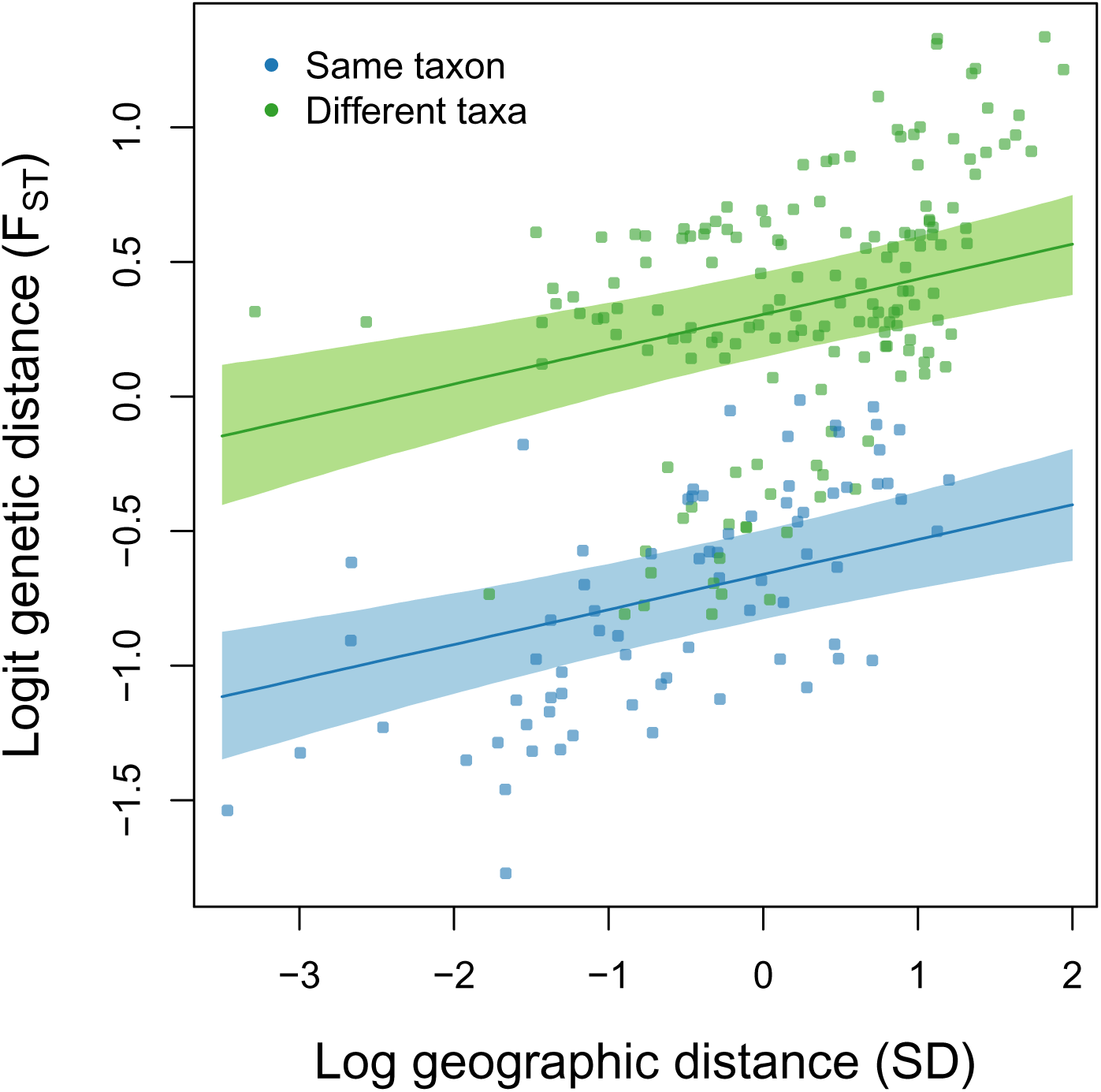
Plots shows the relationship between geographic and genetic distances among populations. We show centered logit F_ST_ as a function of standardized (centered and SD = 1) log geographic distance. Points are colored to denote whether each of the 22 population pairs (excluding Dubois) are of the same or different nominal taxa (*L. melissa*, *L. idas* or Jackson Hole). Lines and shaded areas denote Bayesian point estimates (posterior median) and 90% credible intervals for the relationship between gegoraphic and genetic distance. A model including geographic distance and same vs. different taxon was preferred (DIC = 92.82) relative to one with only taxon (DIC = 98.67) or only geographic distance (DIC = 360.9). Dubois was not included in this analysis.

**Figure S5:**
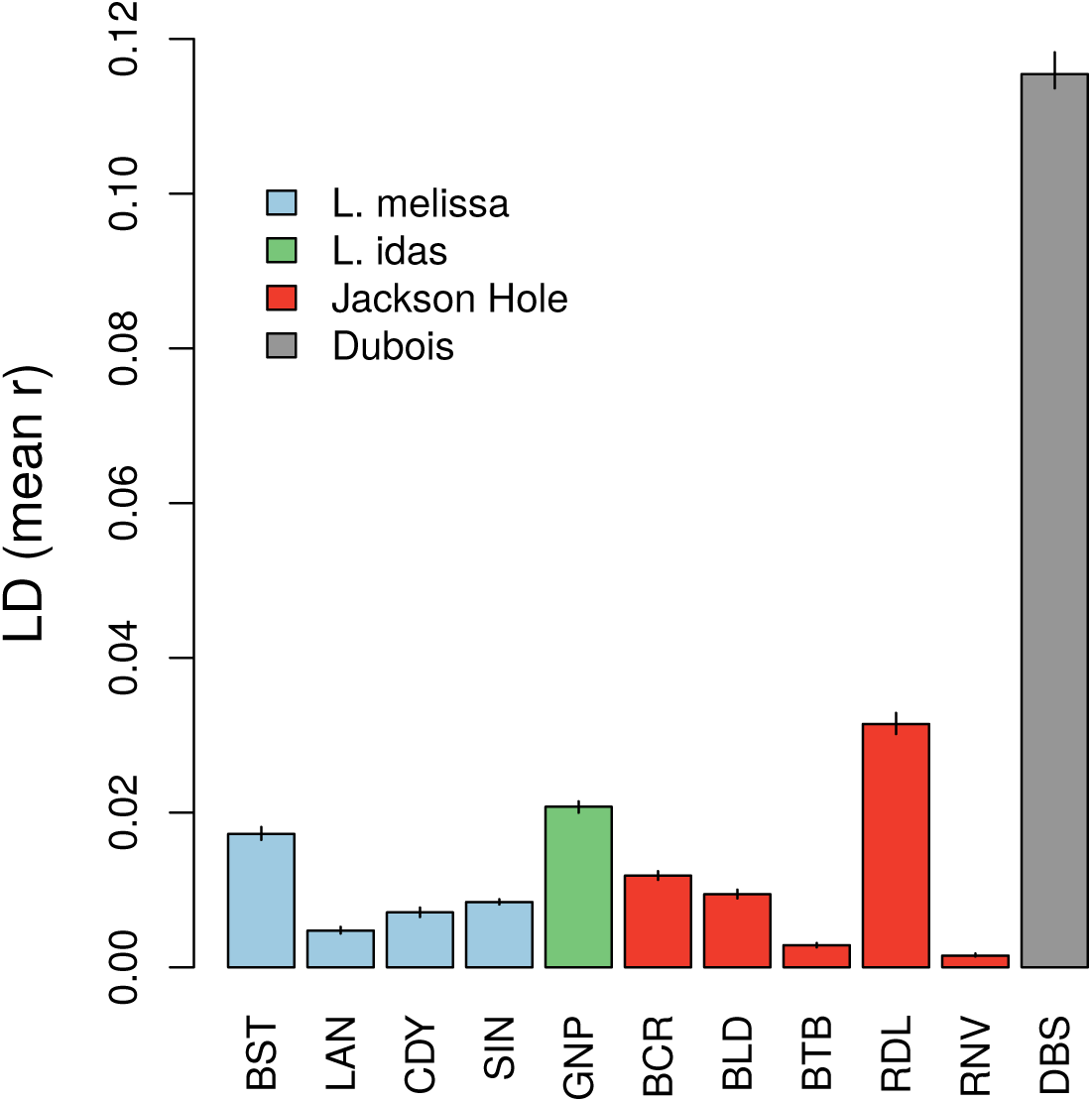
Bars denote the mean value of linkage disequilibrium (LD) for all pairs of ancestry informative SNPs (AIMs) for each population, vertical lines give *±* standard errors. Here, LD was measured as the Pearson correlation between genotypes at pairs of loci (regardless of physical linkage). LD was calculated based on posterior estimates of genotypes from `entropy`. Genotypes were first polarized such that positive LD indicates a positive association between pairs of alleles that were more common in each of the parental species (i.e., coupling LD). We report *r* rather than *r*^2^ to retain this sign information. AIMs were defined as those SNPs with an allele frequency difference of at least 0.3 between *L. idas* and *L. melissa*. Only populations with sample sizes of at least 20 were included in this analysis. Population abbreviations are defined in Table S1.

**Figure S6:**
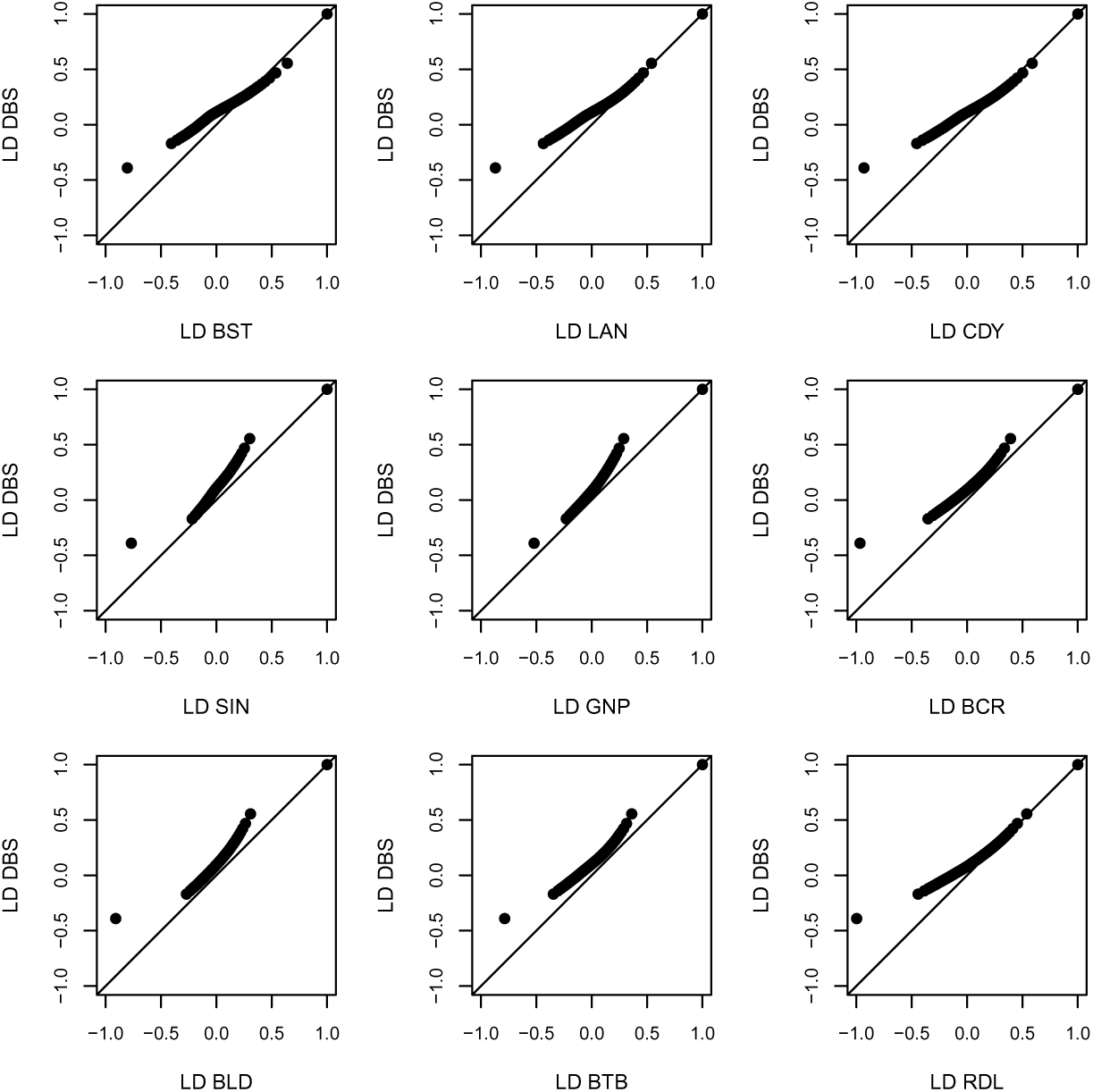
Quantile-quantile plots show the distribution of linkage disequilibrium (LD) for all pairs of ancestry informative SNPs (AIMs) for each population when compared to Dubois. Here, LD was measured as the Pearson correlation between genotypes at pairs of loci (regardless of physical linkage). LD was calculated based on posterior estimates of genotypes from `entropy`. Genotypes were first polarized such that positive LD indicates a positive association between pairs of alleles that were more common in each of the parental species (i.e., coupling LD). We report *r* rather than *r*^2^ to retain this sign information. AIMs were defined as those SNPs with an allele frequency difference of at least 0.3 between *L. idas* and *L. melissa*. Only populations with sample sizes of at least 20 were included in this analysis. Population abbreviations are defined in Table S1.

**Figure S7:**
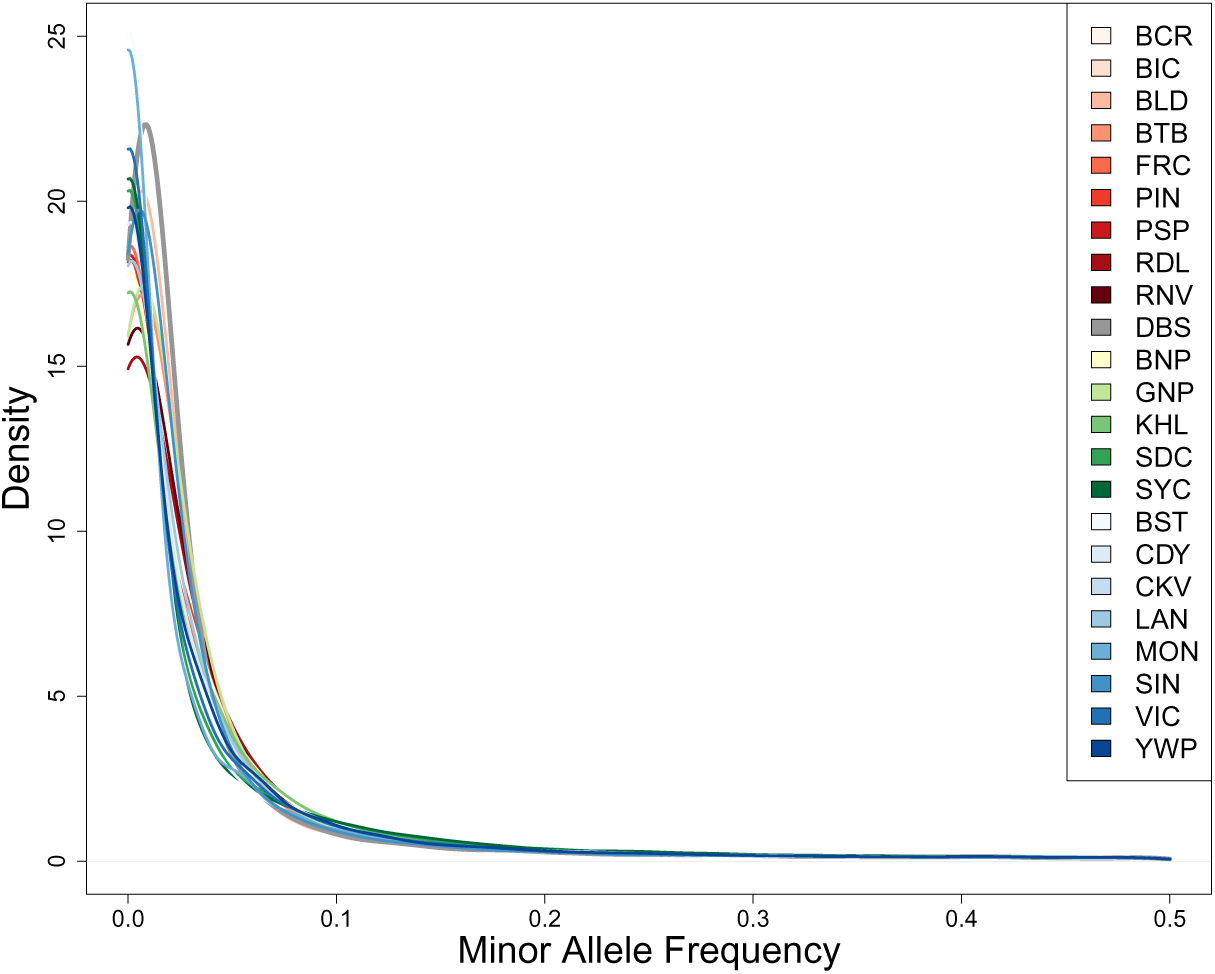
Density plot shows the estimated minor allele frequency distribution across all loci (N = 39,139 SNPs) for all populations included in this study. The population abbreviations are defined in Table S1. Mean minor allele frequencies for the subset of ancestry informative SNPs (AIMs) are showin in Fig. S8.

**Figure S8:**
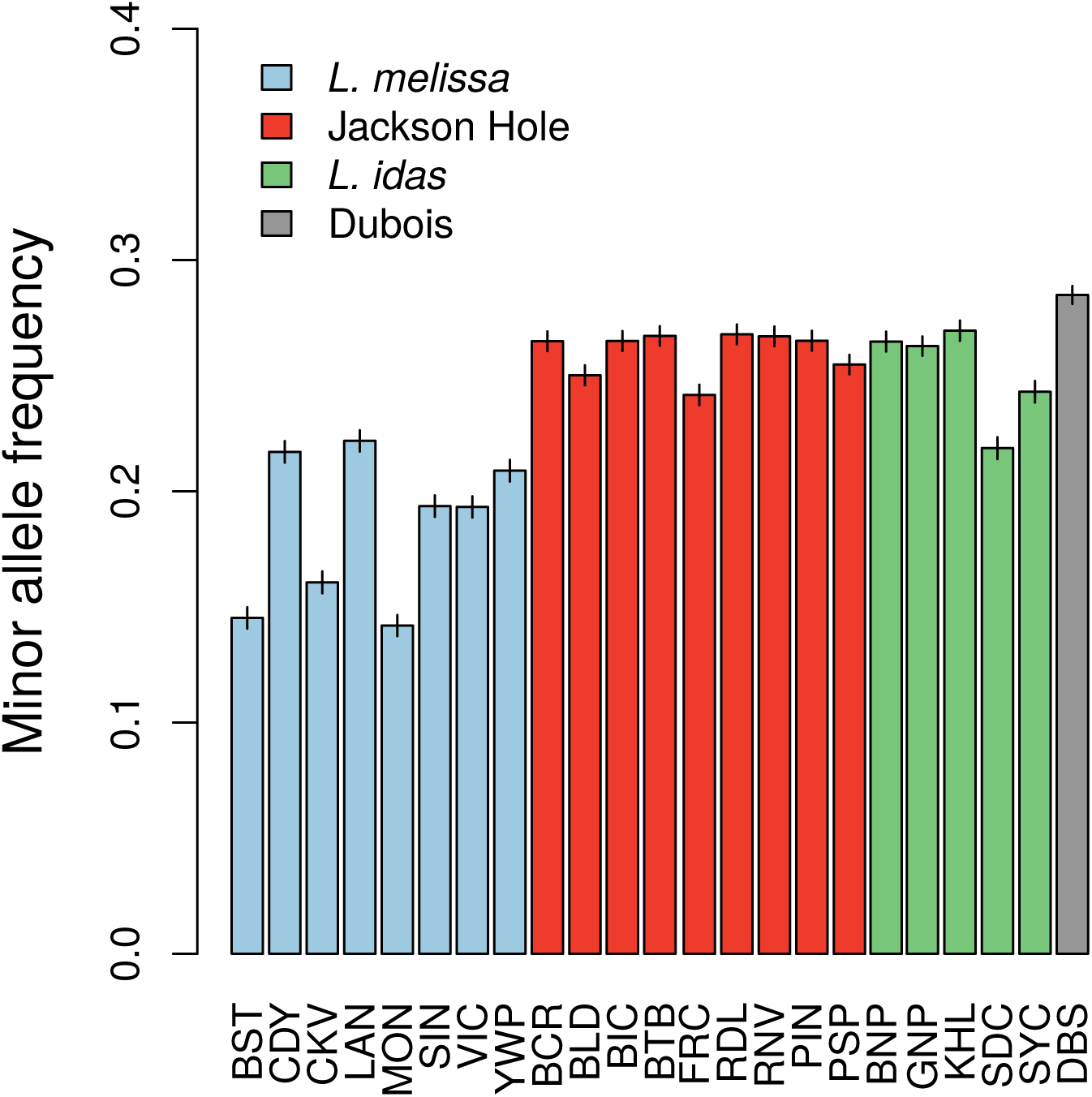
Bars show the mean estimated minor allele frequency distribution for ancestry informative SNPs (AIMs) (N = 1164) for all populations included in this study. Vertical lines give *±* standard errors for the means. Population abbreviations are defined in Table S1.

**Figure S9:**
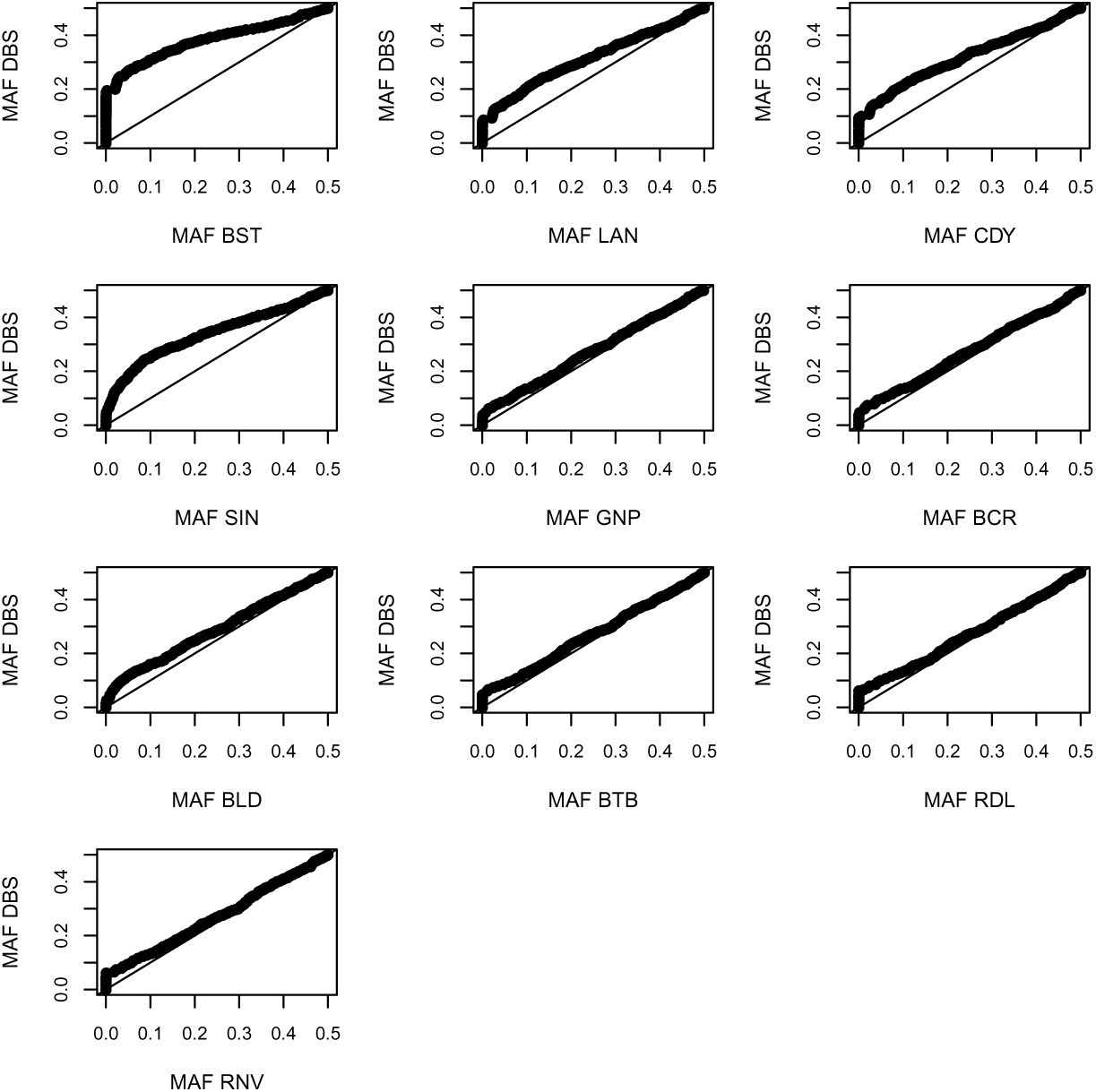
Quantile-quantile plots show the distribution of minor allele frequency (MAF) for all pairs of ancestry informative SNPs (AIMs) for each population when compared to Dubois. Here, AIMs were defined as those SNPs with an allele frequency difference of at least 0.3 between *L. idas* and *L. melissa*. Only populations with sample sizes of at least 20 were included in this analysis. Population abbreviations are defined in Table S1.

**Figure S10:**
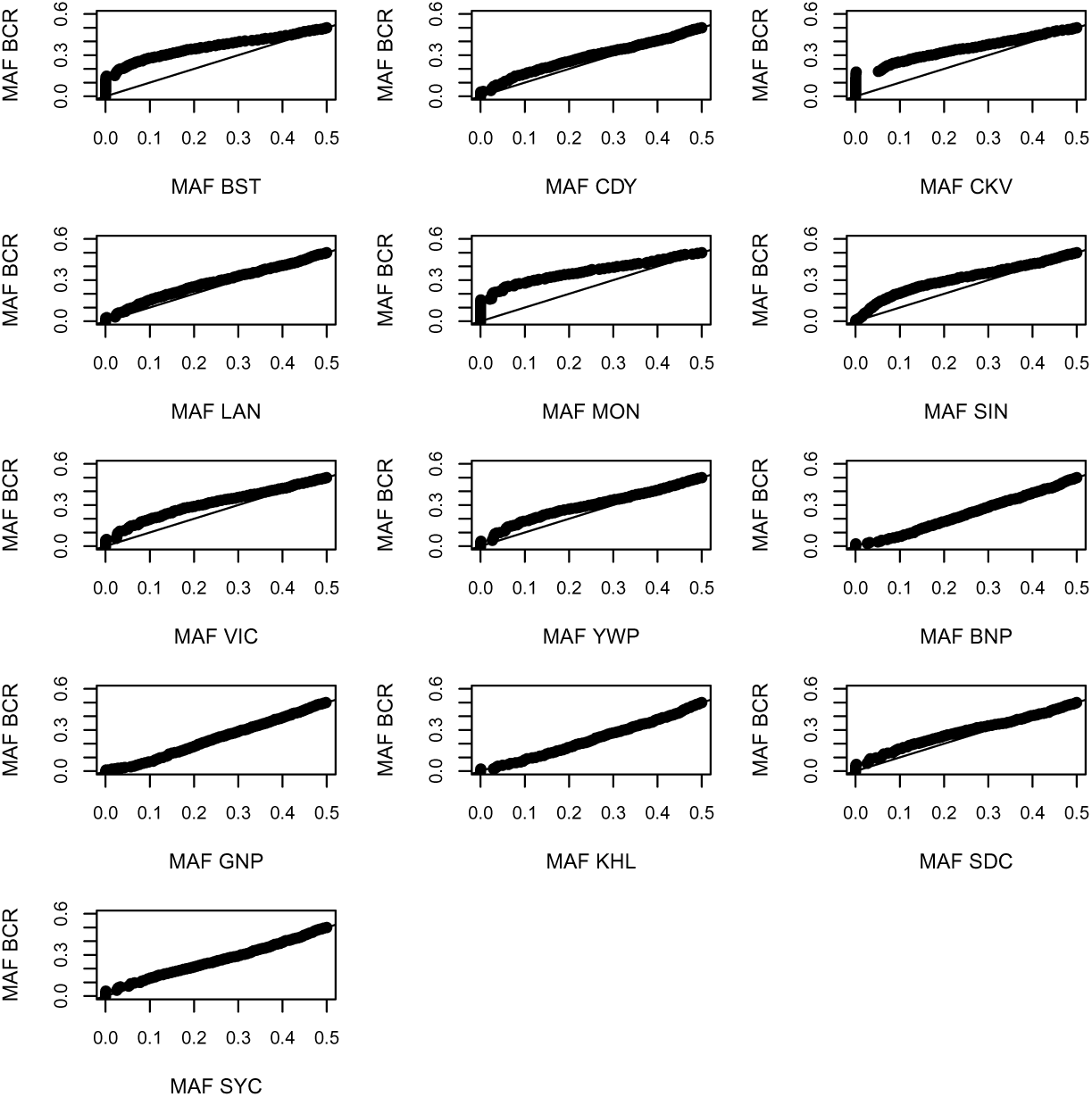
Quantile-quantile plots show the distribution of minor allele frequency (MAF) for all pairs of ancestry informative SNPs (AIMs) for each population when compared to Bull Creek (BCR). Here, AIMs were defined as those SNPs with an allele frequency difference of at least 0.3 between *L. idas* and *L. melissa*. Population abbreviations are defined in Table S1.

**Figure S11:**
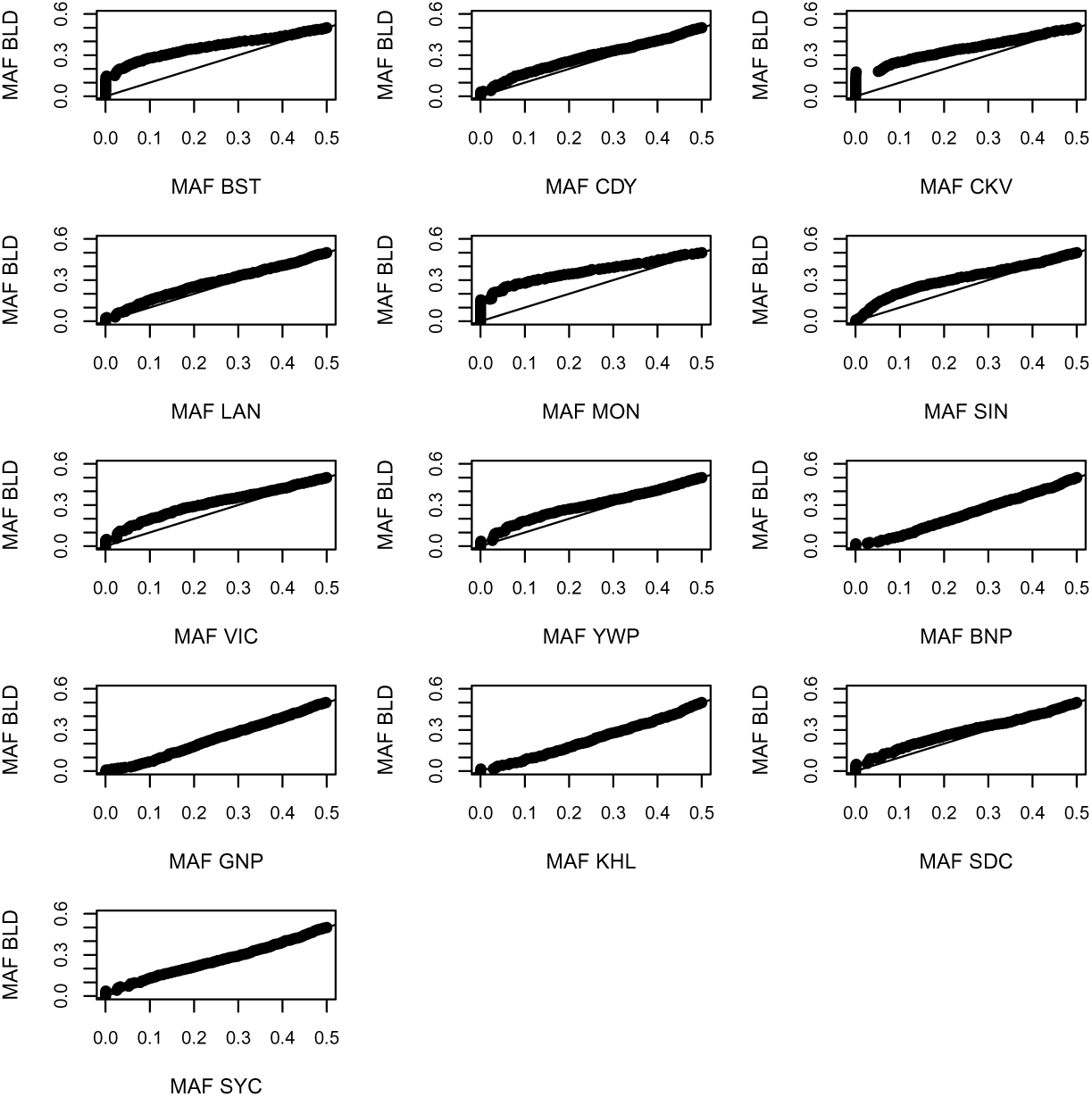
Quantile-quantile plots show the distribution of minor allele frequency (MAF) for all pairs of ancestry informative SNPs (AIMs) for each population when compared to Bald Mountain (BLD). Here, AIMs were defined as those SNPs with an allele frequency difference of at least 0.3 between *L. idas* and *L. melissa*. Population abbreviations are defined in Table S1.

**Figure S12:**
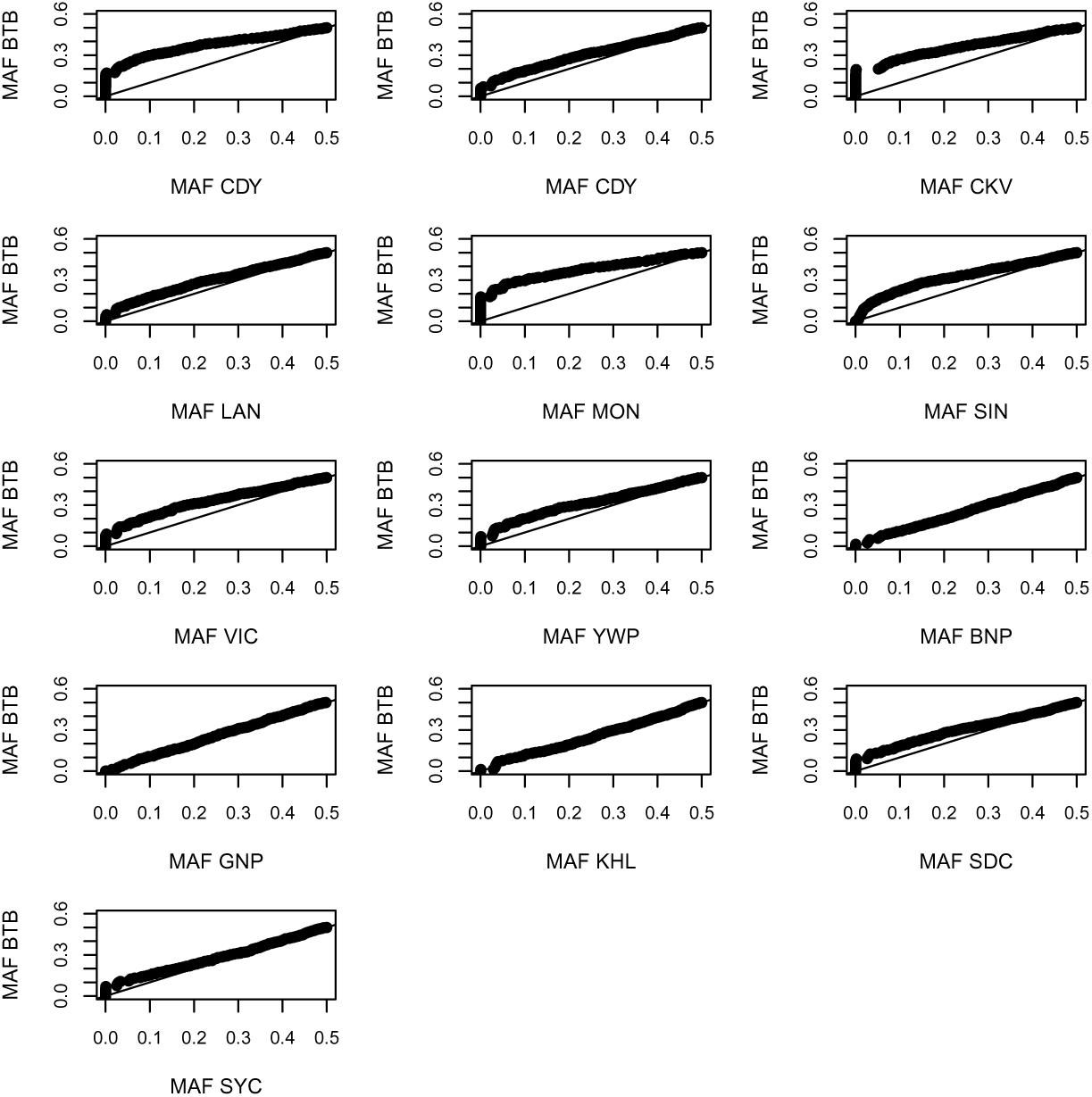
Quantile-quantile plots show the distribution of minor allele frequency (MAF) for all pairs of ancestry informative SNPs (AIMs) for each population when compared to Blacktail Butte (BTB). Here, AIMs were defined as those SNPs with an allele frequency difference of at least 0.3 between *L. idas* and *L. melissa*. Population abbreviations are defined in Table S1.

**Figure S13:**
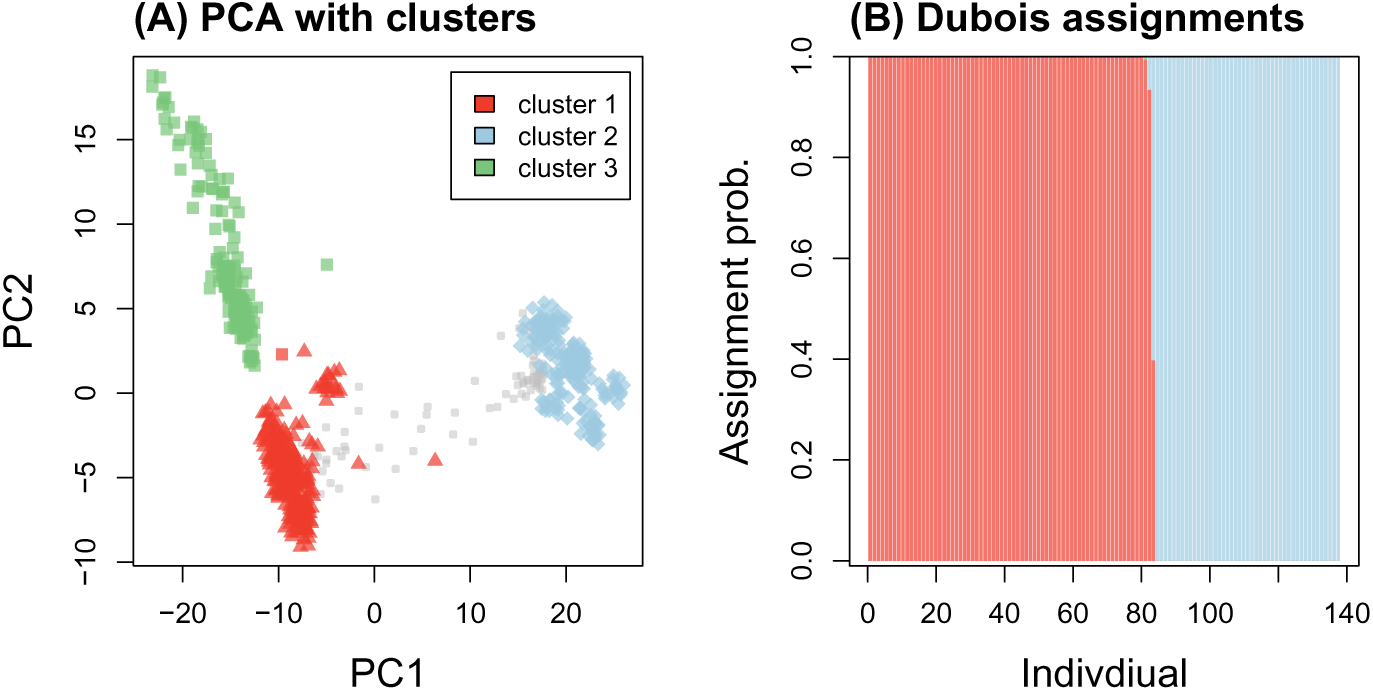
Genetic clusters and genetic assignment of Dubois individuals. Panel (A) shows a results of a PCA for all 835 *Lycaeides*. Points symbols denote nominal taxon (as in Fig. 2): *L. idas* = square, *L. melissa* = diamond, Jackson Hole = triangle. Colors denote cluster assignment from k-means clustering. Dubois individuals (shown in light gray for reference) were not included in the k-means cluster analysis. Panel (B) shows assignment probabilities from discriminant function analysis of the genetic PCs (PC1 and PC2) for the Dubois individuals to each of the three genetic clusters from (A). Individuals are ordered based on their assignment probability to cluster 1 (which corresponds with Jackson Hole). No individuals had assignment probabilites *>* 0.01 to *L. idas*.

**Figure S14:**
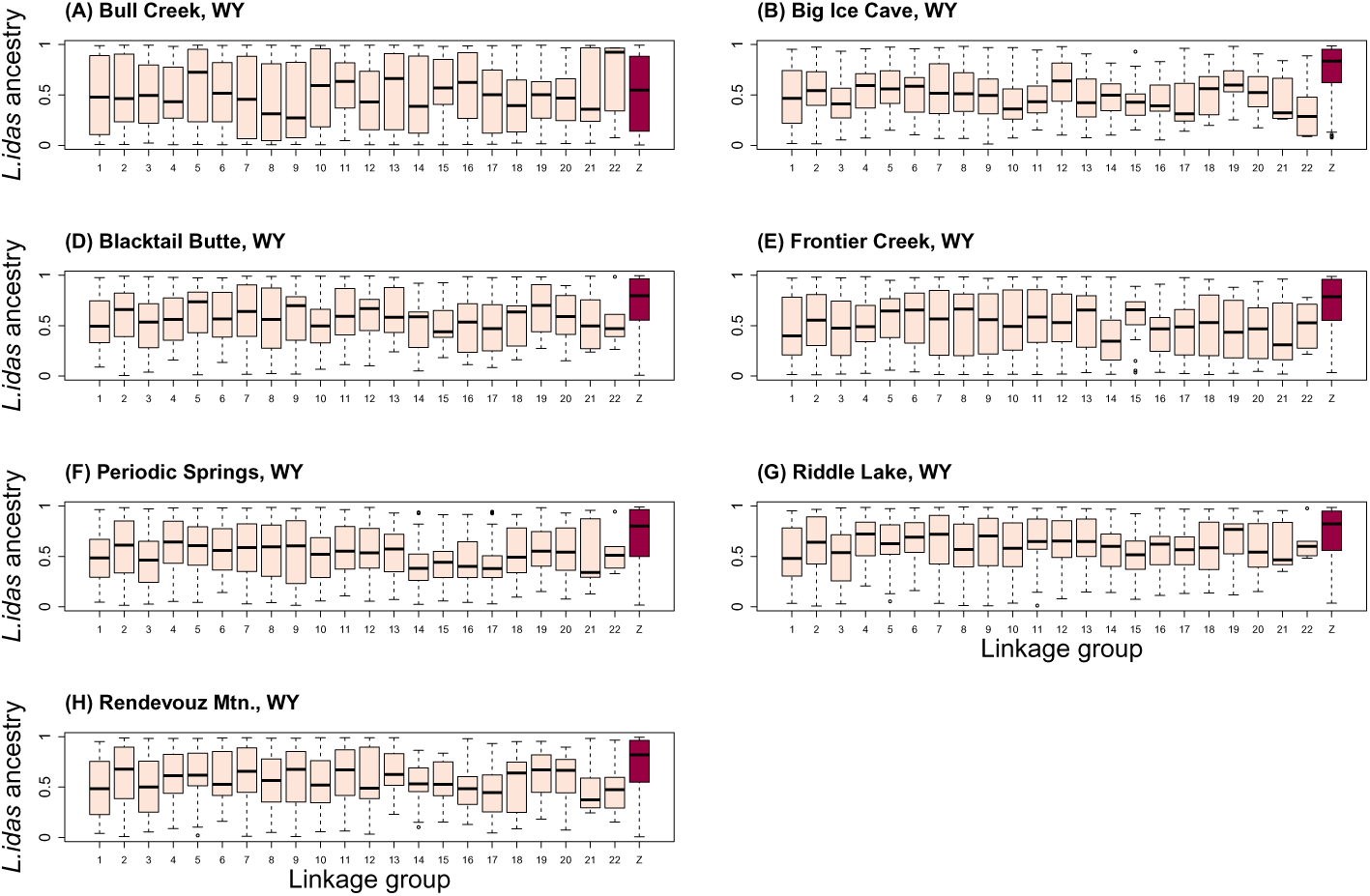
Boxplots show the distribution of *L. idas* ancestry across ancestry informative SNPs (AIMs) for each linkage group in remaining Jackson Hole *Lycaeides* populations. The Z sex chromosome is shown in red.

**Figure S15:**
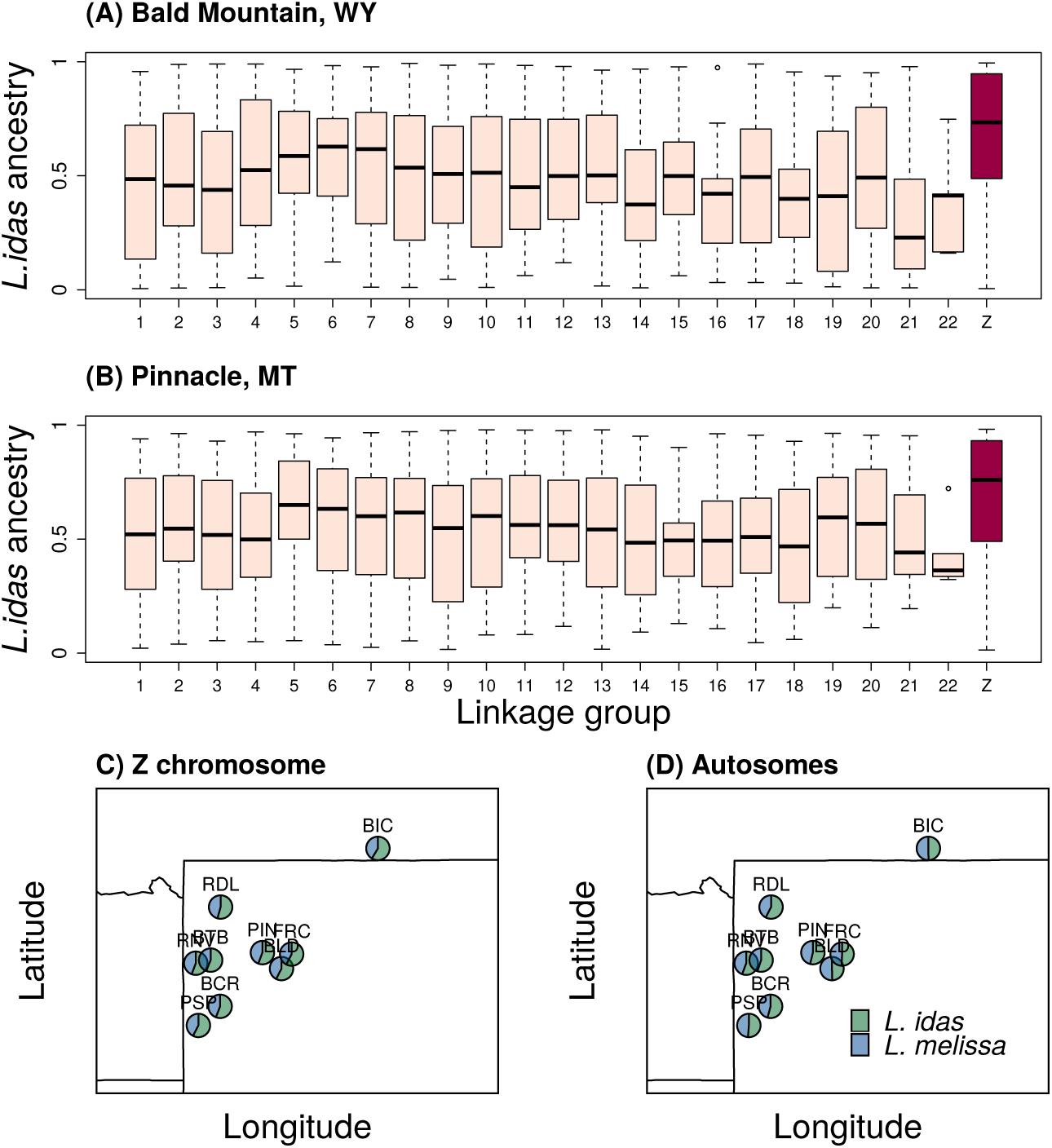
Boxplots show the distribution of *L. idas* ancestry across ancestry informative SNPs (AIMs) for each linkage group in two representative Jackson Hole *Lycaeides* populations–Bald Mountain, WY (BLD) (A) and Pinnacle, WY (PIN) (B). Results are shown for **male butterflies** only. See Fig. S16 for additional populations. The Z sex chromosome is shown in red. Panels (C) and (D) show maps with pie charts reflecting the proportion of *L. idas* and *L. melissa* ancestry (mean) for the Z chromosome (C) and autosomes (D) for each of the nine populations (see Table S1 for population IDs).

**Figure S16:**
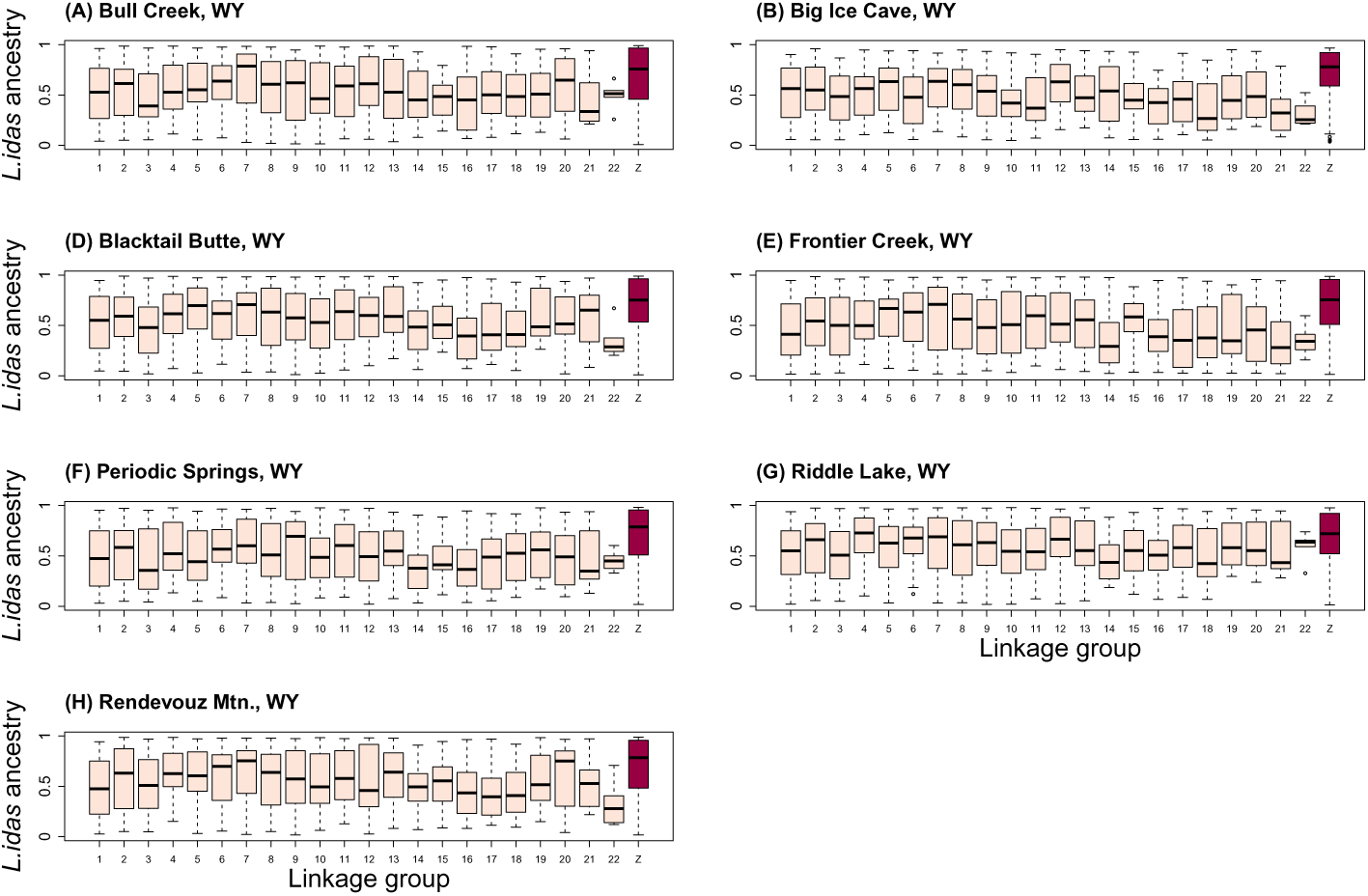
Boxplots show the distribution of *L. idas* ancestry across ancestry informative SNPs (AIMs) for each linkage group in remaining Jackson Hole *Lycaeides* populations. Results are shown for **male butterflies** only. The Z sex chromosome is shown in red.

**Figure S17:**
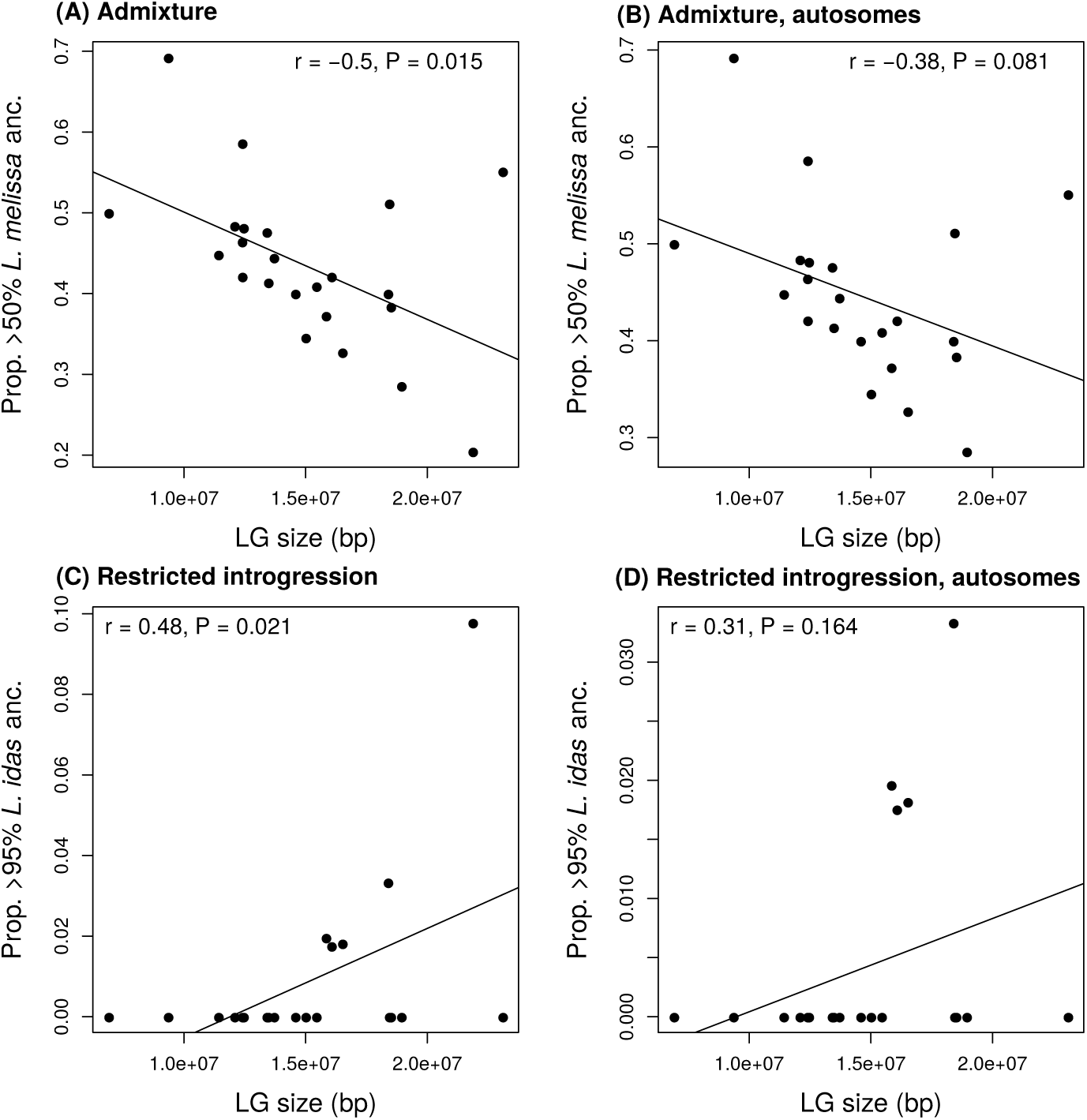
Scatterplots show the relationship between chromosome size and admixture/introgression in Jackson Hole *Lycaeides*. Panels (A) and (B) show the proportion of loci with mean (across populations) ancestry from *L. melissa* > 50% (i.e., where *L. melissa* ancestry is more common than *L. idas* ancestry) for all chromosomes (A) or only the 22 autosomes (B). Panels (C) and (D) show the proportion of loci with mean (across populations) ancestry from *L. idas* > 95% (i.e., nearly fixed for *L. idas* ancestry across all Jackson Hole populations) for all chromosomes (C) or only the 22 autosomes (D). We report the Pearson correlation between ancestry and LG size, and give the best-fit line from a linear regression of ancestry on chromosome size.

**Figure S18:**
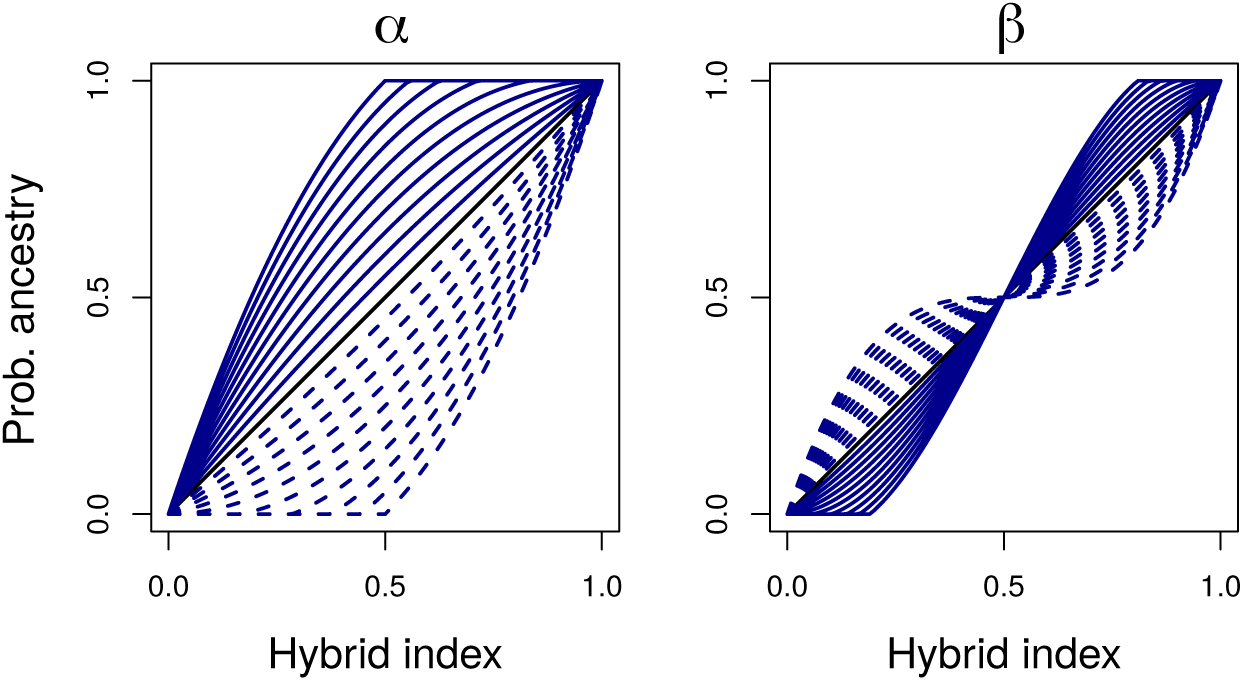
Hypothetical genomic clines. In each panel, the solid black line denotes the probability of ancestry from species one when *α* and *β* are zero, which is equal to an individual’s hybrid index. In the left panel *β* = 0 and *α* varies from 0.25 to 1.25 (solid blue lines) and from −0.25 to −1.25 (dashed blue lines). Similarly, in the right panel *α* = 0 and *β* varies from 0.25 to 1.25 (solid blue lines) and from −0.25 to −1.25 (dashed blue lines).

**Figure S19:**
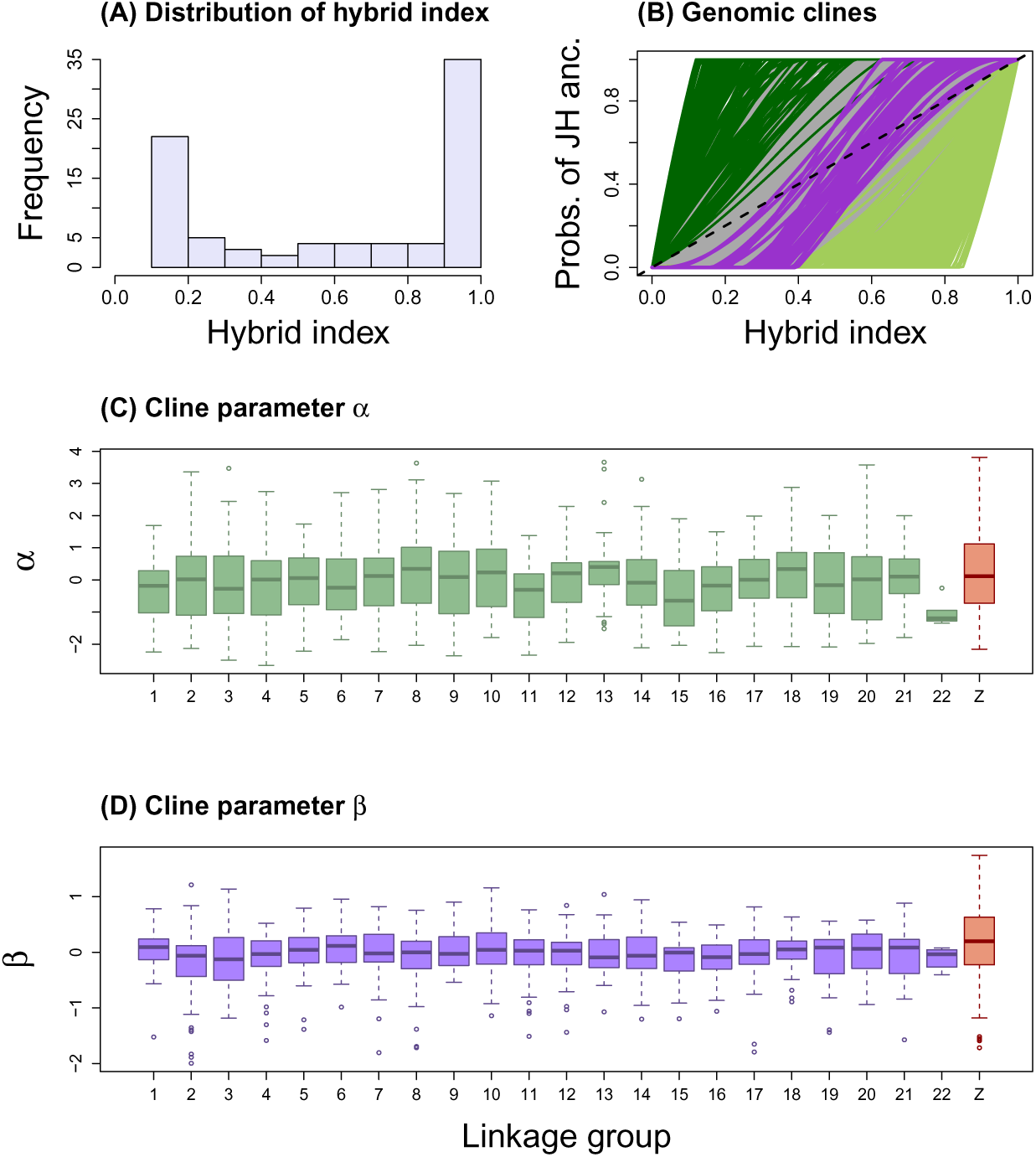
Summary of the genomic clines analysis for **male butterflies**. (A) The histogram depicts the distribution of hybrid indices in the Dubois hybrid zone. (B) This plot shows estimated genomic clines for a subset of ancestry informative SNPs (AIMs). Each solid line gives the estimated probability of Jackson Hole (JH) ancestry for an AIM. Green lines denote cases of credible directional introgression (95% CIs for *α* that exclude zero) and purple lines denote credible cases of restricted introgression (95% CIs for *β* > 0) (gray lines denote clines not credibly different from the genome-average). The dashed line gives the null expectation based on genome-wide admixture. Twenty-seven (*β* > 0), 139 (*α* > 0) and 272 (*α* < 0) AIMs showed credible deviations from null expectations, and for *β* > 0 and *α* > 0 these were over-represented on the Z chromosome (*β* > 0, number on Z = 25, x-fold enrichment = 4.48, *P* < 0.001; *α* > 0, number on Z = 59, x-fold enrichment = 2.06, *P* < 0.001). Boxplots show the distribution of cline parameters *α* (C) and *β* (D) across loci for each linkage group.

**Figure S20:**
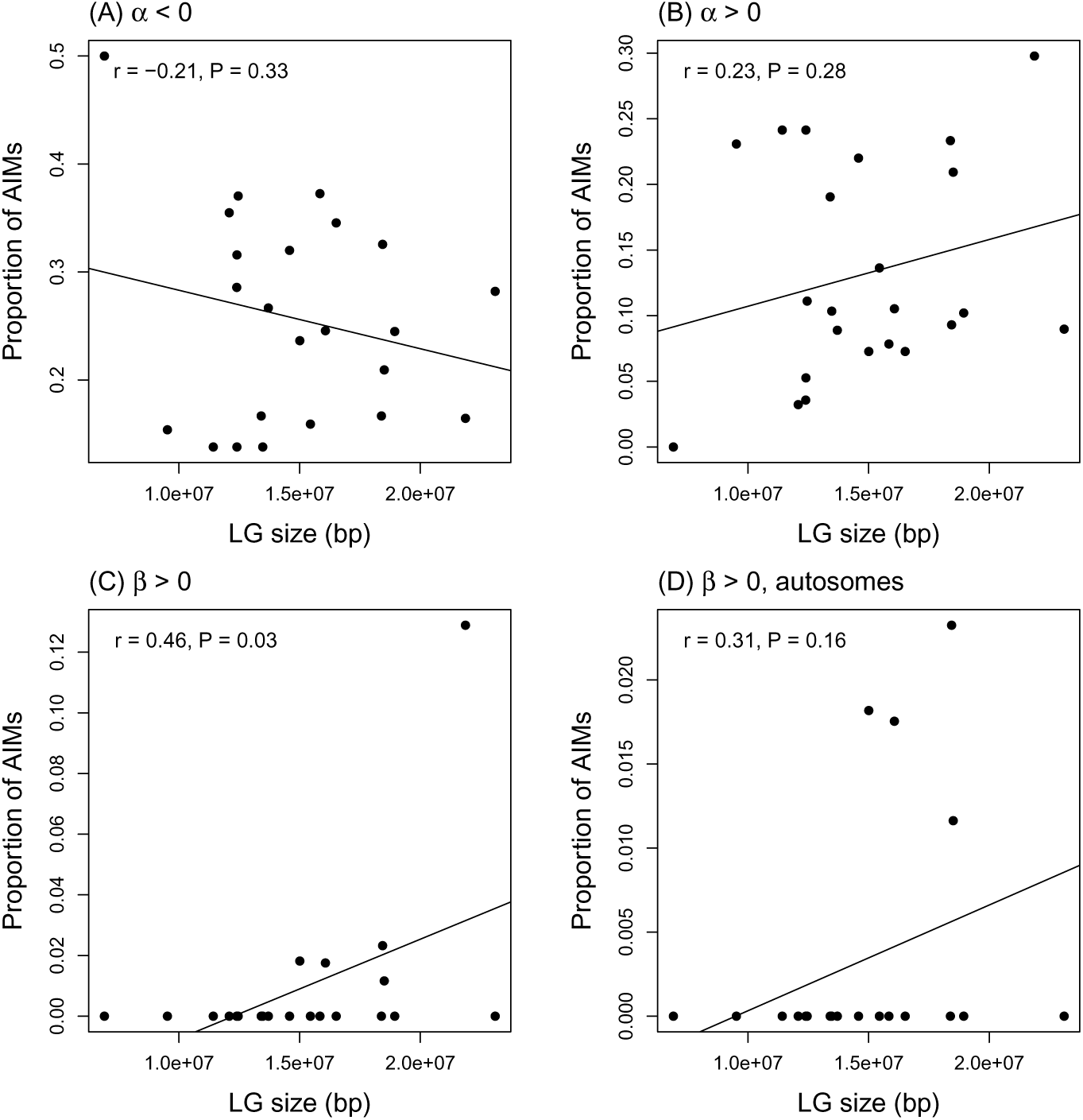
Scatter plots depict the proportion of AIMs on each linkage group (LG) with patterns of introgression that deviate from genome-average expectations as a function of LG size. Panels (A-C) shows all chromosomes, whereas panel (D) considers only the 22 autosomes. Results are shown for *α* > 0 (excess Jackson Hole *Lycaeides* introgression), *α* < 0 (excess *L. melissa* introgression), and *β* > 0 (restricted introgression). The latter is shown with and without the Z chromosome. We report the Pearson correlation between the proportion of AIMs and LG size and associated *P* value, and give the best-fit line from a linear regression.

**Figure S21:**
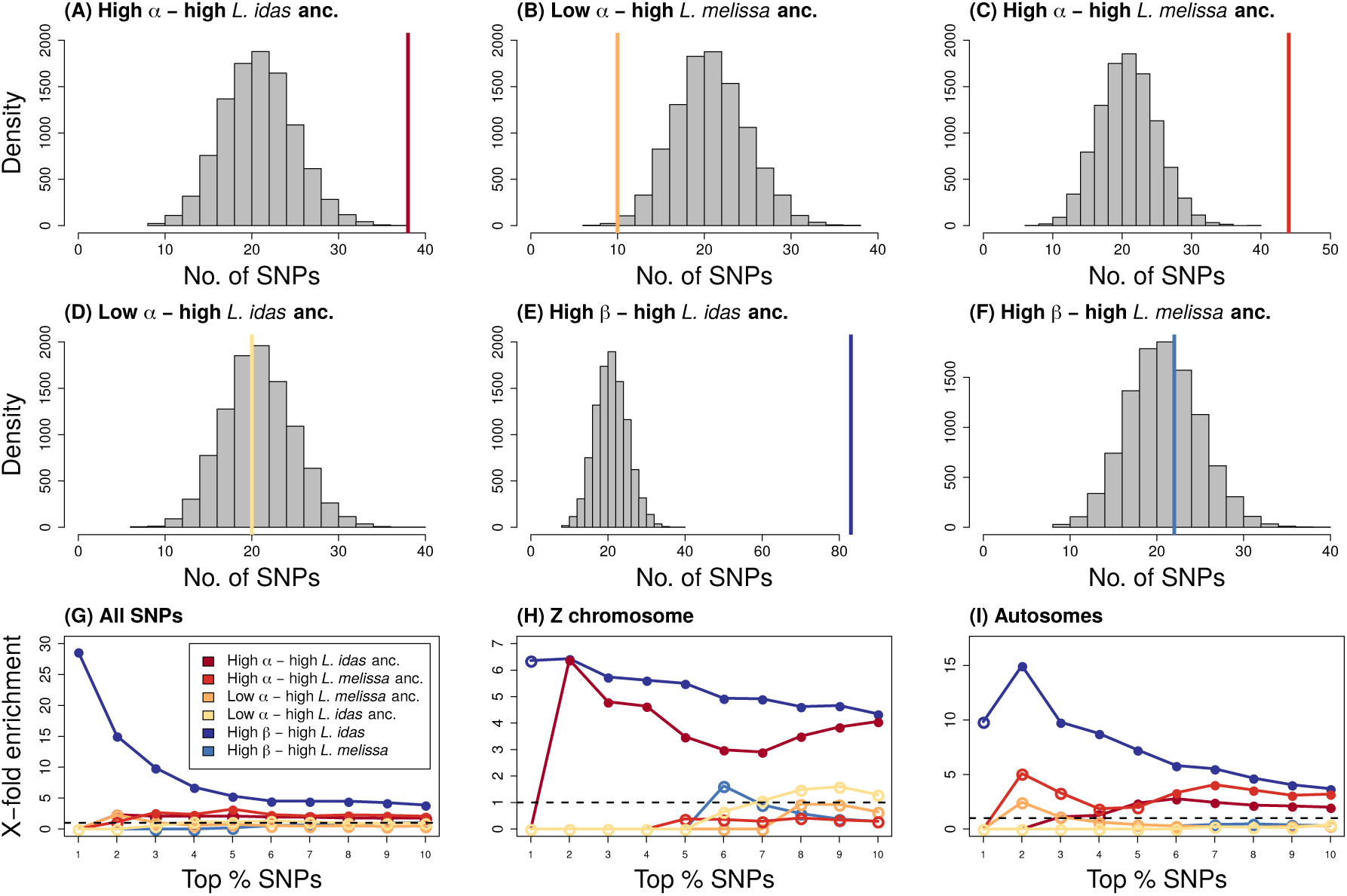
Expected and observed numbers of SNPs with exceptional patterns of introgression in the Dubois hybrid zone and extreme ancestry frequencies in Jackson Hole *Lycaeides*. Here, results are shown for the **2126 SNPs with allele frequency differences of 0.2** or more between *L. idas* and *L. melissa*. Panels A-F show results when considering the top 10% of AIMs in each category. Histograms give null expectations from randomization tests and vertical solid lines show the observed number of AIMs exhibiting a given pattern. Comparisons shown are: directional introgression of Jackson Hole alleles (high *α*) and high *L. idas* ancestry (in Jackson Hole) (A), directional introgression of *L. melissa* alleles (low *α*) and high *L. melissa* ancestry (B), directional introgression of Jackson Hole alleles (high *α*) and high *L. melissa* ancestry (C), directional introgression of *L. melissa* alleles (low *α*) and high *L. idas* ancestry (D), restricted introgression (high *β*) and high *L. idas* ancestry (E), and restricted introgression (high *β*) and high *L. melissa* ancestry (F). Panels G-I show how these results are affected by considering different levels of stringency (i.e., by examining the most extreme 10% to the top 1% of AIMs with each pattern), and when considering only the Z chromosome (G) or only the autosomes (H). Here, circles denote the ratio of the observed to expected overlap from the null, and the circles are filled (*p* ≤ 0.05) or not (*p* > 0.05) to denote whether the overlap is greater than expected by chance.

**Figure S22:**
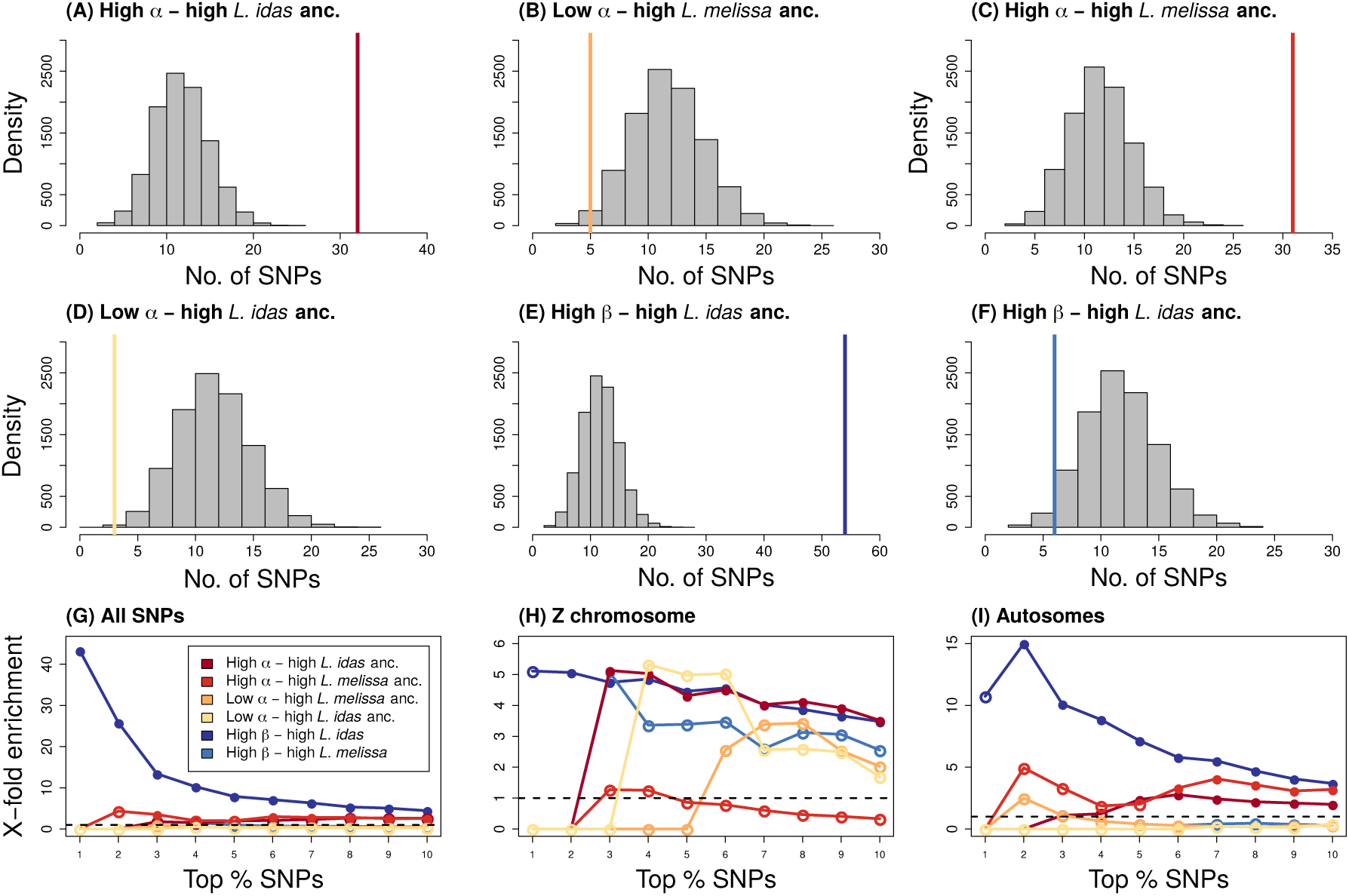
Expected and observed numbers of SNPs with exceptional patterns of introgression in the Dubois hybrid zone and extreme ancestry frequencies in Jackson Hole *Lycaeides*. Here, results are shown for **male butterflies only** and for the **1164 SNPs with allele frequency differences of 0.3** or more between *L. idas* and *L. melissa*. Panels A-F show results when considering the top 10% of AIMs in each category. Histograms give null expectations from randomization tests and vertical solid lines show the observed number of AIMs exhibiting a given pattern. Comparisons shown are: directional introgression of Jackson Hole alleles (high *α*) and high *L. idas* ancestry (in Jackson Hole) (A), directional introgression of *L. melissa* alleles (low *α*) and high *L. melissa* ancestry (B), directional introgression of Jackson Hole alleles (high *α*) and high *L. melissa* ancestry (C), directional introgression of *L. melissa* alleles (low *α*) and high *L. idas* ancestry (D), restricted introgression (high *β*) and high *L. idas* ancestry (E), and restricted introgression (high *β*) and high *L. melissa* ancestry (F). Panels G-I show how these results are affected by considering different levels of stringency (i.e., by examining the most extreme 10% to the top 1% of AIMs with each pattern), and when considering only the Z chromosome (G) or only the autosomes (H). Here, circles denote the ratio of the observed to expected overlap from the null, and the circles are filled (*p* ≤ 0.05) or not (*p* > 0.05) to denote whether the overlap is greater than expected by chance.

**Figure S23:**
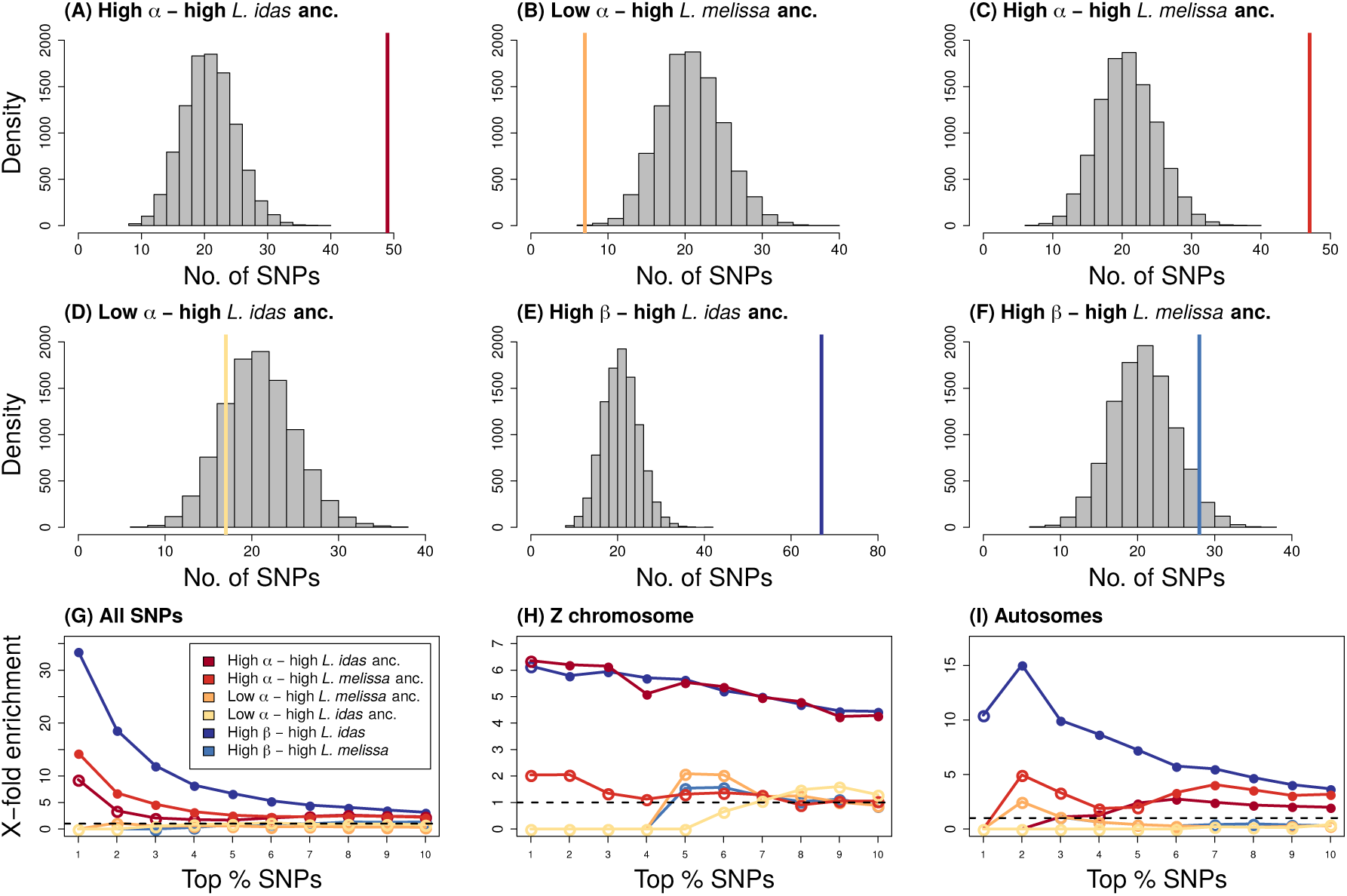
Expected and observed numbers of SNPs with exceptional patterns of introgression in the Dubois hybrid zone and extreme ancestry frequencies in Jackson Hole *Lycaeides*. Here, results are shown for **male butterflies only** and for the **2126 SNPs with allele frequency differences of 0.3** or more between *L. idas* and *L. melissa*. Panels A-F show results when considering the top 10% of AIMs in each category. Histograms give null expectations from randomization tests and vertical solid lines show the observed number of AIMs exhibiting a given pattern. Comparisons shown are: directional introgression of Jackson Hole alleles (high *α*) and high *L. idas* ancestry (in Jackson Hole) (A), directional introgression of *L. melissa* alleles (low *α*) and high *L. melissa* ancestry (B), directional introgression of Jackson Hole alleles (high *α*) and high *L. melissa* ancestry (C), directional introgression of *L. melissa* alleles (low *α*) and high *L. idas* ancestry (D), restricted introgression (high *β*) and high *L. idas* ancestry (E), and restricted introgression (high *β*) and high *L. melissa* ancestry (F). Panels G-I show how these results are affected by considering different levels of stringency (i.e., by examining the most extreme 10% to the top 1% of AIMs with each pattern), and when considering only the Z chromosome (G) or only the autosomes (H). Here, circles denote the ratio of the observed to expected overlap from the null, and the circles are filled (*p* ≤ 0.05) or not (*p* > 0.05) to denote whether the overlap is greater than expected by chance.

**Figure S24:**
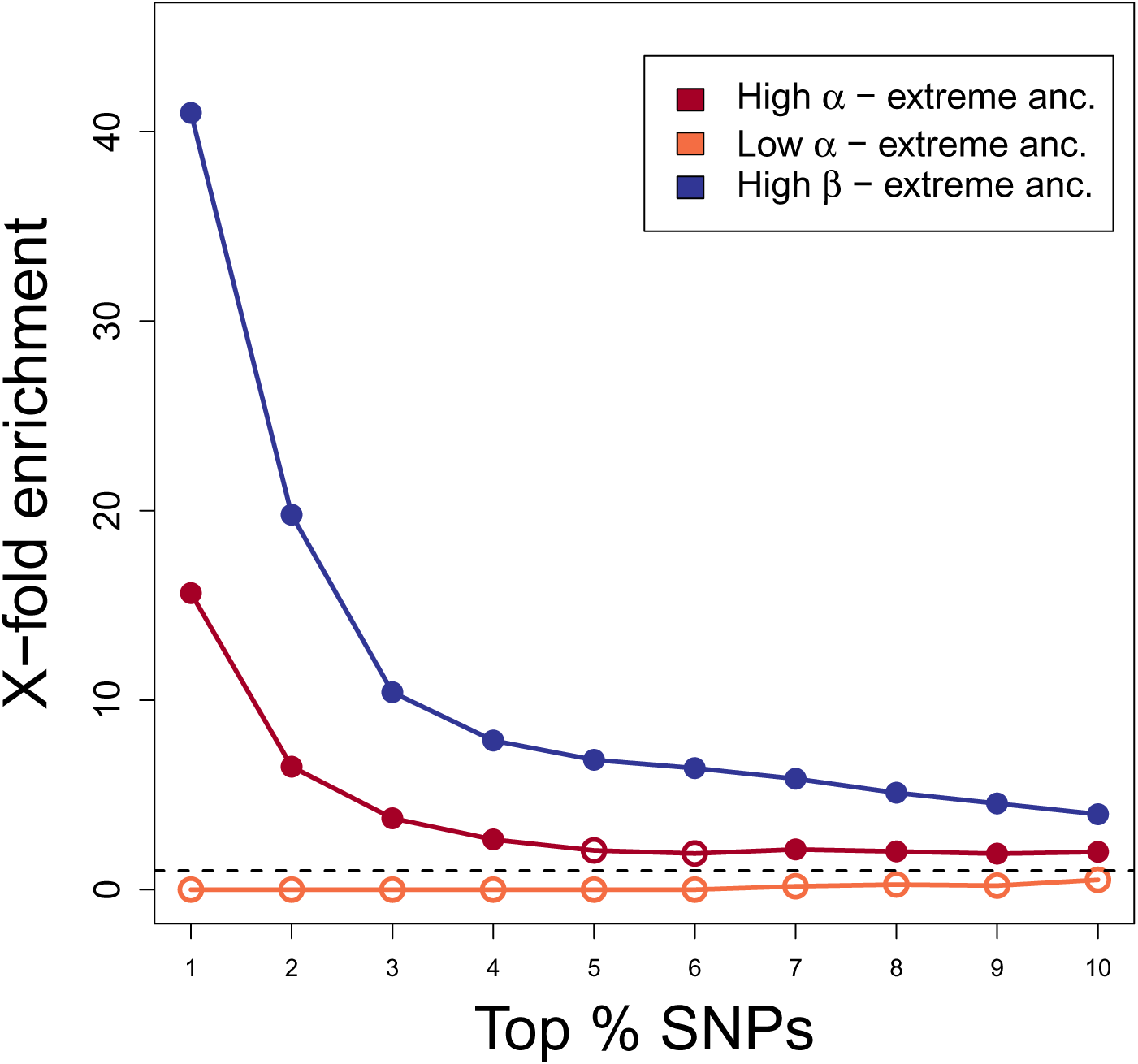
Expected and observed numbers of SNPs with exceptional patterns of introgression in the Dubois hybrid zone and extreme ancestry frequencies in Jackson Hole *Lycaeides*. Comparisons shown are: directional introgression of Jackson Hole alleles (high *α*) and high *L. idas* or *L. melissa* ancestry (i.e., ancestry frequencies closest to 0 or 1), directional introgression of *L. melissa* alleles (low *α*) and high *L. idas* or *L. melissa* ancestry, and restricted introgression (high *β*) and high *L. idas* or *L. melissa* ancestry. Line plot shows comparisons of how these results are affected by considering different levels of stringency (i.e., by examining the most extreme 10% to the top 1% of AIMs with each pattern), and when considering all chromosomes. Here, circles denote the ratio of the observed to expected overlap from the null, and the circles are filled (*p* ≤ 0.05) or not (*p* > 0.05) to denote whether the overlap is greater than expected by chance.

**Figure S25:**
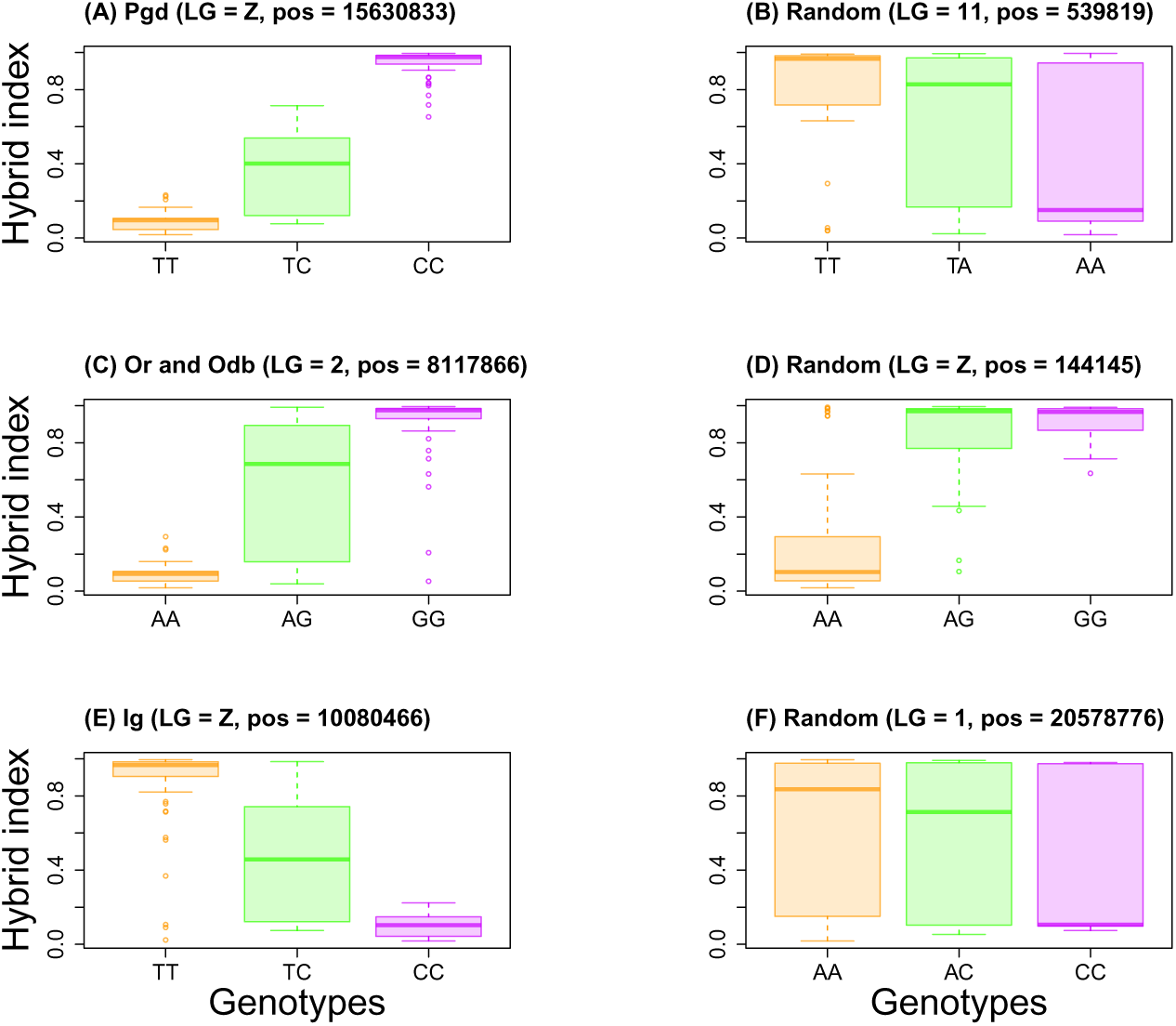
Boxplots shows distribution of hybrid index at each of six AIMs. This includes three of our candidate barrier loci–Pgd = phospogluconate dehydrogenase activity, Or = olfactory receptor activity, Odb = Odorant binding protein, and Ig = Immunoglobulin–and three randomly selected SNPs.

**Figure S26:**
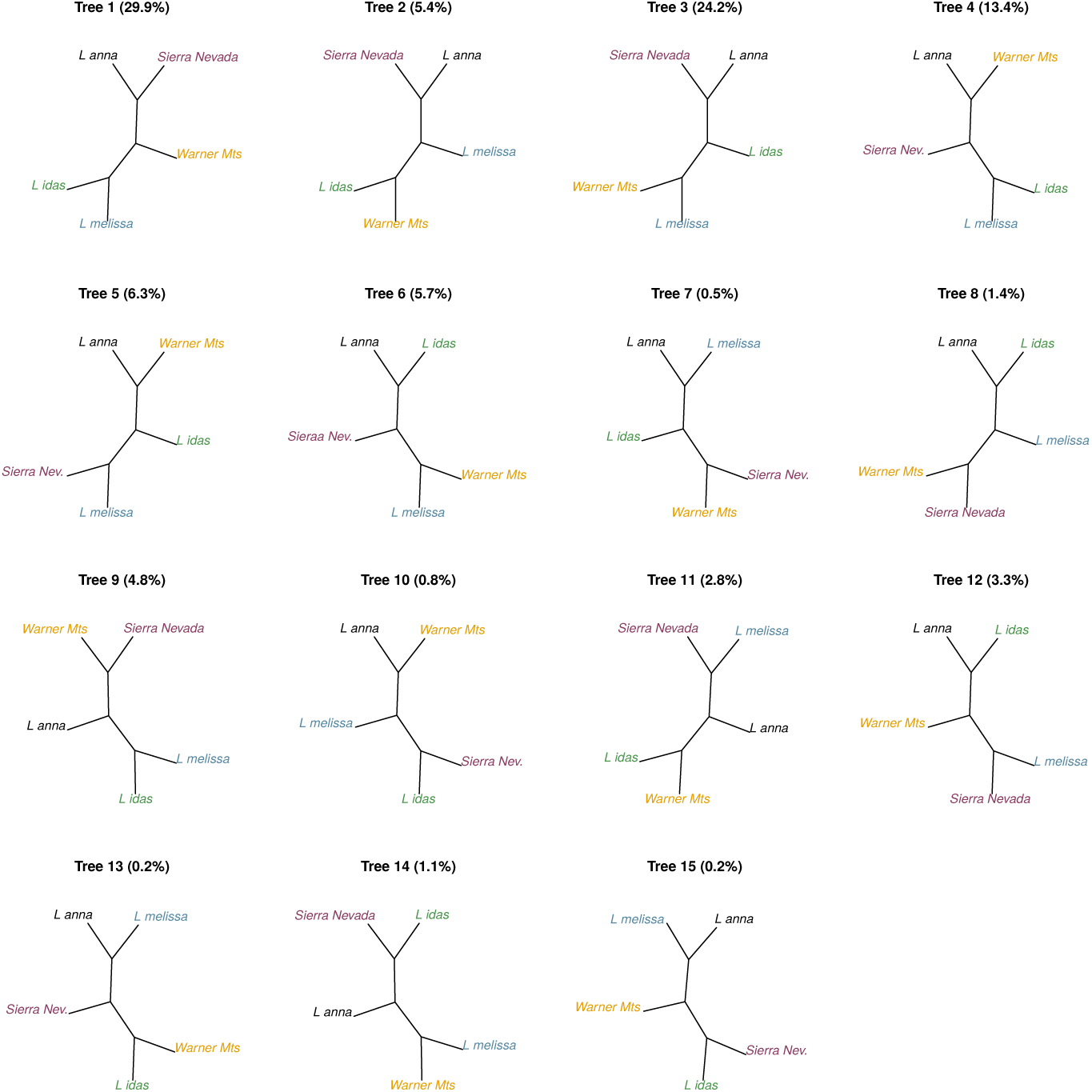
Plot of all 15 observed, unrooted tree topologies. Unrooted maximum likelihood trees were inferred from non-overlapping 1000 SNP windows designated on each linkage group. Number in parentheses give the percent of trees that matched each topology.

**Figure S27:**
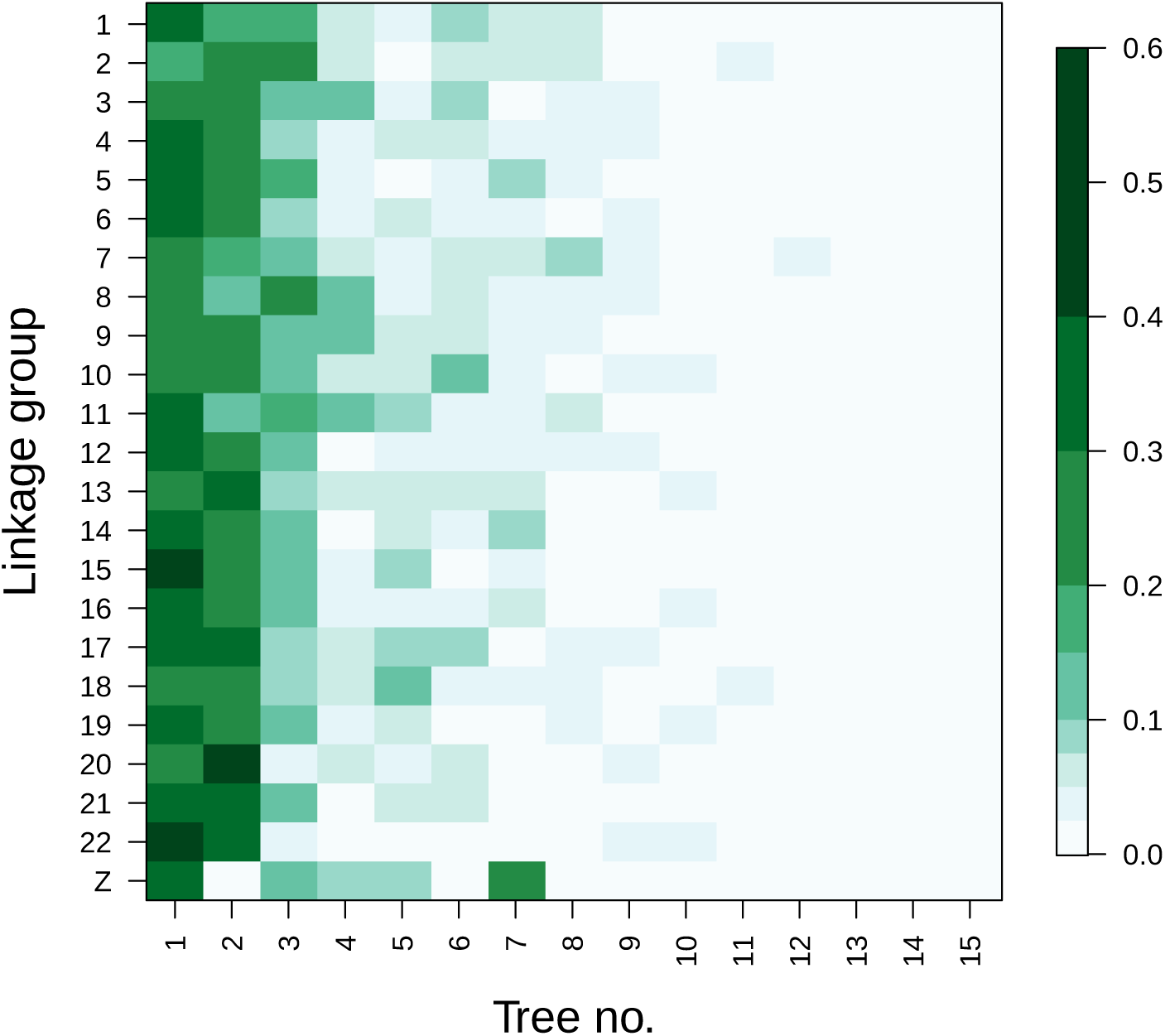
This heatmap shows the proportion of unrooted trees matching each of 15 tree topologies for each linkage group. Unrooted maximum likelihood trees were inferred from non-overlapping 1000 SNP windows designated on each linkage group. Trees 1, 2 and 7 correspond to the topologies shown in Fig. 6 panels (C), (D) and (E), respectively.

**Figure S28:**
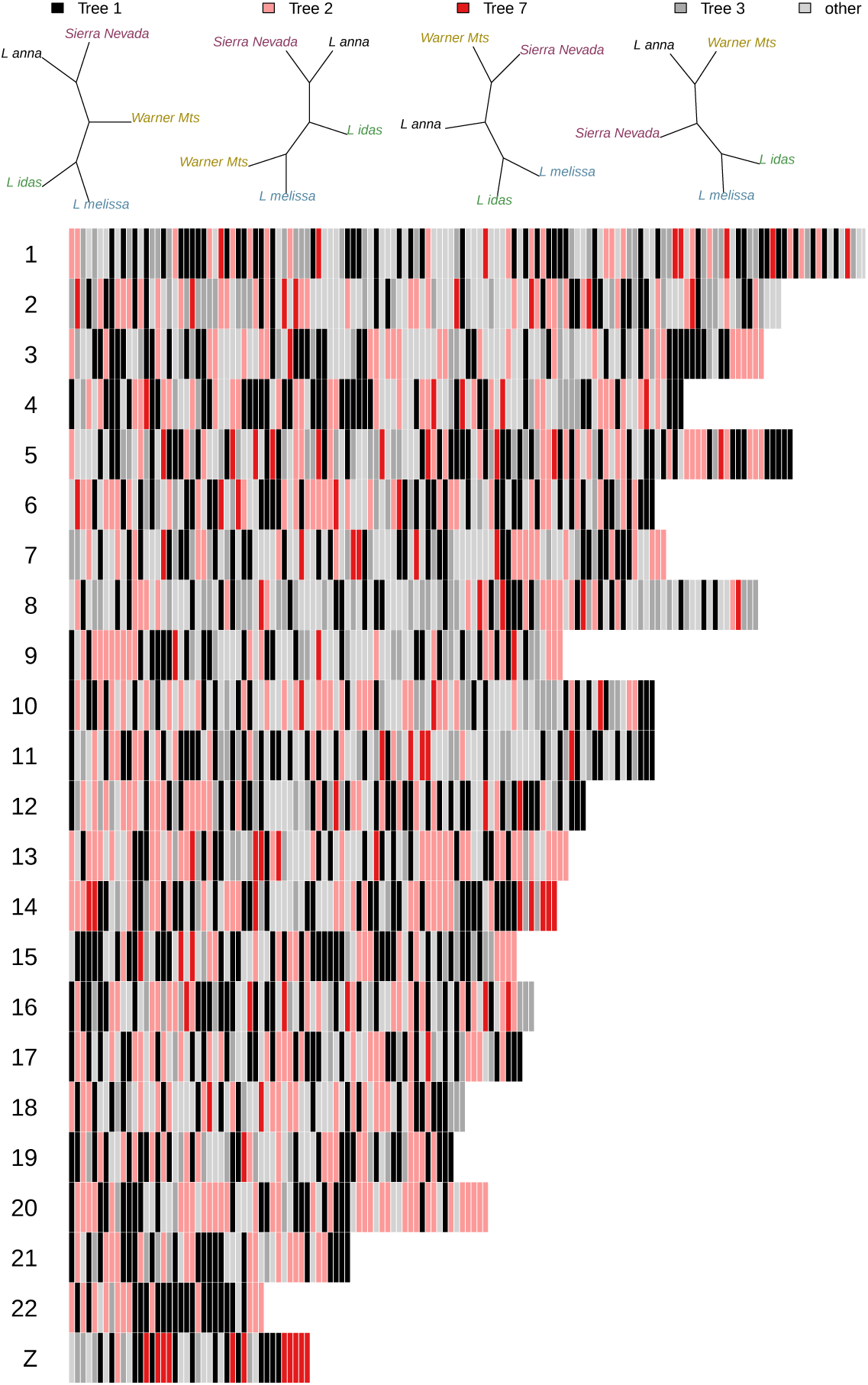
Colored bars denote tree topologies as they vary along linkage groups. Unrooted maximum likelihood trees were inferred from non-overlapping 1000 SNP windows designated on each linkage group. Trees 1, 2 and 7 correspond to the topologies shown in Fig. 6 panels (C), (D) and (E), respectively. Colored bars denote the topology for each 1000 SNP window and these are ordered according physical position along linkage groups. Light gray denotes all topologies except for the trees shown in the legend. Tree topologies exhibit autocorrelations along chromosomes. For example, autocorrelations for tree topology 7 (the restricted introgression tree) on the Z chromosome are greater than expected by chance at lags of one^4^(^3^*r* = 0.41, *P* = 0.002) and two windows (*r* = 0.43, *P* = 0.001).

**Figure S29:**
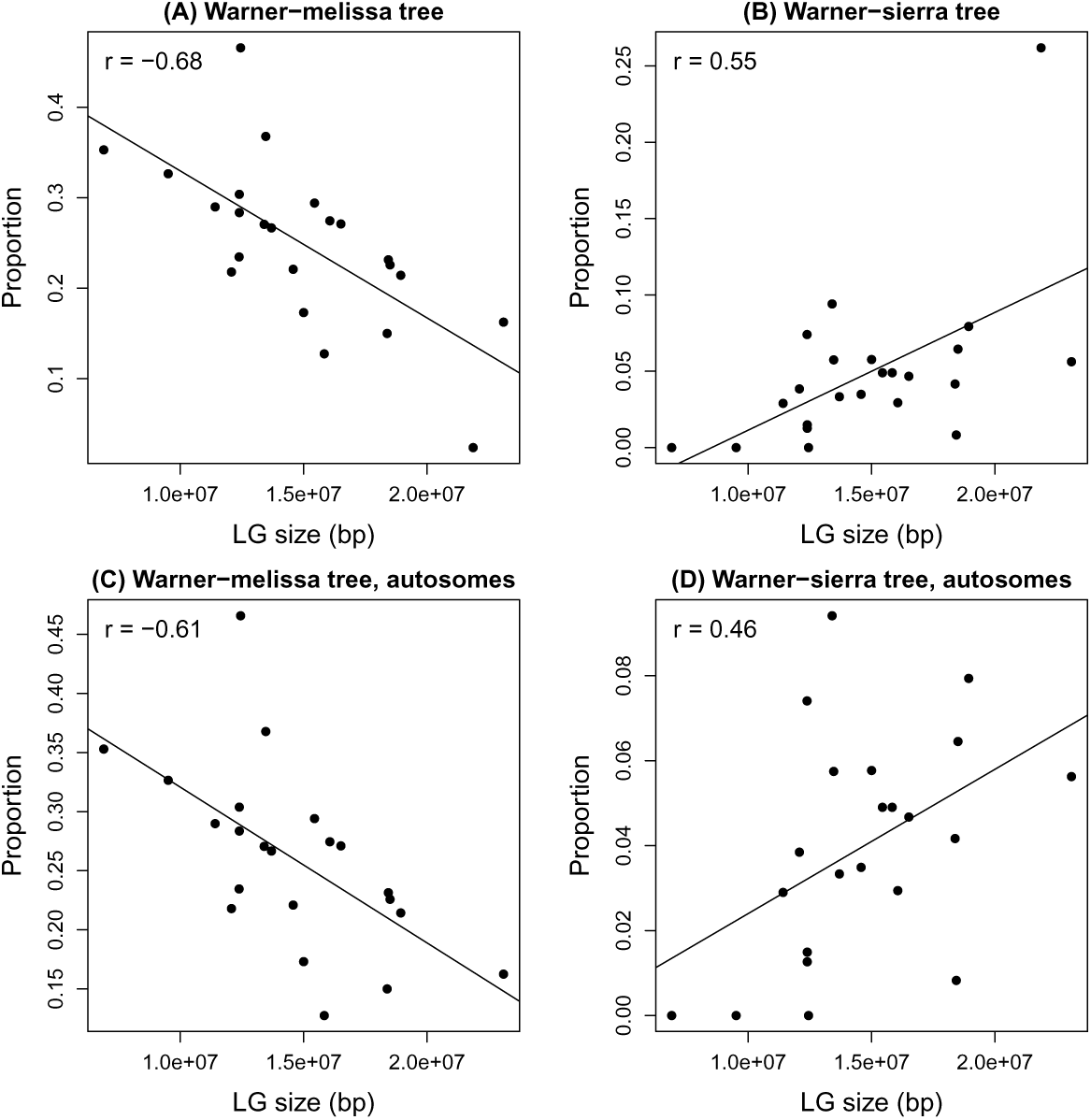
Scatter plots depict the proportion of tree topologies in 1000 SNP windows for each linkage group (LG) as a function of linkage group size in base pairs (bp). Panels (A) and (C) show the proportion of trees with the Warner-*melissa* topology (tree 2), whereas panels (B) and (D) who the proportion of trees with the Warner-sierra topology (tree 7). The top panels show all chromosomes, whereas the bottom panels consider only the 22 autosomes (i.e., these exclude the Z chromosome). We report the Pearson correlation between the proportion of the tree topology and LG size, and give the best-fit line from a linear regression (in all cases, *P* < 0.05 for the effect of LG size).

**Figure S30:**
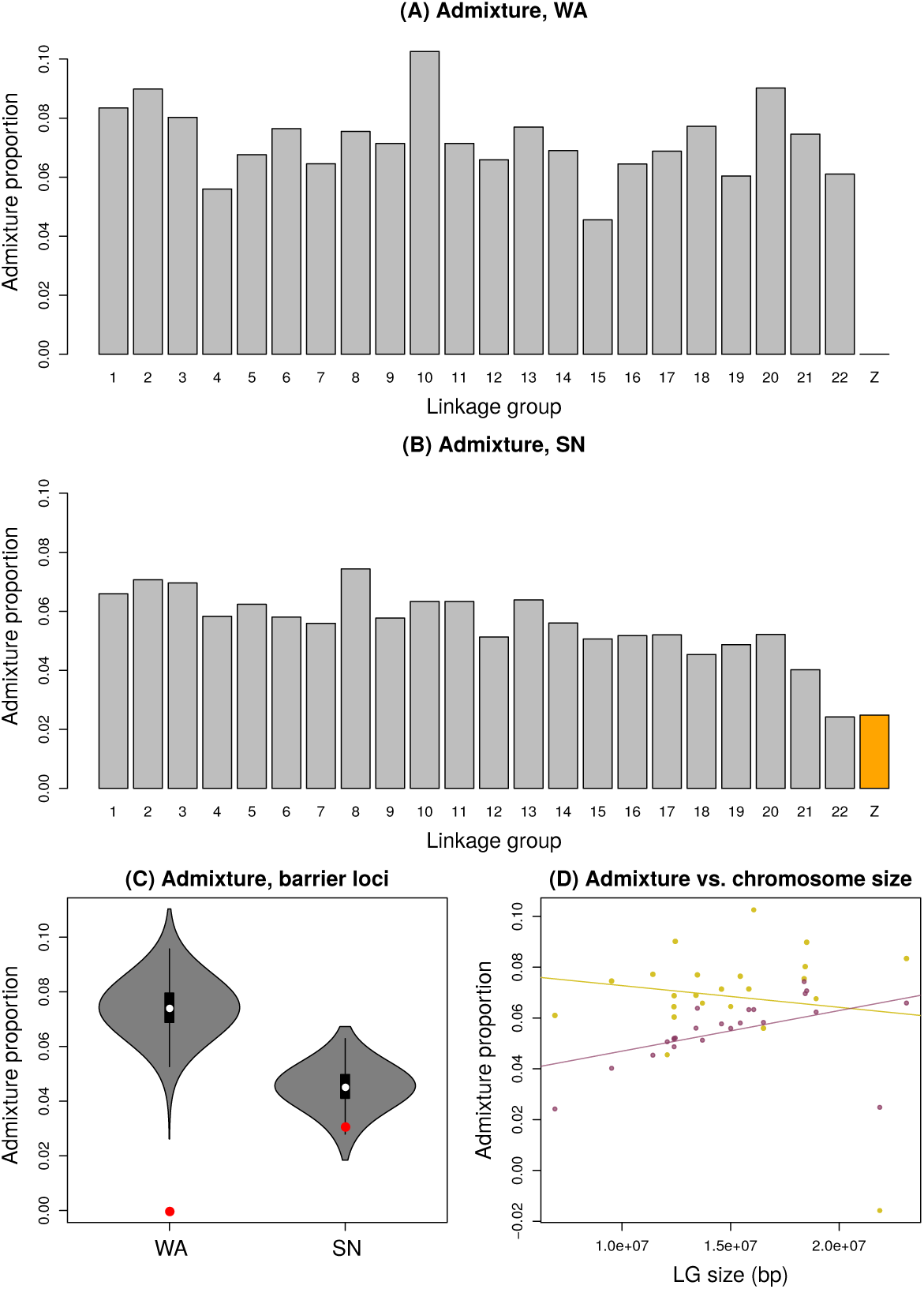
Summary of admixture proportions (*f_d_* estimates) from the phylogenomic analyses. Panels (A) and (B) show admixture proportions for four-taxon trees with the Warner mountains = WA (A) or Sierra Nevada = SN (B) populations by linkage group. Violin plots in panel (C) show the distribution of expected (from a randomization test) admixture proportions for the 51 barrier loci. Red dots show the observed values. Panel (D) shows the relationship between chromosome (LG = linkage group) size and admixture proportion for the Warner mountains (red) (*β* = −8.5*e^−^*^10^, *P* = 0.50, *r*^2^ = 0.02), and Sierra Nevada (gold) (*β* = 1.6*e^−^*^9^, *P* = 0.02, *r*^2^ = 0.23). Points show individual values and the lines show the best fit relationship.

